# Regulatory Changes in the Fatty Acid Elongase *eloF* Underlie the Evolution of Sex-specific Pheromone Profiles in *Drosophila prolongata*

**DOI:** 10.1101/2024.10.09.617394

**Authors:** Yige Luo, Ayumi Takau, Jiaxun Li, Tiezheng Fan, Ben R. Hopkins, Yvonne Le, Santiago R. Ramirez, Takashi Matsuo, Artyom Kopp

## Abstract

Pheromones play a key role in regulating sexual behavior throughout the animal kingdom. In *Drosophila* and other insects, many cuticular hydrocarbons (CHCs) are sexually dimorphic, and some are known to perform pheromonal functions. However, the genetic control of sex-specific CHC production is not understood outside of the model species *D. melanogaster*. A recent evolutionary change is found in *D. prolongata*, which, compared to its closest relatives, shows greatly increased sexual dimorphism in both CHCs and the chemosensory system responsible for their perception. A key transition involves a male-specific increase in the proportion of long-chain CHCs. Perfuming *D. prolongata* females with the male-biased CHCs reduces copulation success, suggesting that these compounds function as sex pheromones. The evolutionary change in CHC profiles correlates with a male-specific increase in the expression of multiple genes involved in CHC biosynthesis, including fatty acid elongases and reductases and other key enzymes. In particular, *elongase F*, which is responsible for producing female-specific pheromones in *D. melanogaster*, is strongly upregulated in *D. prolongata* males compared both to females and to males of the sibling species. Induced mutations in *eloF* reduce the amount of long-chain CHCs, resulting in a partial feminization of pheromone profiles in *D. prolongata* males while having minimal effect in females. Transgenic experiments show that sex-biased expression of *eloF* is caused in part by a putative transposable element insertion in its regulatory region. These results reveal one of the genetic mechanisms responsible for a recent evolutionary change in sexual communication.

## Introduction

Communication, both between and within the sexes, plays a pivotal role in sexual selection and the evolution of sexual dimorphism (Andersson 1994; West-Eberhard 2014; Schaefer and Ruxton 2015; Broder et al. 2021; Buchinger and Li 2023). However, our understanding of the genetic control of both signaling and signal perception remains limited outside traditional model systems. In insects, as well as other animals, pheromones are one of the key methods of communication (Steiger and Stökl 2014; Yew and Chung 2015; Stökl and Steiger 2017; Buchinger and Li 2023). Among the most common insect pheromones are cuticular hydrocarbons (CHCs), which affect a wide range of social and non-social functions including maintaining water balance (Chung et al. 2014; Chung and Carroll 2015; Wang et al. 2022), resource acquisition (Bartelt et al. 1985), social aggregation (Suzuki 1980), cohort recognition (Thome 1982), mate choice (Smadja and Butlin 2008; Laturney and Moehring 2012; Chung et al. 2014), aggression (Wang and Anderson 2010; Zwarts et al. 2012), and signaling fecundity (Monnin 2006) and immunocompetence (Lawniczak et al. 2007).

Much of our understanding of pheromone communication comes from *Drosophila,* where several chemicals have been confirmed to have pheromonal effects (Ferveur 2005; Bontonou and Wicker-Thomas 2014; Yew and Chung 2015; Khallaf et al. 2021). In *D. melanogaster*, the male-specific compounds cis-Vaccenyl Acetate (cVA) and 7-Tricosene (7T) promote aggression when perceived by males and increase receptivity when perceived by females (Grillet et al. 2006; Kurtovic et al. 2007; Datta et al. 2008; Ruta et al. 2010; Wang et al. 2011). In contrast, the female-specific 7,11-heptacosadiene (7,11-HD) functions as an aphrodisiac (Ferveur and Sureau 1996). 7,11-HD initiates a neural cascade that flows from peripheral chemoreceptors to the central nervous system to stimulate male courtship behavior, whereas the perception of 7T inhibits the courtship circuitry in males and regulates reproductive functions in females (Bray and Amrein 2003; Toda et al. 2012; Seeholzer et al. 2018).

The importance of CHCs in mating behavior can contribute to the evolution of reproductive barriers (Fan et al. 2013; Combs et al. 2018). The best-studied example is found in *D. melanogaster* and its sibling species *D. simulans*, where interspecific differences in the processing of the 7T and 7,11-HD signals contribute to pre-mating isolation (Fan et al. 2013; Combs et al. 2018). Within *D. melanogaster*, a higher abundance of female-specific 5,9-heptacosadiene in African populations contributes to the partial isolation between African and non-African strains (Wu et al. 1995; Ferveur et al. 1996; Fang et al. 2002; Grillet et al. 2012). Divergent pheromone profiles also contribute to reproductive isolation in other *Drosophila* species, including the 9-pentacosene between different populations of *D. elegans* (Ishii et al. 2001), 2-methyl hexacosane between *D. serrata* and *D. birchii* (Howard et al. 2003; Chung et al. 2014), and 10-heptadecen-2-yl acetate between different subspecies of *D. mojavensis* (Khallaf et al. 2020).

Understanding the genetic basis of pheromone evolution has been facilitated by a well-characterized pathway for CHC biosynthesis. In insects, key steps in this process, including fatty acid synthesis, desaturation, elongation, and decarboxylation, are highly conserved (Blomquist and Bagnères 2010; Wicker-Thomas et al. 2015). These reactions take place mainly in adult oenocytes, a specialized cell type located beneath the abdominal epidermis (Ferveur et al. 1997; Billeter et al. 2009; Makki et al. 2014). Dietary lipids, such as palmitic and stearic acids, are CoA-activated by fatty acyl synthases, followed by the introduction of position-specific double bonds catalyzed by desaturases. Elongation proceeds with the incorporation of malonyl-CoA, adding two carbons at a time to the growing precursor chain. The synthesis of very-long-chain CHCs is catalyzed by fatty acid elongases (FAEs), with the additional involvement of three other categories of enzymes: 3-keto-acyl-CoA-reductase (KAR), 3-hydroxy-acyl-CoA dehydratase (HACD), and trans-enoyl-CoA-reductase (TER) (Chertemps et al. 2007; Wicker-Thomas et al. 2015; Yew and Chung 2015). Fatty acyl-CoA reductases (FARs) act on the very long chain fatty acyl-CoAs produced by the elongation process, reducing them to aldehydes. From these aldehydes, mature CHCs are produced by oxidative decarboxylation catalyzed by insect-specific cytochrome P450 (Qiu et al. 2012). The multi-stage chemistry that creates the final structure of CHCs offers multiple points at which the end products can be modified. Variation in CHC profiles has been attributed to genes controlling the positions of double bonds (Dallerac et al. 2000; Chertemps et al. 2006), methyl branches (Chung et al. 2014), and chain length (Chertemps et al. 2007; Combs et al. 2018; Pei et al. 2021; Rusuwa et al. 2022).

Sexually dimorphic CHCs have been observed in most *Drosophila* species that have been examined (81/99) (Khallaf et al. 2021), but our understanding of how sex-specific pheromones are produced continues to be based on genetic studies in *D. melanogaster*. A complete feminization of pheromone profiles can be achieved by targeted expression of the female sex determiner, *transformer* (*tra*), in adult male oenocytes (Ferveur et al. 1997). Downstream, at least two key enzymes are under the control of the sex differentiation pathway: *elongase F* (*eloF*) and *desaturase F* (*desatF*, also known as *Fad2*), which control carbon chain elongation and the production of alkadienes, respectively (Chertemps et al. 2006; Chertemps et al. 2007). Female-specific expression of these enzymes contributes to the production of 7,11-HD, the critical female pheromone in *D. melanogaster*, as well as to the higher abundance of very long chain CHCs in females. However, a comparative analysis has shown that the female-restricted expression of *desatF* has evolved relatively recently, in the common ancestor of *D. melanogaster* and *D. erecta,* and that more distantly related *Drosophila* species *express desatF* in a sexually monomorphic manner that correlates with sexually monomorphic diene abundance (Shirangi et al. 2009). The evolution of sex-biased *desatF* expression in the *D. melanogaster* lineage was associated with the gain of binding sites for *doublesex* (*dsx*), the key transcription factor that acts downstream of *tra* to direct the sexual differentiation of somatic cells, in the oenocyte enhancer of *desatF* (Shirangi et al. 2009). Outside of *D. melanogaster* and its closest relatives, the genetic basis of sex-specific pheromone production, and especially the synthesis of male-specific pheromones that are found in many *Drosophila* species (Khallaf et al. 2021), is largely unknown.

In this report, we examine the genetic basis and evolutionary origin of male-biased pheromones in *D. prolongata*. This species exhibits multiple derived sex-specific traits compared to its close relatives (Singh and Gupta 1977), making it an attractive model for investigating coevolution between signals and receptors that mediate sexual communication. Along with many species-specific features of mating behavior and male-male aggression (Setoguchi et al. 2014; Amino and Matsuo 2020; Minekawa et al. 2020), *D. prolongata* has strongly diverged from its relatives both in the chemical signals and in their receptors. On the sensory perception side, this species shows a dramatic, sex-specific increase in the number of gustatory organs on the front legs of males (Luecke et al. 2022). Leg gustatory organs have a well-characterized role in sex-specific pheromone perception in *D. melanogaster* (Bray and Amrein 2003; Toda et al. 2012; Seeholzer et al. 2018), and *D. prolongata* males use their front legs extensively in both courtship and male-male aggression (Setoguchi et al. 2014; Setoguchi et al. 2015; Amino and Matsuo 2020; Minekawa et al. 2020; Yoshimizu et al. 2022), suggesting that this morphological change may have important behavioral consequences. And on the signaling side, *D. prolongata* shows a recently evolved, strongly sex-biased CHC profile (Luo et al. 2019). Specifically, the difference involves the relative amounts of three serial chemical homologs, 9-tricosene (9T), 9-pentacosene (9P), and 9-heptacosene (9H). These molecules differ only in the length of the carbon backbone, and are likely to share common biosynthetic origin. While its closest relatives such as *D. rhopaloa* and *D. carrolli* are sexually monomorphic in the abundance of these CHCs, *D. prolongata* males show a dramatic increase in the amounts of 9P and 9H, and a concomitant reduction in the amount of 9T, compared to females.

To identify the genetic changes responsible for the evolutionary transition from sexually monomorphic to sexually dimorphic CHC profiles, we compared gene expression in pheromone-producing tissues between *D. prolongata* and *D. carrolli*. We find that *D. prolongata* males show increased expression of many enzymes involved in CHC synthesis, including multiple fatty acyl elongases and reductases. We show that *eloF*, which is responsible for the female-biased abundance of long-chain CHCs in *D. melanogaster*, is expressed in a male-specific manner in *D. prolongata,* due in part to changes in its *cis*-regulatory sequences, and is partly responsible for the increased abundance of 9P and 9H in *D. prolongata* males. Finally, we confirm that these CHCs affect sexual behavior. Together, our results reveal one of the genetic mechanisms responsible for a recent evolutionary change in sexual communication.

## Results

### Perfuming with male-specific pheromones reduces female mating success

Male-biased chemical cues are often used in a reproductive context, for example as inhibitory signals against female remating that function as chemical mate-guarding strategy (Ferveur & Sureau, 1996; Jallon et al., 1981; Laturney & Billeter, 2016; Ng et al., 2014). We previously showed that *D. prolongata* exhibits a male-specific increase in the abundance of two long-chain CHCs, 9-pentacosene (9P) and 9-heptacosene (9H) (Luo et al. 2019). To investigate the role of these hydrocarbons in mating behavior, we examined male-female interactions by pairing single virgin males with single virgin females perfumed with synthetic 9P or 9H. On average, each female received ∼350 ng of extra 9P in the 9P treatment and ∼90 ng of additional 9H in the 9H treatment, as shown by GC-MS (Fig. 1 A’, B’). Perfumed females, therefore, had a masculinized pheromone profile with an abundance of male-biased hydrocarbons intermediate between those observed in normal *D. prolongata* males and females (Fig. 1 A, B).

**Figure 1.**
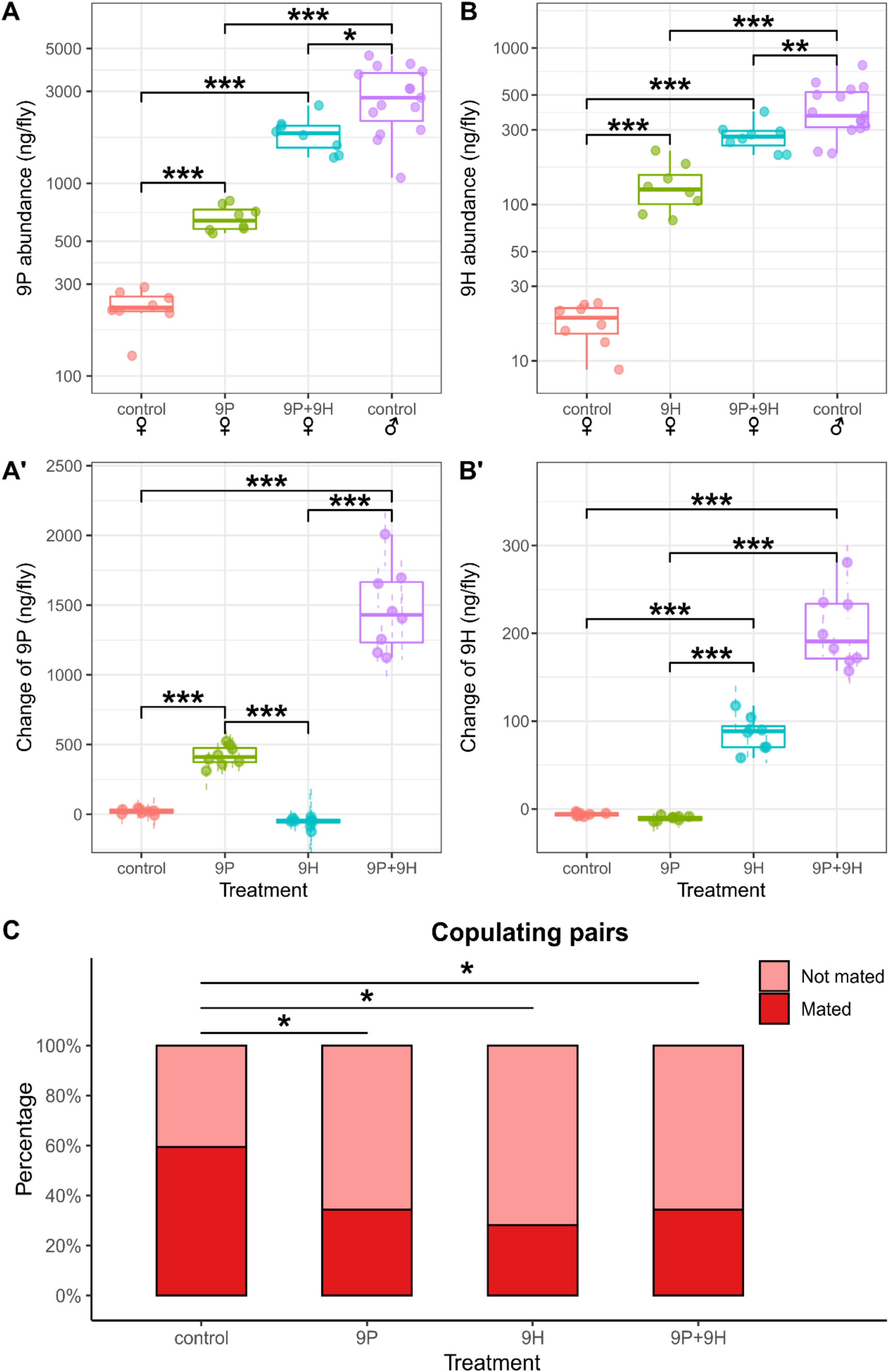
Perfuming male-biased long-chain hydrocarbons on virgin females reduces copulation success in *D. prolongata*. (A, A’) Boxplots showing the total abundance of 9P (A) and change in the abundance of 9P (A’) in ng/fly after the perfuming treatment indicated on the X axis. Female flies were perfumed with blank hexane (control), synthetic 9P, synthetic 9H, or 9P+9H; untreated males are shown for comparison. Each dot represents a pool of 4 females or a single male. Dots and dashed lines are point estimates and 95% confidence for each treatment based on the regression approach described in Methods. The significance of changes was determined by ANOVA, followed by pairwise comparison using Tukey’s method. (B, B’) Total abundance of 9H (B) and change in the abundance of 9H (B’) after the perfuming treatment indicated on the X axis. (C) Stacked bar plots of copulation success after the perfuming treatment indicated on the X axis (N = 32 for each). Z-tests were performed on coefficients from logistic regression to determine the p-value for each perfuming treatment. P values are as follows: *** p < 0.001, ** p < 0.01, *, p < 0.05, × p < 0.1.

In mating trials, nearly all males encountered their female partners at the mating arena (Fig. S1 A). Males rarely showed threatening behavior toward the perfumed females, a stereotypical aggressive behavior displayed towards other males (Fig. S1 B), suggesting that males could still recognize the sex identities of females with modified CHC profiles. In the 9H treatment, we observed a non-significant decrease in the rate of courtship initialization (N = 32, logistic regression z-test, p = 0.25, Fig. S1 C) and leg vibration (p = 0.066, Fig. S1 D), which may suggest reduced motivation in males. Despite the non-significant effects on the individual elements of courtship behavior, we found a strong decrease in copulation success when females were perfumed with either 9P (N = 32, p = 0.0475) or 9H (p = 0.023, Fig. 1 C) compared with the hexane control (59%, N = 32). The proportion of pairs that mated in the 9H treatment (28% mated) was reduced by half compared to the hexane control (59% mated). The success rate was reduced less in the 9P treatment (34% mated), even though a larger amount of the synthetic CHC was introduced. This disparity may suggest that 9H was perceived as a stronger masculinity cue than 9P, and therefore outweighed 9P in mate evaluation and decision-making during courtship. Simultaneous perfuming with both 9P and 9H did not result in further inhibition of courtship and copulation (Fig. 1; Fig. S1).

A key *Drosophila* pheromone, *cis*-vaccenyl-acetate (cVA), is transferred from males to females during mating and subsequently inhibits courtship by rival males (Jallon et al. 1981; Ferveur and Sureau 1996; Everaerts et al. 2010; Ng et al. 2014). Male-biased CHCs are also transferred to females in many other *Drosophila* species (Khallaf et al. 2021). We therefore tested whether *D. prolongata* males transferred 9P or 9H to females during mating. However, no transfer was observed (Fig. S2), suggesting that while these CHCs reduce female attractiveness, they are unlikely to be involved in chemical mate guarding or male-male competition in a manner similar to cVA.

### Gene expression shows stronger sexual dimorphism in *D. prolongata* than in *D. carrolli*

We previously showed that sexually dimorphic pheromone profiles, with an increased abundance of 9P and 9H in males, have evolved in *D. prolongata* from a sexually monomorphic ancestor (Luo et al. 2019). To identify the genes responsible for this evolutionary transition, we performed RNA sequencing on dissected oenocyte-enriched tissues (abbreviated as oenocyte dissections) in sexually mature adults of both sexes of *D. prolongata* and *D. carrolli*, followed by differential gene expression analysis. We defined our candidate genes as those that show (1) differential expression between males and females in *D. prolongata* (Fig. 2A) and (2) differential expression between males of *D. prolongata* and *D. carrolli* (Fig. 2B). To also account for the possibility that both *D. prolongata* and *D. carrolli* are sexually dimorphic, but the extent or direction of sex bias differs between the two species, we also required that the differentially expressed genes show interaction effects between species and sex (Fig. 2C).

**Figure 2.**
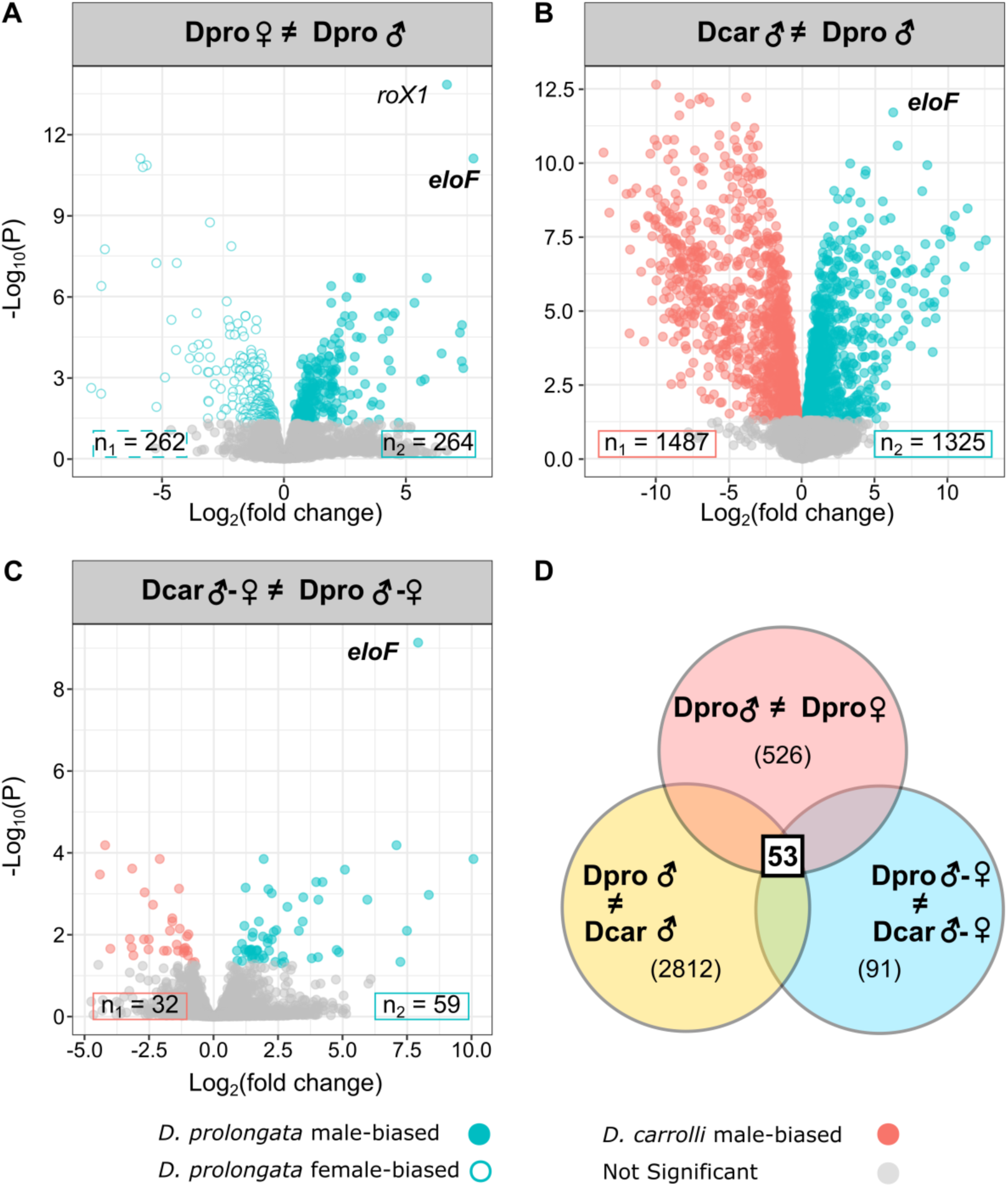
Differential expression analysis reveals strongly male-biased expression of *eloF* in *D. prolongata*. Volcano plots showing genes with differential expression between *D. prolongata* males and females (A), differential expression between *D. prolongata* and *D. carrolli* males, and interaction effects between species and sex (C). The interaction effects in (C) indicate that either the magnitude of sex differences varies between species, or the direction of sex bias is flipped between species. The x-axis is the log_2_ fold difference, and the y-axis is the negative log_10_ of FDR-adjusted P values. Numbers of genes that pass the FDR < 0.05 cutoff for biased expression in either direction are indicated in boxes. (D) Venn diagram showing candidate gene selection criteria, with 53 final candidates. Numbers of differentially expressed genes (FDR < 0.05) are labeled in parentheses for each one-way comparison.

In the comparison between male and female *D. prolongata*, 526 genes were identified as differentially expressed (Fig. 2A). Differentially expressed genes (DEGs) are almost equally likely to be female-biased (262 genes) as male-biased (264 genes). We found many more genes (2812) that were differentially expressed between males of *D. prolongata* and *D. carrolli* (Fig. 2B). These genes were slightly more likely to be enriched in *D. carrolli* (Binomial test, p = 2.4e-3), with 46.7% (1325 genes) having higher expression in *D. prolongata*. 91 genes showed significant interaction between species and sex (Fig. 2C). Consistent with *D. prolongata* being more sexually dimorphic in various phenotypes, the latter DEGs are more likely to show stronger sexual dimorphism in *D. prolongata* than in *D. carrolli* (Binomial test, p = 2.6e-10), with only 17.6% (16 genes) showing stronger dimorphism in *D. carrolli*.

### Sexually dimorphic and species-biased genes are enriched for lipid metabolism functions

The sexually dimorphic pheromone profile of *D. prolongata* is mainly attributable to a lower abundance of the shorter-chain 9T, and a higher abundance of the longer-chain 9P and 9H, in males (Luo et al. 2019). These compounds differ only in the number of carbons, suggesting a simple chemical basis for their differences – namely, a higher carbon chain elongation activity in males compared to females. To identify the molecular pathways that may underlie male-female differences in CHC profiles, we performed Gene Ontology (GO) enrichment analysis of the genes that show sex-biased expression in *D. prolongata* and oenocyte expression in *D. melanogaster*. We found 26 significantly enriched GO terms, of which the top 6 categories are all associated with lipid metabolism (Table 1; Fig. 3; Fig. S3).

**Figure 3.**
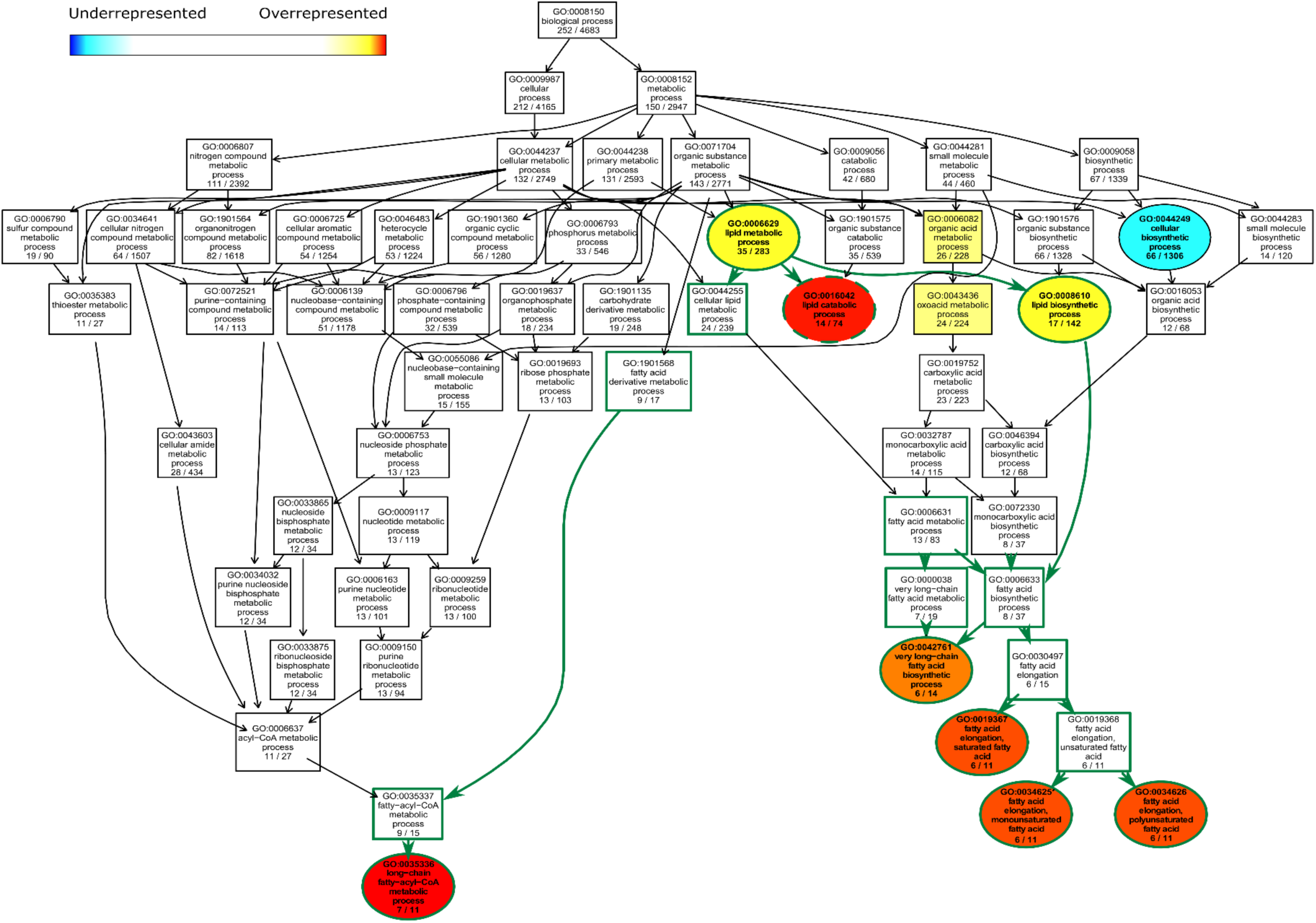
Terminal processes of lipid metabolism show differential gene expression between males and females of *D. prolongata*. Directed Acyclic Graph (DAG) of significant biological process GO terms and their parent terms. Significant (p < 0.05) and non-significant GO terms are color-coded and represented by ellipses and rectangular boxes, respectively. Significant GO terms can be underrepresented (blue) or overrepresented (red) based on Fisher’s exact test. Arrows indicate hierarchical relationships. GO terms at the same hierarchical level are placed at the same vertical position. GO terms under the lipid metabolic process (GO:0006629) are connected by green arrows and have green borders. Significant GO terms that are also enriched between males and females of *D. carrolli* have dashed borders.

**Table 1.** Significant GO terms in the comparison between *D. prolongata* males and females.

| GO.ID | Term | Annotated | Significant | Expected | Fisher | KS | Rank in<br>KS | Rank in<br>Fisher | Mean<br>rank |
| --- | --- | --- | --- | --- | --- | --- | --- | --- | --- |
| GO:0035336 | long-chain fatty-acyl-CoA<br>metabolic process | 11 | 7 | 0.59 | 3.30E-07 | 7.00E-05 | 3 | 1 | 2 |
| GO:0016042 | lipid catabolic process | 74 | 14 | 3.98 | 1.40E-06 | 0.00068 | 6 | 2 | 4 |
| GO:0019367 | fatty acid elongation,<br>saturated fatty acid | 11 | 6 | 0.59 | 8.40E-06 | 0.00221 | 13 | 3 | 8 |
| GO:0034625 | fatty acid elongation,<br>monounsaturated fatty acid | 11 | 6 | 0.59 | 8.40E-06 | 0.00221 | 14 | 4 | 9 |
| GO:0034626 | fatty acid elongation,<br>polyunsaturated fatty acid | 11 | 6 | 0.59 | 8.40E-06 | 0.00221 | 15 | 5 | 10 |
| GO:0042761 | very long-chain fatty acid<br>biosynthetic process | 14 | 6 | 0.75 | 4.80E-05 | 0.00895 | 30 | 6 | 18 |
| GO:0007548 | sex differentiation | 58 | 8 | 3.12 | 0.00025 | 0.01291 | 39 | 7 | 23 |
| GO:0055085 | transmembrane transport | 283 | 26 | 15.23 | 0.00035 | 1.00E-04 | 4 | 8 | 6 |
| GO:0070887 | cellular response to chemical<br>stimulus | 306 | 21 | 16.47 | 0.00039 | 0.01185 | 36 | 9 | 22.5 |
| GO:0006836 | neurotransmitter transport | 79 | 5 | 4.25 | 0.00087 | 0.00393 | 19 | 11 | 15 |
| GO:0007530 | sex determination | 18 | 5 | 0.97 | 0.00208 | 0.03854 | 78 | 13 | 45.5 |
| GO:0035725 | sodium ion transmembrane<br>transport | 19 | 5 | 1.02 | 0.0027 | 0.02581 | 62 | 14 | 38 |
| GO:0008284 | positive regulation of cell population proliferation | 57 | 9 | 3.07 | 0.00546 | 0.00172 | 11 | 17 | 14 |
| GO:0007472 | wing disc morphogenesis | 213 | 20 | 11.46 | 0.00656 | 0.02081 | 52 | 18 | 35 |
| GO:0044248 | cellular catabolic process | 618 | 36 | 33.26 | 0.00667 | 0.00114 | 9 | 19 | 14 |
| GO:0009063 | cellular amino acid catabolic process | 33 | 6 | 1.78 | 0.00742 | 0.00049 | 5 | 20 | 12.5 |
| GO:0015849 | organic acid transport | 37 | 7 | 1.99 | 0.00814 | 0.00692 | 26 | 21 | 23.5 |
| GO:1901606 | alpha-amino acid catabolic process | 25 | 5 | 1.35 | 0.00949 | 0.00782 | 28 | 24 | 26 |
| GO:0014019 | neuroblast development | 10 | 3 | 0.54 | 0.01394 | 0.01152 | 35 | 27 | 31 |
| GO:1901361 | organic cyclic compound catabolic process | 100 | 5 | 5.38 | 0.01437 | 0.04982 | 96 | 31 | 63.5 |
| GO:0006835 | dicarboxylic acid transport | 12 | 3 | 0.65 | 0.02359 | 0.03147 | 70 | 37 | 53.5 |
| GO:0008610 | lipid biosynthetic process | 142 | 17 | 7.64 | 0.02438 | 0.00522 | 23 | 39 | 31 |
| GO:0006629 | lipid metabolic process | 283 | 35 | 15.23 | 0.02957 | 0.03956 | 81 | 45 | 63 |
| GO:0044249 | cellular biosynthetic process | 1306 | 66 | 70.28 | 0.03223 | 0.02261 | 55 | 46 | 50.5 |
| GO:0006869 | lipid transport | 47 | 6 | 2.53 | 0.03681 | 0.0108 | 33 | 52 | 42.5 |
| GO:0048871 | multicellular organismal homeostasis | 51 | 6 | 2.74 | 0.03703 | 0.04385 | 86 | 53 | 69.5 |
| GO:0050795 | regulation of behavior | 56 | 9 | 3.01 | 0.04932 | 0.01687 | 43 | 62 | 52.5 |

Fisher: raw p-values from Fisher's exact test
KS: raw p-values from the Kolmogorov-Smirnov test

Long-chain fatty acyl CoA metabolic process (GO:0035336) shows particular enrichment in *D. prolongata* males compared to females (Table 1; Fig. 3; Fig. S3). These genes include seven fatty acyl reductases (FARs): CG17560, CG17562, CG14893, CG4020, CG5065, CG8306, and CG30427 (Chiang et al. 2016; Finet et al. 2019) and six fatty acid elongases (FAEs), including *elongase F* (*eloF*), CG9458, CG16904, CG9459, CG33110, and *bond*. Some members of both FAR and FAE gene families have been shown to affect the production and relative ratios of long-chain and short-chain pheromones (Chertemps et al. 2007; Dembeck et al. 2015; Pei et al. 2021; Rusuwa et al. 2022). We also found an enrichment of genes associated with transmembrane transport (26 genes, GO:0055085) (Table 1; Fig. S3). This may reflect the need for CHCs to be transported from the oenocytes to the cuticle, which likely involves crossing the intervening epithelium and several layers of cell membrane (Blomquist and Bagnères 2010). As expected, we also found an enrichment of genes involved in somatic sex differentiation (GO:0007548), including *Sex-lethal* (*Sxl*), *transformer* (*tra*), *doublesex* (*dsx*), and yolk proteins *yp1*, *yp2*, and *yp3*, which are known molecular targets of Dsx (Williams and Carroll 2009; Kopp 2012; Hopkins and Kopp 2021).

In the GO enrichment analysis of genes that show differential expression between *D. prolongata* and *D. carrolli* males and oenocyte expression in *D. melanogaster,* we identified 32 overrepresented and 7 underrepresented GO terms (Table S1; Fig. S4). As in the sex bias analysis, the enriched GO terms include terminal lipid metabolism processes, such as fatty acid elongation (GO: 0030497, see child GO terms GO:0034625, GO:0034626 and GO:0019367 in Fig. S4). A closely related process is the very-long-chain fatty acid biosynthetic process (GO:0042761), which contains the 3-hydroxy-acyl-CoA-dehydratase (HACD) *Hacd2* and the trans-enoyl-CoA-reductase (TER) *Sc2,* which exhibits extremely *D. carrolli*-biased expression (>12,000-fold change) and is sexually monomorphic in *D. prolongata*. Both HACDs and TERs are required for the elongation step during the synthesis of very-long-chain fatty acids, as is another enzyme class, 3-keto-acyl-CoA-reductases (KAR, (Wicker-Thomas et al. 2015)). Among predicted KARs, CG13284 showed differential expression between *D. prolongata* and *D. carrolli*. The long-chain fatty acyl CoA metabolic process (GO:0035336) also showed strong enrichment in this analysis (Table S1; Fig. S4). In addition to the 5 FARs identified in the male-female comparison, we detected significant differential expression of CG18031, a FAR that was previously shown to function in larval oenocytes (Cinnamon et al. 2016).

Beyond the genes immediately related to the pheromone synthesis pathway, we also observed wider differences in lipid metabolism. The lipid catabolic process (GO:0016042), in addition to being male-biased in *D. prolongata,* also shows higher expression in *D. prolongata* compared to *D. carrolli* (Fig. 3, Fig. S4). This GO term contains 74 species-biased genes, many of which have annotated or predicted function in lipid storage, mobilization, and transport. Representative examples include the medium-chain acyl-CoA dehydrogenase *Mcad* (Course et al. 2018), hormone-sensitive lipase *Hsl* (Bi et al. 2012), juvenile hormone epoxide hydrolases *Jheh1* (Campbell et al. 1992) and *Jheh2* (González et al. 2009), ABC-type fatty-acyl-CoA transporter *ABCD* (Gordon et al. 2018), long-chain-3-hydroxyacyl-CoA dehydrogenase *Mtpα* (Kishita et al. 2012), and predicted acetyl-CoA C-acetyltransferase *yip2* (Larkin et al. 2021).

In summary, GO enrichment analyses indicate that, consistent with the lipidic nature of CHCs, a disproportionately high number of genes involved in lipid metabolism are differentially expressed between males and females and between *D. prolongata* and *D. carrolli*. These genes, in particular fatty acid elongases and fatty acyl reductases, could underlie the evolution of sexually dimorphic pheromone profiles in *D. prolongata*.

### Candidate genes show increased male bias in *D. prolongata*

By intersecting the three selection criteria (male vs. female *D. prolongata* (Fig. 2A), male *D. prolongata* vs. *D. carrolli* (Fig. 2B), and interaction effects of species and sex (Fig. 2C)), we reduced the number of top candidate genes to 53, most of which show their highest expression in *D. prolongata* males (Fig. 2D). To test for correlated expression among these genes, we performed hierarchical clustering of genes and samples. The samples of *D. prolongata* males showed the greatest differences from the other samples (Fig. 4, left). We identified four major clusters of genes with distinct expression patterns (Fig. 4, top). The largest two clusters (red and purple) consist of 39 genes that show higher expression in *D. prolongata* males compared both to *D. carrolli* and to conspecific females. Most of these genes do not show significant sex differences in *D. carrolli*. Compared to the monomorphic *D. carrolli*, genes in the red cluster (24 genes) are strongly upregulated in *D. prolongata* males, while those in the purple cluster (15 genes) are downregulated in *D. prolongata* females (Fig. 4).

**Figure 4.**
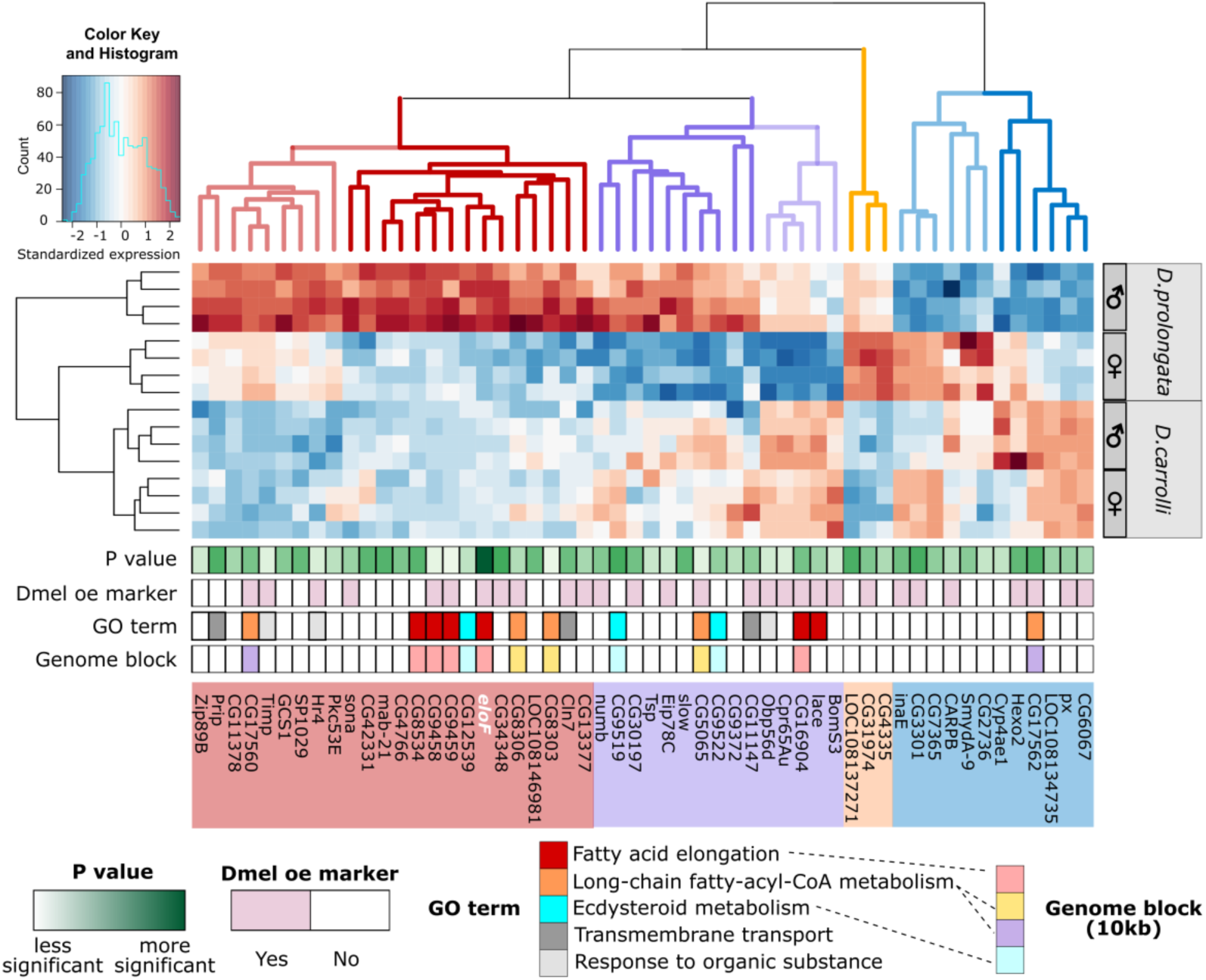
Candidate genes that show sex- and species-biased expression are involved in fatty acid biosynthesis and are arranged in gene clusters. Heatmap showing expression levels, standardized across samples, of 53 candidate genes (Fig. 2) + CG9459 (a member of the 5-elongase cluster), with red for relatively high expression and blue for low expression. UPGMA was used to perform hierarchical clustering on columns (genes) and rows (samples) based on pairwise Euclidean distances. The dendrogram of genes was cut into four clusters based on distinct co-expression profiles (e.g., red branches showing upregulation in *D. prolongata* males). FDR-adjusted p values from the 3-way comparison are annotated from light green (less significant) to dark green (more significant). Genes expressed in *D. melanogaster* oenocytes (Dmel oe expressors) (Li et al., 2021) are colored in light pink. Gene Ontology (GO) terms for enriched biological processes and candidates that fall in the same genome block (10kb neighborhood) are annotated by color as shown. Dashed lines indicate relationships between GO annotation and genomic clusters.

In principle, the evolution of male-biased pheromone profiles in *D. prolongata* could be explained either by species-specific increase or by species-specific reduction in the expression of genes in the pheromone biosynthesis pathway. The former pattern appears to dominate. In the third-largest cluster (blue in Fig. 4, 12 genes), most genes have mildly dimorphic expression in *D. carrolli* and increased dimorphism in *D. prolongata*, while fewer have overall lower expression in *D. prolongata*. The last and smallest cluster (orange, 3 genes) shows generally higher expression in *D. prolongata*, especially in females (Fig. 4).

In summary, *D. prolongata* males show a distinctive gene expression profile due mainly to male-specific upregulation of multiple genes. While the genes showing monomorphic or female-biased expression in *D. prolongata* (orange and blue clusters) may be necessary for the synthesis of species-specific pheromones, the genes directly responsible for the male-biased pheromone profile are more likely to be part of the male-enriched red and orange clusters. These clusters contain a number of fatty acid elongases and other genes involved in fatty acid metabolism (Fig. 4), whose increased expression may account for the evolution of male-specific CHC profile in *D. prolongata*.

### Functionally related genes are distributed in local genomic blocks

Genes involved in CHC metabolism are likely to be expressed in the oenocytes of other *Drosophila* species, including the well-studied model *D. melanogaster*. We intersected the 53 candidate genes identified above with the oenocyte-expressed genes from the *D. melanogaster* Fly Cell Atlas data (Li et al. 2021). For genes that do not have gene-level annotations, we inferred their functions from the associated GO terms. Most functionally related candidates, including genes from the very-long-chain fatty-acyl-CoA metabolic process (CG17560, CG17562, CG8306, CG5065) and fatty acid elongation (*eloF*, CG9458, CG9459, CG16904) show oenocyte expression in *D. melanogaster* (Fig. 4). We cannot rule out that some of the other genes are not detected in the Fly Cell Atlas due to the gene drop-out typical of single-nucleus data, and or that some genes are expressed in oenocytes in *D. prolongata* but not in *D. melanogaster* or vice versa.

We found that many candidate genes are spatially clustered in the genome (Fig. 4). In particular, the fatty acid elongases *eloF*, CG8534, CG9458, CG9459, and CG16904 are all located in a ∼10 kb genomic neighborhood (Fig. 4). Except for CG16904, which shows reduced expression in *D. prolongata* females compared to *D. carrolli*, the other four genes in this cluster have increased expression in *D. prolongata* males. Between *D. prolongata* males and females, *eloF* is 217-fold enriched in males, CG8534 156-fold, CG16904 87-fold, CG9458 148-fold, and CG9459 54-fold (Fig. 5A). This combination of spatial clustering and common sex bias suggests that these genes may share some *cis*-regulatory elements.

**Figure 5.**
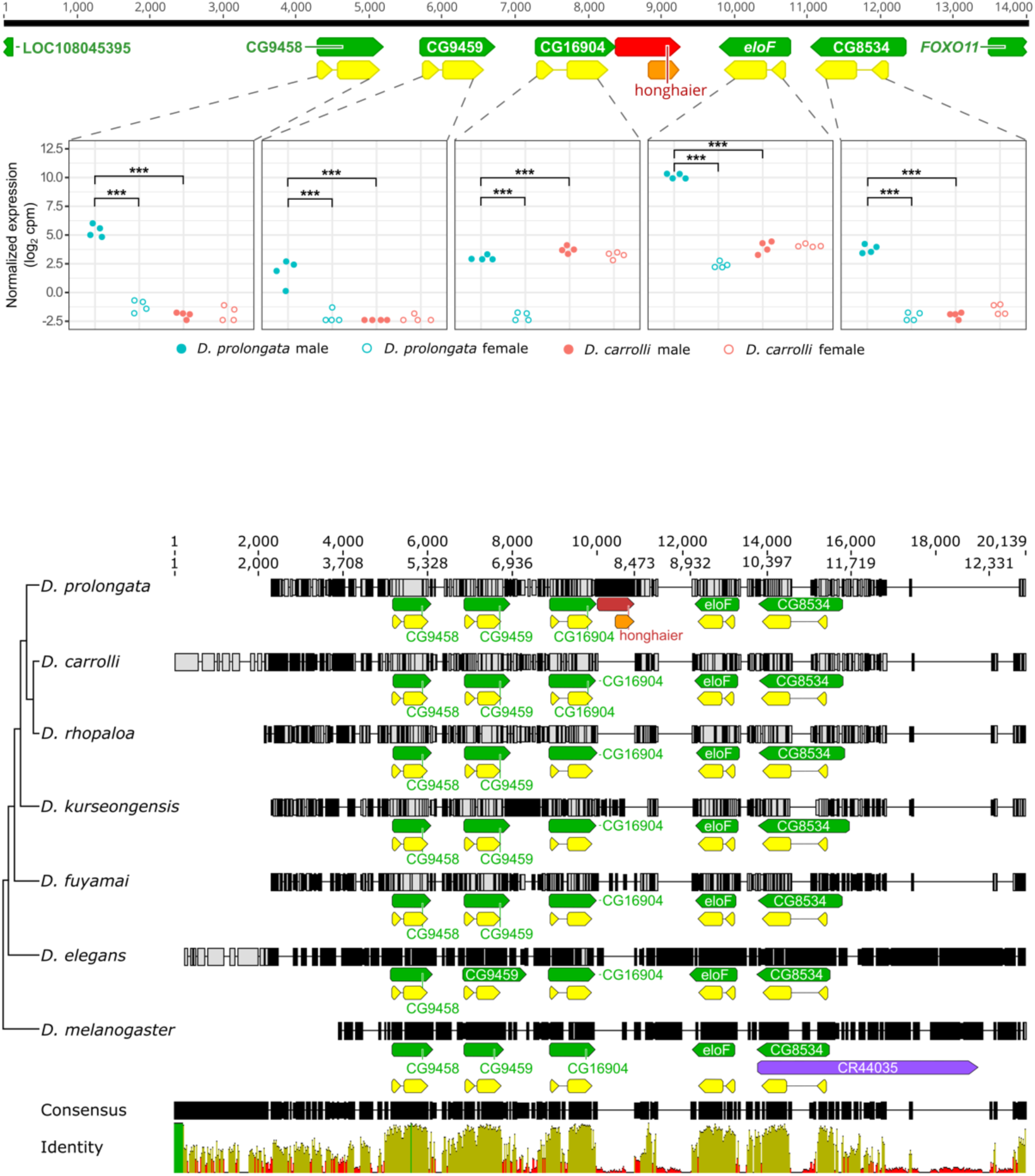
Structure and expression of the 5-elongase cluster. (A) All 5 elongases show a concerted expression increase in *D. prolongata* males. Dot plots showing normalized expression levels of each gene (RNA-seq data in log_2_ cpm). For each group, four biological replicates are represented by jitter points, color-coded by species. Males are in filled symbols; females are in open symbols. *** P < 0.001, ** P < 0.01, * P < 0.05. The structure of the ∼14kb genomic neighborhood is displayed on top. Numbers above the consensus sequence constructed from the reference genomes of *D. prolongata* and *D. carrolli* are coordinates showing the alignment length. Feature annotations are shown with green boxes representing genes, yellow boxes representing CDS, and the orange box representing a predicted ORF in the *honghaier* insertion, a TE-like repetitive sequence colored in red. The direction of all features is indicated. (B) The genomic organization of the 5-elongase cluster is conserved. Multiple alignment of DNA sequence across seven species, with species phylogeny on the left and consensus sequence at the bottom. Numbers above all sequences are coordinates showing the length of the consensus (12,702bp) or alignment (20,139bp). For each species, site-wise disagreement with the consensus is represented in a vertical gray line for nucleotide substitutions, a vertical black line for nucleotide insertions, and a horizontal line for nucleotide deletions. Feature annotations are displayed as in (A), with the additional purple box representing an antisense RNA. Percent identity per nucleotide across all species is displayed below the consensus sequence, with green indicating perfect (100%) agreement, yellow indicating intermediate (30-99%) agreement, and red indicating low (<30%) agreement.

The best-studied elongase gene is *eloF*, which encodes a *bona fide* fatty acid elongase. Oenocyte-specific knockdown of *eloF* in *D. melanogaster* leads to a reduction in the abundance of long-chain hydrocarbons, which are female-specific in that species (Chertemps et al. 2007). On the other hand, oenocyte-specific knockdown of CG9458 is not sufficient to change the balance between long- and short-chain CHCs in *D. melanogaster* (Dembeck et al. 2015).

Another example of spatial clustering is found among 5 fatty acyl reductases. CG8303, CG8306, and CG5065 are tandemly arranged in the genome (Fig. 4). These FARs are likely to be involved in essential lipid metabolism as RNAi knockdown leads to lethality (Finet et al. 2019). Two other FARs, CG17560 and CG17562, are located in a separate genomic cluster. Oenocyte-specific knockdown of CG17562 affects the production of short-chain mono alkenes and long-chain alkanes in *D. melanogaster* females (Chiang et al. 2016). Lastly, a local cluster is formed by three genes involved in ecdysteroid metabolism (CG9519, CG9522, and CG12539). Ecdysone regulates pheromone biosynthesis in *D. melanogaster* (Chiang et al. 2016; Baron et al. 2018) and houseflies (Adams et al. 1984; Blomquist et al. 1984; Blomquist et al. 1992; Blomquist et al. 1995). Interestingly, *hormone receptor 4* (*Hr4*), which encodes a nuclear receptor responding to ecdysone (King-Jones et al. 2005), is also among the 53 candidate genes. All 4 genes (CG9519, CG9522, CG12539, and *Hr4*) show strongly sexually dimorphic expression in *D. prolongata* while being sexually monomorphic in *D. carrolli* (Fig. 4). In conclusion, it is possible that correlated changes in the expression of genes involved in CHC synthesis were facilitated in part by their clustered genomic arrangement.

### Fatty-acid elongase *eloF* shows extremely male-biased expression in *D. prolongata*

Among the 53 candidate genes, we identified *elongase F* (*eloF*) as the top candidate underlying the observed sexual dimorphism of CHC profiles in *D. prolongata* (p = 7.29e-10, Fig. 2A-C, Fig. 4). Expression of this gene is strongly male-biased in *D. prolongata*, with a 217-fold difference between males and females based on RNA-seq data, but is not sexually dimorphic in *D. carrolli* (p = 0.80, Fig. 5). The only other gene with a comparable sex bias is *roX1*, a long non-coding RNA involved in X-chromosome dosage compensation. Moreover, *eloF* shows 79-fold higher expression in *D. prolongata* males compared to *D. carrolli* males.

To validate these results, we used quantitative PCR (qPCR) to amplify *eloF* transcripts from an independent set of oenocyte dissections. qPCR results support the strong male bias in *D. prolongata* (3413-fold enrichment, Fig. 6A) and higher expression level in males of *D. prolongata* over *D. carrolli* (82-fold enrichment, Fig. 6A). Contrary to the RNA-seq results, qPCR results suggest a modest (4-fold) but significant male-biased *eloF* expression in *D. carrolli* (Fig. 6A). This is consistent with previously described CHC phenotypes, where longer-chain hydrocarbons are slightly more enriched in males than females of *D. carrolli* (Luo et al. 2019). Despite this discrepancy between RNA-seq and qPCR, it is clear that sex differences are far less pronounced in *D. carrolli*, suggesting that a transition toward a strongly sexually dimorphic expression of *eloF* has occurred in *D. prolongata*.

**Figure 6.**
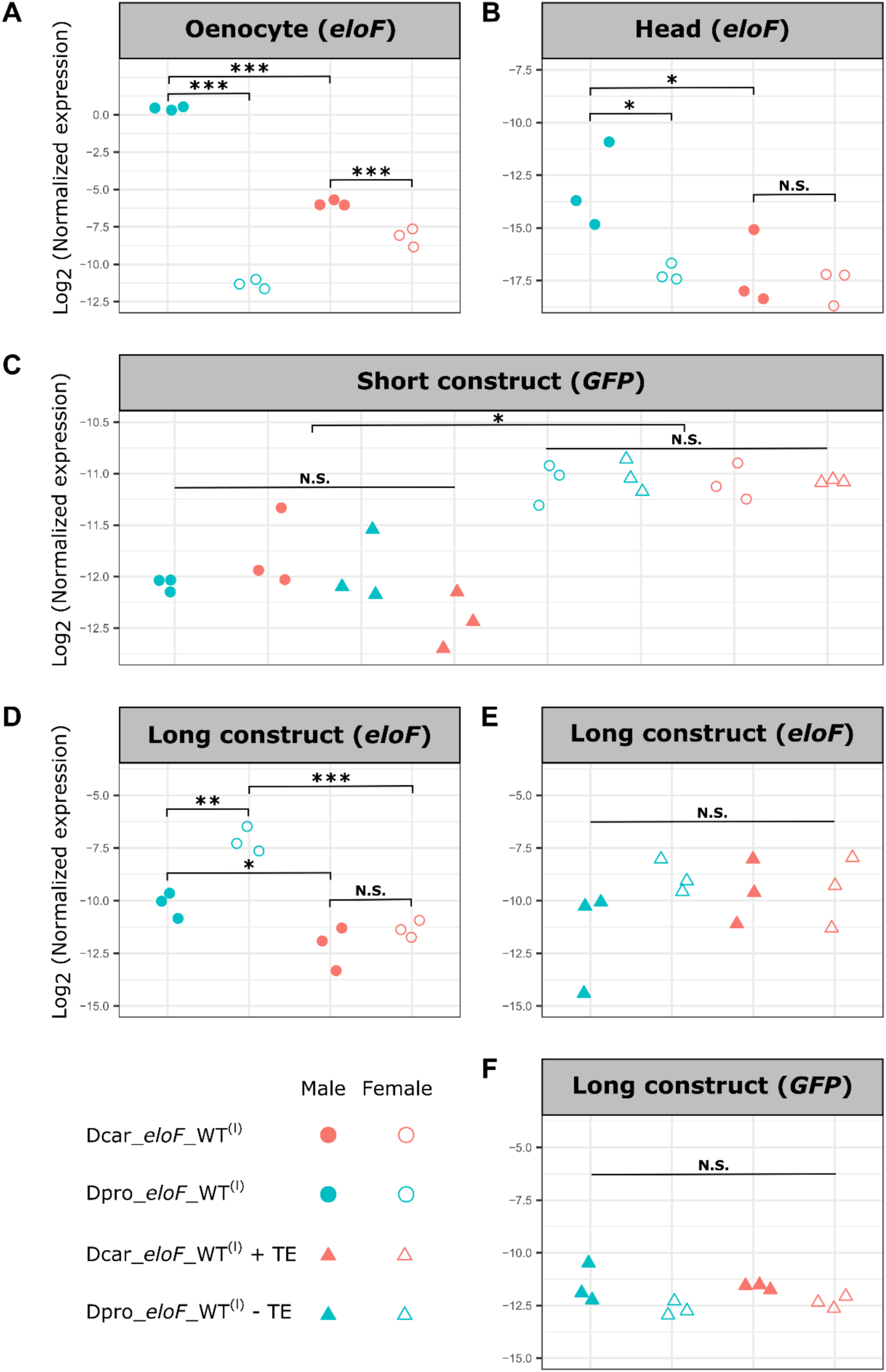
qPCR quantification of native *eloF* and transgenic reporter expression. Y axis shows the relative expression of *eloF* or GFP with respect to the reference gene *Rpl32* (measured in ΔCt). For each group, three biological replicates, each an average of three technical replicates, are represented by jitter points. Males are in filled symbols; females are in open symbols. Wild-type alleles are represented in circles and TE-swapped alleles in triangles. *** P < 0.001, ** P < 0.01, * P < 0.05, N.S.: not significant. (A) Male-biased *eloF* expression in oenocytes is much stronger in *D. prolongata* than in *D. carrolli*. (B) Head expression of *eloF* is also male-biased in *D. prolongata*, but sexually monomorphic in *D. carrolli*. (C) In the “short” reporter constructs containing only the downstream *eloF* region (Fig. S11), all *eloF* alleles have similar effects on GFP expression in transgenic *D. melanogaster*, with a slight (∼2-fold) female bias (p <0.05). (D) In the “long” reporter constructs containing the entire *eloF* locus (Fig. S11), the *D. prolongata* allele (Dpro_*eloF*_WT^(l)^), but not the *D. carrolli* allele (Dcar_*eloF*_WT^(l)^), causes sexually dimorphic expression of the donor *eloF* gene in transgenic *D. melanogaster*. (E, F) Removal of the *honghaier* TE from the *D. prolongata* allele (Dpro_*eloF*_WT^(l)^-TE) or addition of the *D. prolongata* TE to the *D. carrolli* allele (Dcar_*eloF*_WT^(l)^+TE) eliminates sex- and species-specific differences in the expression of *eloF* (E) and GFP (F).

To test whether the evolution of sexual dimorphism in *eloF* expression was tissue-specific, we also included in our qPCR study the heads of the same flies from which oenocyte samples were collected. In the brain, *eloF* shows little, if any, expression in either sex (Kudo et al. 2017). We found consistently low but detectable levels of *eloF* transcripts in the heads, which were several orders of magnitude lower than in oenocytes (Fig. 6B) and could be due to the presence of fat body tissue in the head. We also found that *eloF* expression was significantly higher in *D. prolongata* male heads than in the other groups (p < 0.05), resulting in a sexually dimorphic pattern in *D. prolongata* (27-fold difference) and a sexually monomorphic pattern in *D. carrolli*. Therefore, male-biased *eloF* expression is not entirely limited to oenocytes, although the extent of sexual dimorphism is much greater in oenocytes than in the head.

### Loss of *eloF* partially feminizes the pheromone profile of male *D. prolongata*

Increased expression of *eloF* correlates with the increased abundance of long-chain CHCs in male *D. prolongata*. The *D. melanogaster eloF*, which is a 1:1 ortholog of the *D. prolongata* gene, encodes a *bona fide* fatty acid elongase sufficient to elongate fatty acids in yeast heterologous expression assays, whereas its RNAi knockdown reduces the amount of female-biased long-chain hydrocarbons in *D. melanogaster* (Chertemps et al. 2007). This suggests that evolutionary changes in *eloF* expression could be responsible for the male-specific increase in the abundance of the 9P and 9H pheromones in *D. prolongata*. To test this hypothesis, we generated two loss-of-function *eloF* mutants (*eloF*[-]) in *D. prolongata* using CRISPR/Cas9 mutagenesis: an early frameshift resulting in a likely null allele and a 45 bp in-frame deletion, which disrupts a predicted transmembrane domain of EloF that is conserved with multiple mammalian fatty elongases (Chertemps et al. 2007) and may affect protein localization (Fig. 7A). Gas chromatography and mass spectrometry (GC-MS) analysis of these mutants and their wild-type progenitors showed that, qualitatively, they contained the same CHCs that were previously reported in wild-type *D. prolongata* (Table S3). The one exception is a minor alka-diene constituent, x,y-tricosadiene, which is shared between sexes and is not fully characterized.

**Figure 7.**
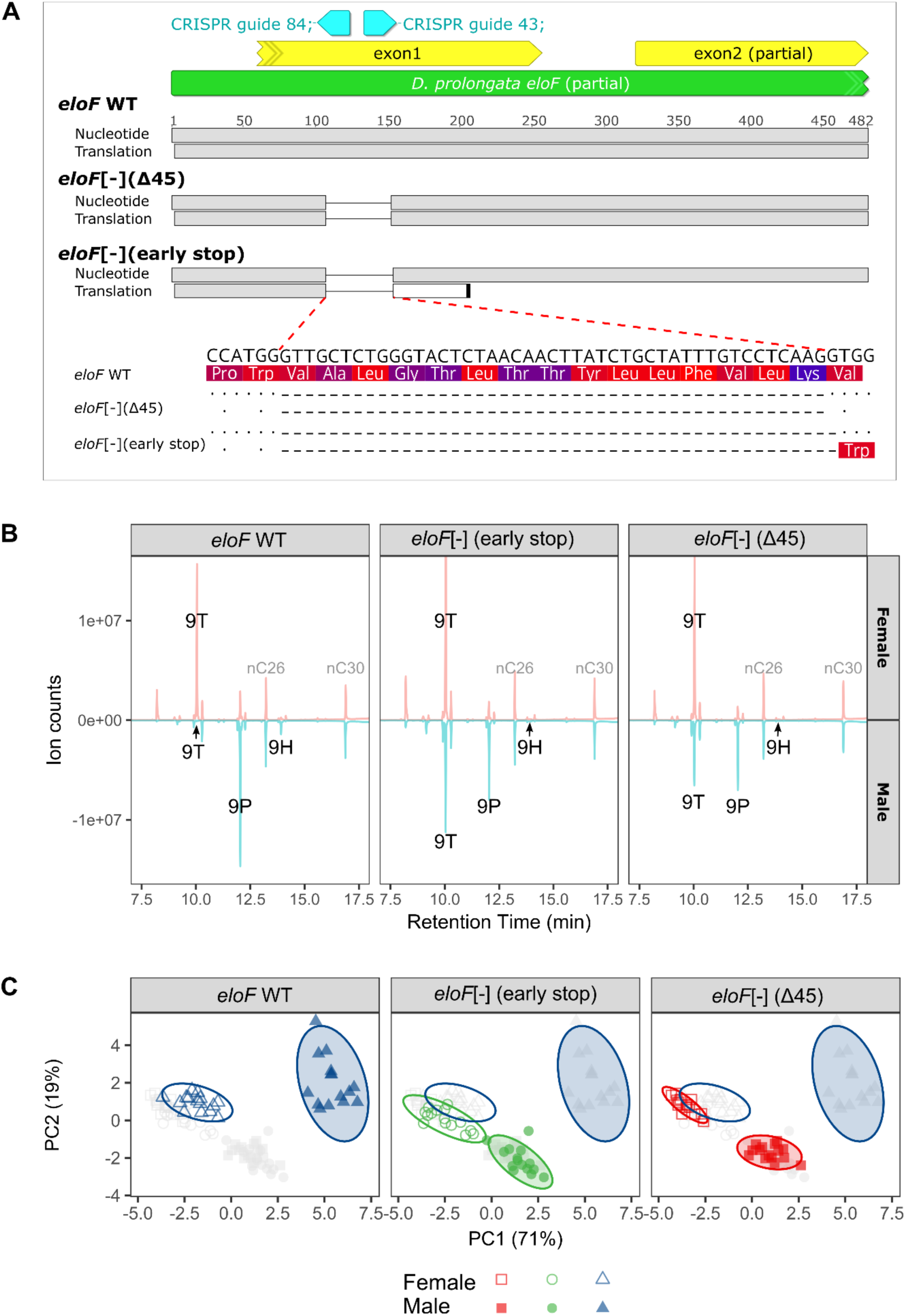
*eloF* mutations cause partial feminization of pheromone profiles in male *D. prolongata*. (A) Schematic diagram of two CRISPR mutant strains: one strain with a 45 bp deletion (“*eloF*[-] Δ45”) and the other with an early stop codon (“*eloF*[-] early stop”). Partial *eloF* locus is shown in green, first and part of second exon are in yellow, and the positions of the two guide RNAs used to generate these mutations are in cyan. The orientation of all features is indicated by arrows. Nucleotide sequences and their translations are shown, with deleted (dashed lines) and surrounding sequences zoomed in to show the amino acid changes. (B) GC traces of representative (closest to ellipse center) samples for each sex * genotype combination, with male signals (in blue) inverted relative to female signals (in red). Three 9-Monoenes (9T, 9P, 9H) that are most sexually dimorphic in wild-type *D. prolongata* are labeled, with two corresponding external standards (nC26, nC30) labeled in gray. (C) PCA ordination of logarithm transformed CHC abundances, partitioned by genotype. Axes are the first two principal components extracted from the variance-covariance matrix of 18 consensus CHCs (Table S3), with the % variance explained in parenthesis. The first two principal components collectively explain 90% of variation. Points, color-coded by genotype, represent samples, with females in open symbols and males in filled symbols. Ellipses represent 95% confidence regions constructed by bivariate t-distribution. Gray points representing the samples of other genotypes are embedded in each panel as a reference, with wild-type males and females indicated by open ellipses.

We observed a strong feminization of male pheromone profiles in both *eloF[-]* mutants (Fig. 7B, C; Fig. S5-6). In contrast, only a subtle effect is seen in females (Fig. 7C). Consistent with its molecular function, *eloF*[-] flies show decreased production of long-chain hydrocarbons, which is much more pronounced in males than in females (Fig. 7B; Fig. S5-6). In males, we found a ∼50% reduction in the amount of 9P and a near absence of 9H. Concurrently, *eloF*[-] males show increased abundance of 9T to a level comparable to wild-type females (Fig. 7B; Fig. S5). Since 9T is an early terminal product derived from a common metabolic precursor with 9P and 9H during carbon chain elongation, the increased abundance of 9T may be a direct consequence of reduced 9P and 9H synthesis. Notably, the total abundance of all CHCs as well as that of 9-monoenes remained unchanged (Fig. S7), indicating that disruption of *eloF* inhibits elongation of specific male-biased pheromone precursors without having a general inhibitory effect on CHC synthesis. While the degree of sexual dimorphism is reduced in *eloF[-]* mutants, CHC profiles remain dimorphic (Fig. 7C). This incomplete feminization suggests that other FAE or FAR genes, which also show strongly male-biased expression in *D. prolongata* (Fig. 4; Fig. 5A) may act in parallel with *eloF* in the production of long-chain CHCs.

*eloF[-]* mutations did not significantly affect courtship or copulation in single male-female pairs (Table S4) despite reduced 9H and 9P abundance in males. Instead, they affected male-male interactions, although the effects were not consistent between the two *eloF[-]* mutants. Males with the 45 bp deletion in the transmembrane domain of *eloF* showed increased rate of boxing, a typical male-male aggressive behavior in *D. prolongata* (Table S4), whereas decapitated males with the early stop codon elicited higher frequency of misdirected courtship from other males (Table S4). These results suggest that other signals must be involved in male-male communication alongside 9H and 9P.

### The elongase gene cluster including *eloF* is conserved

We compared the genomic neighborhood of the *eloF* locus between the *rhopaloa* species subgroup (*D. prolongata, D. fuyamai*, *D. kurseongensis*, *D. rhopaloa*, and *D. carrolli*), its nearest outgroup *D. elegans*, and *D. melanogaster*. In *D. melanogaster*, *eloF* is part of a ∼10 kb cluster with four other fatty acid elongases, which likely evolved by tandem duplication (Szafer-Glusman et al. 2008). We found the same five predicted elongases, in the same order and orientation and with the same exon/intron structure, in all species of our focal clade (Fig. 5B), indicating deep origin and strong conservation of the elongase cluster.

Despite the strong evidence of sex- and species-specific regulation of *eloF* (Fig. 2; Fig. 5A; Fig. 6A), it is possible that changes in EloF protein activity contribute to the derived male-specific CHC profile seen in *D. prolongata*. To test this hypothesis, we compared the coding sequences of *eloF* between *D. prolongata* and the other four species of the *rhopaloa* subgroup, which are sexually monomorphic in the abundance of 9P and 9H (Luo et al. 2019). There is a high overall degree of protein sequence conservation (>90%) in the coding region (Fig. S8), and the protein identity between *D. prolongata* and *D. carrolli* is 96.5%. While we found 20 single nucleotide variants (SNVs) distinguishing the reference genomes of these two species, with 9 of them resulting in predicted amino acid substitutions, our RNA-seq data show that all these substitutions are polymorphic in one or both species and none are fixed between species (Fig. S8, Table S5). In the absence of fixed amino acid differences between *D. prolongata* and *D. carrolli*, coding sequence divergence in *eloF* is unlikely to contribute to the evolution of CHC profiles.

### A species-specific transposable element insertion in *eloF* in *D. prolongata*

Our evidence points to changes in *eloF* transcription as the main cause of sex-specific pheromone profiles in *D. prolongata*. To identify the likely *cis*-regulatory elements of *eloF*, we examined the flanking intergenic and intronic regions of *eloF* in the *rhopaloa* subgroup. The most drastic difference between *D. prolongata* and all other species is a ∼900 bp insertion in the otherwise conserved (∼500bp, >72% sequence identity) downstream region of *eloF* (Fig. 5). The inserted sequence contains two predicted binding sites for the *doublesex* (*dsx*) transcription factor, the main regulator of somatic sexual differentiation in *Drosophila* and other insects (Williams and Carroll 2009; Kopp 2012; Hopkins and Kopp 2021), as well as several predicted binding sites for *bric-à-brac 1* (*bab1*), a TF that regulates the development of abdominal segments (Kopp et al. 2000; Rogers et al. 2013) (Fig. S9). These sites, along with the rest of the insertion, are absent in the other species. Our inspection of the single-cell Fly Cell Atlas (Li et al. 2021) shows that both TFs are expressed in adult oenocytes, and that the upstream regions of oenocyte-biased genes show a significant enrichment for *bab1* binding motifs (p = 1.75e-60). These observations suggest that the insertion in the 3’ region of *eloF* may have contributed to the species- and sex-specific increase in *eloF* expression in *D. prolongata*.

We found hundreds of highly similar copies of this insertion throughout the genome of *D. prolongata* (Table S6), suggesting that it may be a transposable elements (TE). We named this putative TE “*honghaier*” after the mythical Chinese character capable of self-duplication. *honghaier* is found in high copy numbers in all five species of the *rhopaloa* subgroup, but not in *D. elegans* or *D. melanogaster* (Table S6), suggesting that it originated or invaded this lineage relatively recently. *honghaier* is AT-rich (∼60%, Fig. S9), a common feature of miniature inverted-repeat transposable elements (MITE), and has a 414 bp open reading frame (ORF), which is predicted to be transcribed in the direction opposite to *eloF* (Fig. S9). The strongest sequence similarity between *honghaier* and known TEs is found with DNAREP1_DM (53%) (Kapitonov and Jurka 2003) and *wukong* (52.5%), a mosquito MITE element (Tu 1997). However, *honghaier* is unlikely to be a true MITE, as these elements usually do not have coding potential (Tu 1997). *honghaier* also lacks typical hallmarks of TEs such as terminal inverted repeats (TIRs) and flanking short direct repeats stemming from target site duplication (Senft and Macfarlan 2021).

Although changes in gene expression caused by TE insertions are common (Sundaram et al. 2014; Cridland et al. 2015; Hof et al. 2016; Trizzino et al. 2017; Barth et al. 2020), and the *honghaier* insertion in *eloF* correlates with its divergent expression profile in *D. prolongata*, we cannot rule out contributions from other *cis*-regulatory changes. There are multiple fixed SNVs between *D. prolongata* and *D. carrolli* in the upstream (∼300bp, Fig. S10) and intronic (∼70bp, Fig. S11) regions of *eloF,* despite overall high conservation (94.3% for upstream and 96.9% for intron). However, these substitutions do not affect any predicted binding motifs for *dsx*, *bab1*, or other TFs known to regulate abdominal development or sexual differentiation.

### *eloF* downstream region drives gene expression in oenocytes

To test whether increased expression of *eloF* in *D. prolongata* is due to changes in the *cis*-regulatory regions of *eloF*, we generated transgenic GFP reporter strains where the *eloF* loci from *D. prolongata* and *D. carrolli* were transformed into *D. melanogaster*. First, we cloned the entire *eloF* region between the flanking genes *CG16904* and *CG8534* (Dpro *eloF* WT^(l)^ and Dcar *eloF* WT^(l)^). In these constructs, the *eloF* transcript is in the same orientation as the GFP reporter, while the *honghaier* insertion is in the opposite orientation (Fig. S12). To examine the effects of the *honghaier* insertion, we also made two TE-swap constructs, one with *honghaier* removed from *D. prolongata* (Dpro *eloF* WT^(l)^-TE) and the other with *honghaier* added to *D. carrolli* (Dcar *eloF* WT^(l)^+TE). We observed little, if any, GFP expression by qPCR (Fig. 6F; Table S2). In females, GFP transcripts were not detectable (Ct > 40), while in males they were present at very low levels (Ct > 35). We then cloned only the downstream regions of *eloF*, generating both wild-type reporters (Dpro *eloF* WT^(s)^ and Dcar *eloF* WT^(s)^) and TE-swap constructs (Dpro *eloF* WT^(s)^-TE and Dcar *eloF* WT^(s)^+TE). These constructs were designed so that the downstream *eloF* sequences were upstream of the GFP reporter, and the *honghaier* insertion in the forward orientation relative to the promoter (Fig. S12).

In the adult dorsal abdominal epidermis of transgenic flies carrying short *eloF* reporters, we observed GFP expression in both sexes in stripes of tissue in the posterior half of each segment (Fig. S13). This region corresponds to the location of the pheromone-producing oenocytes (Billeter et al. 2009), suggesting that the downstream region of *eloF* contains an oenocyte-specific enhancer.

### *eloF* allele from *D. prolongata* drives sexually dimorphic expression in *D. melanogaster*

As both *D. prolongata* and *D. carrolli* express *eloF* in the abdomen, we expect the differences in enhancer activity to be more quantitative than qualitative. We therefore compared transgenic reporter activity by qPCR. In the long constructs, which contained the *eloF* coding sequence, we compared the *eloF* transcript levels. The *D. carrolli* allele was expressed at similar levels in males and females (Fig 6D). The *D. prolongata* allele was expressed at a ∼20-fold higher level than the *D. carrolli* allele and showed significant sexual dimorphism (Fig 6D). Surprisingly, the *D. prolongata* allele was expressed at a higher level in females compared to males. While this direction is opposite to what is observed at the endogenous *eloF* locus in *D. prolongata*, it matches the phenotype of *D. melanogaster*, in which *eloF* expression and long-chain CHC abundance are higher in females than in males (Chertemps et al. 2007). This indicates that while the *D. prolongata eloF* allele, in contrast to the *D. carrolli* allele, encodes sex-specific regulatory information, its effect on transcription depends on the *trans*-regulatory background, which appears to have diverged between *D. prolongata* and *D. melanogaster*. The removal of the *honghaier* insertion from the *D. prolongata* allele, or the addition of this insertion to the *D. carrolli* allele, eliminated the differences in reporter activity both between species and between males and females (Fig. 6E), suggesting that this insertion is necessary, but not sufficient, for driving sex-specific expression of *eloF*.

We then used the short reporter constructs to compare GFP transcript expression driven by the wild-type and TE-swapped alleles of the downstream *eloF* region that contains the *honghaier* insertion in *D. prolongata*. We observed a modest (2-fold) but significant sexual dimorphism, also in the direction of females having higher expression (Fig. 6C). However, GFP expression was low in both sexes (∼29 Ct), and there was no significant difference between the *D. prolongata* and *D. carrolli* alleles in either sex (Fig. 6C), suggesting that the downstream region and the *honghaier* insertion alone are not sufficient to confer species-specific transcriptional regulation, at least in the *D. melanogaster* genetic background. Alternatively, it is possible that *eloF* enhancers are sensitive to the sequence, position, and relative orientation of the interacting promoter.

## Discussion

In this study, we show that sexually dimorphic pheromones affect mating behavior in *D. prolongata* and identify a key gene underlying the evolution of sex-specific pheromone profiles in this species (Fig. 8). A *cis*-regulatory change in the *eloF* gene is an important, though not the only, component of the genetic changes that distinguish *D. prolongata* from its close, sexually monomorphic relatives. Below, we discuss these findings in the context of our still limited but growing knowledge of the evolution and functional roles of *Drosophila* pheromones.

**Figure 8.**
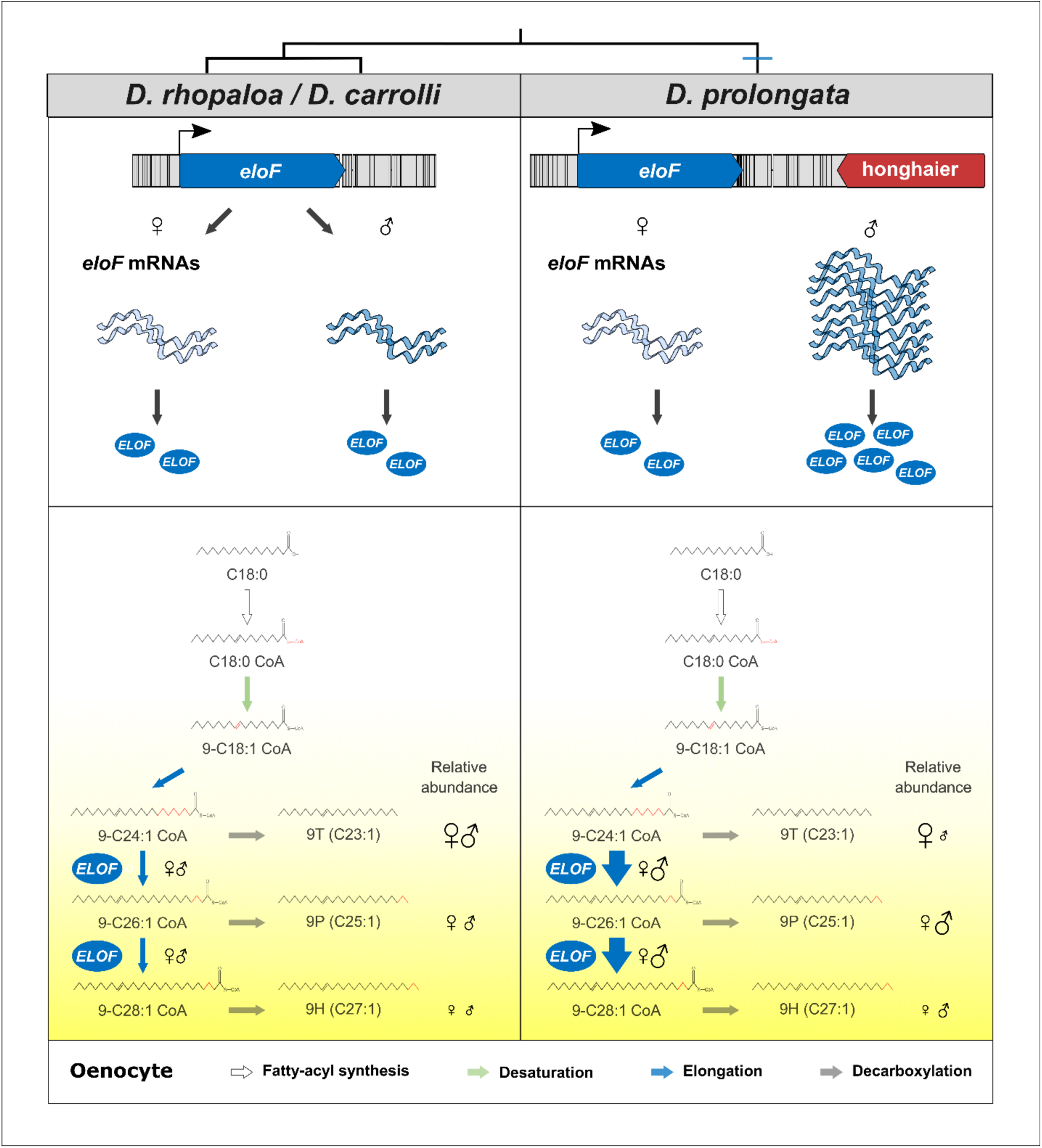
Proposed molecular mechanism underlying the evolution of sexually dimorphic CHCs in *D. prolongata*. Schematic diagram of the expression of *eloF* and the CHC biosynthetic pathway in adult oenocytes, showing quantitative differences between the sexually monomorphic *D. rhopaloa* and *D. carrolli* and the sexually dimorphic *D. prolongata*. Species phylogeny is on top. Colored arrows represent the four major steps in CHC synthesis. Illustrative chemical structures are shown below the substrates and products. Quantitative differences in reaction rate are indicated by arrow thickness, and the quantities of produced CHCs are indicated by the size of sex symbols (♀♂). The yellow shade gradient corresponds to the increasing carbon chain length of the metabolite. *ELOF* is the elongase F protein responsible for producing 9P and 9H from the shorter 9T precursor.

### Male-specific hydrocarbons reduce female mating success

Sex-specific visual, acoustic and chemical cues play vital roles in mate recognition. The divergence of communication systems helps maintain species boundaries and can drive the evolution of reproductive isolation, as seen in the coevolution of nuptial colors and color vision in sticklebacks (Boughman et al. 2005), wing color patterns and co-evolved mate preferences in *Heliconius* butterflies (Jiggins et al. 2004), matching conspecific mating duets sung by male and female lacewings (Wells and Henry 1992), or the divergent pheromone blends between two sympatric races of the European corn borer (Linn et al. 1997). In *Drosophila*, sexually dimorphic CHCs mediate mate recognition and allow males to differentiate potential mates from competitors (Howard and Blomquist 2005). For example, in *D. melanogaster*, the male-biased 7-tricosene (7T) evokes male-male aggression, whereas the female-biased 7,11-heptacosadiene (7,11-HD) elicits courtship behavior even when applied to a dummy female (Jallon 1984; Ferveur and Sureau 1996).

The male-biased 9P and 9H in *D. prolongata* may serve as one of the cues that facilitate mate recognition, though other signals including visual cues are clearly important (Takau and Matsuo 2022). Our perfuming studies show that 9H, and to a lesser extent 9P, reduce mating success when applied to females. This reduction is not due to a lack of courtship interest, but could instead be related to reduced leg vibration, a species-specific behavior performed by *D. prolongata* males that increases female receptivity (Setoguchi et al. 2014). This suggests that the lower relative amounts of 9P and 9H in females compared to males are important for the proper progression of male courtship toward females. Identifying other functions of 9P and 9H is complicated both by the fact that *eloF* mutations do not fully block the synthesis of these compounds, and by the complex mix of visual, chemical, and auditory signals that mediate *Drosophila* mating behavior. Some pheromones, such as *cis*-vaccenyl-acetate (cVA), are transferred from males to females during mating, and function as an anti-aphrodisiac signal (Jallon et al. 1981; Ferveur and Sureau 1996; Everaerts et al. 2010; Ng et al. 2014). However, we find no evidence that *D. prolongata* males transfer 9P or 9H to females, suggesting that the long-chain CHCs are unlikely to act by reducing female re-mating. The effect of long-chain CHCs on male-male interactions appears to be limited. While we observe an increase in male-male aggression and misdirected courtship toward males, these effects are inconsistent between the two mutant alleles of *eloF* although both alleles reduce the abundance of 9H and 9P and increase 9T levels.

Beyond intraspecific communication, 9P and 9H could contribute to sexual isolation between sibling species, similar to the roles of 7T and 7,11-HD in the isolation between *D. melanogaster* and *D. simulans* (Jallon 1984; Seeholzer et al. 2018). Pre-mating isolation between *D. prolongata* and its relatives is strong; we have never observed an interspecific mating. Female *D. prolongata* could potentially use a lack of 9P or 9H to reject mating attempts from males of other species, although, as in the intraspecific communication, this would likely be only one of several cues. Compared to all of its relatives, *D. prolongata* has highly derived male mating behavior and greatly exaggerated sexual dimorphism in multiple traits, including reversed sexual size dimorphism, pigmentation, and the organization of the chemosensory system (Setoguchi et al. 2014; Luecke and Kopp 2019; Luecke et al. 2022). In this context, deciphering the behavioral and ecological roles of 9P and 9H may elucidate why a strongly male-biased pheromone profile has evolved in *D. prolongata* but not in any of its close relatives.

### *eloF* is a major gene controlling long-chain pheromone production in male *D. prolongata*

Our results show that the evolution of sexually dimorphic pheromone profiles in *D. prolongata* is due to a large extent to changes at the *eloF* locus (Fig. 8). *eloF* mutations have a particularly strong effect on the elongation of C25 to C27, and a somewhat milder effect on the elongation of C23 to C25. *eloF* is a well-characterized fatty acid elongase that has been shown to catalyze the conversion of long-chain fatty acyl CoA to very long-chain fatty acyl CoA in yeast assays (Chertemps et al. 2007). In *D. melanogaster*, long-chain hydrocarbons are enriched in females (Ferveur, 2005; Ferveur & Jallon, 1996), and knocking down *eloF* expression in *D. melanogaster* elicits a female-specific reduction in their abundance. However, ectopic expression of *eloF* in *D. melanogaster* males does not increase the abundance of long-chain CHCs, indicating that *eloF* is necessary but not sufficient for their synthesis (Chertemps et al. 2007). Unlike other genes whose disruption leads to an overall increase or decrease in CHC production (Qiu et al. 2012; Dembeck et al. 2015; Wicker-Thomas et al. 2015), we show that *eloF* mutations in *D. prolongata* alter the relative abundance of short vs. long-chain monoenes without affecting total monoene amounts, or the amounts of CHC more generally.

In principle, increased EloF activity in *D. prolongata* could be due to either regulatory or coding sequence changes. Even small differences in protein sequence can have a major effect on enzyme function, with drastic phenotypic consequences (Nachman et al. 2003; Weill et al. 2003; Gratten et al. 2007). However, we find no fixed coding sequence differences in *eloF* between *D. prolongata* and its sibling species *D. carrolli*, in which the amounts of 9P and 9H are nearly monomorphic. On the other hand, *D. prolongata* shows extreme sexual dimorphism in *eloF* transcript abundance as well as overall higher *eloF* expression relative to *D. carrolli*. In *D. prolongata*, *eloF* expression is >3,000-fold higher in males than in females, while only a 4-fold difference between the sexes is detected in *D. carrolli*. These observations indicate that increased 9P and 9H production in *D. prolongata* males is due to changes in *eloF* expression rather than EloF protein activity (Fig. 8).

### Interaction of *cis*- and *trans*-regulatory factors in the control of sex-biased *eloF* expression

The *eloF* allele of *D. prolongata*, but not *D. carrolli*, drives sexually dimorphic gene expression in *D. melanogaster*, suggesting the presence of *cis*-regulatory elements that respond to the sexual differentiation pathway. This pathway, including the *doublesex* (*dsx*) transcription factor and the *transformer* (*tra*) RNA-binding protein that controls its sex-specific splicing, is the primary mediator of sex-specific cell differentiation in somatic tissues (Williams and Carroll 2009; Kopp 2012; Hopkins and Kopp 2021). Across a number of *Drosophila* species, the binding of Dsx to the regulatory region of the *desatF* (*Fad2*) gene is responsible for female-specific expression of that gene in adult oenocytes, and thus for the female-specific production of the 7,11-HD pheromone (Shirangi et al. 2009). The *D. melanogaster eloF* ortholog, which also contributes to the synthesis of female-specific pheromones, has also been shown to be under the control of *tra* (Chertemps et al. 2007). Overall, however, the regulatory program that controls sex-specific differentiation of oenocytes remains to be characterized.

Perhaps the most surprising part of our results is that the direction of sex bias was reversed in reporter assays. Instead of recapitulating the male-biased expression of *eloF* seen in *D. prolongata*, the *D. prolongata eloF* gene is expressed in a female-biased fashion when placed into the *D. melanogaster* genome. That is, the direction of sex bias replicates the pattern seen in the *D. melanogaster* host (Chertemps et al. 2007) and not in the *D. prolongata* donor. This indicates that, while the *eloF* locus itself encodes the potential for sexually dimorphic expression, the realization of this potential depends on the *trans*-regulatory background, which has clearly diverged between *D. prolongata* and *D. melanogaster*. The mechanistic basis of this reversal is not clear. It could indicate either that the regulation of *eloF* by the sexual differentiation pathway is indirect, or that *dsx* interacts with other transcription factors whose expression differs between species.

The most conspicuous sequence change at the *eloF* locus is the presence of a species-specific insertion of the *honghaier* TE-like element in *D. prolongata*. TEs are a major source of *cis*-regulatory changes underlying the evolution of gene expression (Bourque et al. 2008; Lynch et al. 2011; Sundaram et al. 2014; Chuong et al. 2017; Trizzino et al. 2017; Almojil et al. 2021; Herpin et al. 2021), so it is tempting to speculate that the *honghaier* insertion is at least partly responsible for the increased expression of *eloF* in *D. prolongata* males. Consistent with this idea, the *honghaier* element contains predicted binding sites for the *dsx* and *bab1* transcription factors. However, the results of reporter assays defy a simple explanation, as the downstream *eloF* region that contains the *honghaier* insertion in *D. prolongata* is not sufficient to confer sex- and species-specific expression observed in the longer reporters. Moreover, swapping the *honghaier* insertion between the *D. prolongata* and *D. carrolli* alleles shows that this insertion is necessary but not sufficient for species- and sex-specific expression. This suggests that the downstream region may interact with other parts of the *eloF* locus or the wider elongase cluster. Although enhancers are generally modular (Carroll 2008), numerous exceptions are known where interactions among several regions within the locus are necessary for correct gene expression (Luecke et al. 2022; Museridze et al. 2024). Another, not mutually exclusive explanation is that *eloF* enhancers are promoter-specific – that is, their activity depends on the sequence and the relative position and orientation of the interacting promoter (Ohtsuki et al. 1998; Kwon et al. 2009; Bergman et al. 2022).

Finally, we note that *eloF* is part of a compact genomic cluster with four other elongases, and that all five genes show strongly male-biased expression in *D. prolongata* but not in *D. carrolli*. This raises the possibility that their expression is controlled in part by shared *cis*-regulatory elements, and that some of the enhancers that control *eloF* expression may be located outside of the immediate *eloF* locus. Co-regulation of clustered genes is not uncommon. It contributes, for example, to the co-regulation of *Hox* (Gould et al. 1997) and *Iroquois* (*Irx*/*iro*) (Tena et al. 2011) genes in vertebrates, while in *Drosophila* shared enhancers contribute to the concerted expression of *bab*, *pdm*, and other closely linked genes (Bourbon et al. 2022; Loker and Mann 2022).

### Multiple genes likely contribute to sex-specific pheromone profiles in *D. prolongata*

Disruption of *eloF* causes a strong, but not complete feminization of the male CHC profile. In particular, while the longest-chain CHC, 9H, is almost absent in *eloF* mutants, the most abundant male-biased pheromone 9P shows only a ∼50% reduction in mutant males, and remains the most abundant CHC component. Since one of the mutant alleles, a premature stop codon early in the first exon, is almost certainly a molecular null, this suggests that other genes must contribute to the synthesis of 9P. Consistent with this, *eloF* knockdown in *D. melanogaster* reduces the amount of very long chain CHCs, but does not fully eliminate them (Chertemps et al. 2007; Wicker-Thomas et al. 2009).

RNA-seq shows that sex-specific expression of lipid metabolism genes in *D. prolongata* is not restricted to *eloF*. All four other elongases in the genomic cluster that contains *eloF* show strongly male-biased expression, as do many other enzymes that function in lipid metabolism. Unfortunately, most of these candidate genes have not been characterized nearly as well as *eloF*. In the elongase cluster, only *CG9458* has been studied so far, and its disruption does not apparently impact the elongation of pheromone precursors in *D. melanogaster* (Dembeck et al. 2015). One possibility is that there is a degree of functional redundancy among the five clustered elongases, so that a simultaneous elimination of several (or all) of them would be required to feminize the male CHC profile more completely.

Another class of genes that may contribute to the sex-specific pheromone profiles of *D. prolongata* are fatty acid reductases (FARs). Enzymes in this large (17 genes in *D. melanogaster*) but poorly characterized gene family control the reduction of fatty acyl CoA to aldehydes and alcohols before they are converted to hydrocarbons by decarbonylation (Wicker-Thomas and Chertemps 2010; Qiu et al. 2012; Yew and Chung 2015). In moths, natural variation in FAR genes is responsible for the divergence of pheromone blends between populations and species (Lassance et al. 2010; Liénard et al. 2010), while in *Drosophila serrata*, *FAR2-B*, a recently duplicated ortholog of the *D. melanogaster CG17560*, explains sexually antagonistic variation in the relative amounts of short-chain and long-chain hydrocarbons (Rusuwa et al. 2022). In *D. prolongata*, the set of top candidate genes includes five FARs, including the ortholog of *CG17560*/*FAR2-B* (Fig. 4). Four of these genes are upregulated in *D. prolongata* males compared both to *D. carrolli* and to *D. prolongata* females, while the fifth, *CG17562*, is downregulated in *D. prolongata* males. The functional significance of these differences is unknown at this point. Since FAR-controlled decarboxylation competes with FAE-controlled elongation in determining whether precursors give rise to terminal products (pheromones) or to longer precursors with additional carbons (Fig. 8), one possible explanation is that the reduction of CG17562 in *D. prolongata* males facilitates elongation of 9T precursors to 9P/9H precursors instead of direct production of 9T. More generally, at least some FARs appear to be broad-spectrum enzymes: changes in, or disruption of, a single gene can alter the relative abundance of multiple long- and short-chain CHCs (Lassance et al. 2010; Liénard et al. 2010; Dembeck et al. 2015; Rusuwa et al. 2022). At the same time, FARs, like elongases, show some level of substrate specificity (Chung and Carroll 2015). It is possible that the FARs with male-biased and female-biased expression in *D. prolongata* have different substrate specificity (which may also vary among species), and that changes in their relative expression in males vs females contribute to the production of sex-specific pheromone blends. Genetic and biochemical evidence will be needed to test this hypothesis.

The exchange of signals involved in mating behavior is exceedingly complex. *eloF* is only one of the genes responsible for sex-specific pheromone profiles, while sex-specific pheromones are only one of the sensory cues underlying male-male and male-female communication. Putting together the complete puzzle will require not only identifying the missing pieces but also understanding how they interconnect.

## Materials and Methods

### Fly rearing and dissection for gene expression analysis

Flies were raised on standard cornmeal media and kept at room temperature (20-22°C) under natural light-dark cycle. For behavioral experiments and CRISPR gene editing, we used the reference genome strain of *D. prolongata* (Luecke et al. 2024), which was derived by four generations of full-sib inbreeding from the SaPa strain collected by Dr. H. Takamori (Luo et al. 2019). For RNA-seq and qPCR experiments, we used the BaVi strain of *D. prolongata* and the reference genome strain of *D. carrolli* (Luo et al. 2019). Both *D. prolongata* strains show strongly sexually dimorphic pheromone profiles, with females consistently distinguishable from males (F-type, (Luo et al. 2019)). For RNA-seq analysis, four biological replicates were prepared for each sex of *D. prolongata* and *D. carrolli*, resulting in 16 libraries. For qPCR experiments, three biological replicates were prepared for each species and sex. To obtain tissue samples enriched for oenocytes, we dissected the dorsal abdominal body wall (“cuticle fillet”) as described (Billeter et al. 2009); these samples are referred to as oenocyte dissections hereafter. Each biological replicate contained ten cuticle fillets. For head dissections, each biological replicate contained ten heads coming from the same tissue donors as the oenocyte dissections. All flies used for gene expression analysis were isolated as virgins and aged for seven days (*D. prolongata* and *D. carrolli*) or five days (transgenic *D. melanogaster*). Unless noted otherwise, tissues were dissected in chilled Shields and Sang M3 Insect Medium (Sigma-Aldrich, St. Louis, MO).

### RNA extraction

Total RNA was extracted from dissected fly tissues following the Trizol protocol (Ambion, Carlsbad, CA). Purified RNA was pelleted by isopropanol overnight at −20°C, washed by freshly made, pre-chilled 70% Ethanol (EtOH) 2 times, and dissolved in 20µl of DEPC-treated water (Ambion, Carlsbad, CA). To mitigate batch effects, flies were collected from the same food bottle, and flies collected on different dates were evenly distributed to each Trizol-containing tube. To ensure purity (A260/A280 > 1.9, A260/A230 > 1.5), isolated RNA was analyzed on a Nanodrop Spectrophotometer (ND-1000) using the software Nanodrop 1000 3.8.1. To ensure the integrity of RNA (2 sharp peaks of ribosomal RNA), gel electrophoresis was performed on an Agilent 2100 Bioanalyzer (Agilent) using RNA Nano Chips (Agilent). The concentration of RNA was determined using a Qubit 2.0 Fluorometer (Invitrogen) and Broad Range RNA Assay kit (Life Technologies). Finally, total RNA was DNase treated to remove carry-over genomic DNA following the rigorous DNA removal recommendations of the Turbo DNA-free kit (Invitrogen, Carlsbad, CA).

### Library construction, sequencing, and read mapping

cDNA libraries for RNA-seq were made using TruSeq Stranded RNA kit (Illumina, San Diego, CA) following the low throughput (LT) procedures in the user manual. 500ng of total RNA was used as starting material, and mRNA was selected by polyT enrichment. Reverse-transcribed cDNA was ligated with adapters uniquely barcoded for each library, followed by 11-cycle PCR amplification in an Applied Biosystems 2720 Thermal Cycler (Applied Biosystem, Waltham, MA). Thermocycling conditions were set as follows: 98°C for 30s, 11 cycles of 98°C for 10s, 60°C for 30s, 72°C for 30s, and a final 72°C for 5min. Amplified fragments were analyzed on an Agilent 2100 Bioanalyzer (Agilent) using High Sensitivity DNA Chips (Agilent). Unimodal fragment size distribution was consistently observed in all libraries, with a median fragment size of ∼300bp. The resulting cDNA libraries were initially quantified by Qubit 2.0 Fluorometer (Invitrogen) using the DNA High Sensitivity kit (Life Technologies) and further quantified by qPCR using the KAPA Library Quantification kit (Roche, Cape Town, South Africa). Barcoded cDNA libraries were subsequently pooled in equal molar ratios. To mitigate batch effects, all 16 libraries were prepared on two consecutive days, and within each day, two biological replicates of each group were processed.

Multiplexed libraries were sequenced on HiSeq4000 on PE-150 mode by Novogene (https://www.novogene.com). Reads were preprocessed (quality trimmed and deduplicated) by HTStream (Petersen et al. 2015). Cleaned reads were aligned to species-specific reference genomes (Kim et al. 2021; Luecke et al. 2024) using STAR (version 2.7.3a, (Dobin et al. 2013)) with the following flags:--sjdbOverhang 149--genomeSAindexNbases 13--genomeChrBinNbits 18 -- quantMode GeneCounts. Feature annotations were predicted by combining MAKER pipelines (Cantarel et al. 2008) and Liftoff (Shumate and Salzberg 2020) from existing genomic features of *D. melanogaster* (release 6.36) and *D. elegans* (Gnome annotation version 101). RNA-seq data and alignment statistics are summarized in Tables S7 and S8.

### Differential gene expression analysis

Paired-end fragments data were extracted from concordant read pairs. R packages “limma” (Ritchie et al. 2015) and “edgeR” (Robinson et al. 2010) were used to detect differentially expressed genes. To account for variations in sequence depth and RNA compositions between samples, sample-specific normalization factors were calculated using the Trimmed Mean of M-values (TMM) method (Robinson and Oshlack 2010). For each gene, counts per million (cpm) were computed. To remove genes with low expression, those with less than 2 cpm across all samples were excluded. Genes that could not be identified in both *D. prolongata* and *D. carrolli* (∼400 genes) were also excluded, resulting in a set of 9143 genes. To obtain normalized expression data, TMM-based normalization factors were applied, followed by calculating the log_2_ cpm of these genes. To account for the mean-variance trend associated with each gene (e.g., genes with low mean expression tend to have larger variances), voom transformation was applied to estimate the weights of genes (Law et al. 2014). These weights were then used to fit a weighted linear model on normalized expression data (log_2_ cpm), using groups (sex x species) as predictors.

Three one-way comparisons were defined to identify candidate genes for pheromone divergence: (1) contrasting *D. prolongata* males with *D. prolongata* females; (2) contrasting *D. prolongata* males with *D. carrolli* males and (3) comparing the magnitude of the male-female difference between *D. prolongata* and *D. carrolli*. This last test helps to identify changes that result either from differences in the magnitude of sexual dimorphism in each species (i.e., which one shows greater male-female difference?) or the direction of sexual dimorphism in each species (i.e., is the direction of sex-biasedness the same?). Linear contrasts were made for each one-way comparison. To account for variance that comes from random factors (not low expression), we used empirical Bayes smoothing (Smyth 2004). To adjust for multiple testing, false discovery rate (FDR) corrections were applied to raw p values (Benjamini and Hochberg 1995), and significant differentially expressed genes (DEGs) were reported with FDR < 0.05 (Supplemental files 1-6).

To be considered candidates, genes had to be reported as significantly different in all three one-way comparisons (referred to as the three-way comparison hereafter). P values for the three-way comparison were constructed by taking the maximum of p values from the one-way comparisons, resulting in 53 candidate genes (p < 0.05, Fig. 2). To visualize the expression profiles of top candidates, hierarchical clustering was performed on their standardized expression levels. Euclidean distances were calculated, and UPGMA (i.e., average linkage) was used for the hierarchical clustering. The same metrics were used to cluster both samples and genes. Volcano plots (Fig. 2) were generated by the “ggplot2” package, and heatmaps (Fig. 3) by the “stats” package, both of which were subsequently polished by the Inkscape software (https://inkscape.org).

### Gene Ontology (GO) enrichment analysis

GO enrichment analysis was performed for DEGs identified from each one-way comparison (i.e., male vs. female in *D. prolongata*, males of *D. prolongata* vs. *D. carrolli*, and the magnitude and direction of sexual dimorphism in *D. prolongata* vs. *D. carrolli*). To determine whether these DEGs were expressed in oenocytes ancestrally, we consulted the single-cell Fly Cell Atlas data from *D. melanogaster* (Li et al. 2021). ∼3000 cells annotated as adult oenocytes (FBbt:00003185) were retrieved from the “10x relaxed dataset” on SCope (https://scope.aertslab.org/#/FlyCellAtlas/FlyCellAtlas%2Fr_fca_biohub_oenocyte_10x.loom/ge <u>ne</u>). Oenocyte expressors were defined as genes that were detected in at least ten cells (i.e., at least one transcript in each of 10 cells) and had at least 50 cumulative read counts in either female or male samples. This produced a list of ∼6200 oenocyte expressors (Supplemental file 7).

We annotated GO terms with the following criteria. For genes annotated with *D. melanogaster* CG/CR numbers, their GO annotations from the R Bioconductor package “org.Dm.eg.db” (version 3.14.0) were used. This provided GO annotation for 7741 genes. For genes annotated with *D. melanogaster* CG/CR numbers that did not have GO annotations in “org.Dm.eg.db”, their GO annotations were retrieved from orthoDB (version 10.1, https://www.orthodb.org). This provided GO annotation for another 314 genes. For genes annotated with *D. elegans* LOC numbers, their GO annotations were retrieved from orthoDB (version 10.1). This provided GO annotation for additional 529 genes. In this way, a gene-GO map was built to cover 93.8% (8584) of the entire gene set used in the RNA-seq analysis.

R Bioconductor package “TopGO” (version 2.46.0, (Alexa and Rahnenfuhrer 2022)) was used to perform enrichment analysis on Biological Processes GO terms. To control for potential artifacts due to small GO categories, those with <10 associated genes were excluded, as recommended by the program. To account for the tree topology between GO terms, the modified elimination algorithm weight01 (Alexa et al. 2006) was used. By doing this, parent nodes of significant child nodes are less likely to be annotated unless they contain substantially more significant genes not covered in their children. This also helps balance a low false positive rate and a high recall rate. P values were obtained from Fisher exact test (Drăghici et al. 2006) and the Kolmogorov-Smirnov test (Ackermann and Strimmer 2009). No p-value adjustment was performed as recommended by the program developer (Alexa and Rahnenfuhrer 2022). Instead, candidate GO terms were defined as those with p values <0.05 for both Fisher and Kolmogorov-Smirnov tests.

### Quantitative polymerase chain reaction (qPCR) analysis of gene expression

To quantify the expression of endogenous *eloF* in *D. prolongata* and *D. carrolli*, two-step qPCR was performed (i.e., cDNA synthesis followed by separate qPCR analysis). cDNA was synthesized from 1µg of DNase-treated total RNA using Superscript III (Invitrogen) following kit recommendations. To prime the reverse transcription reaction, a volume ratio of 1:1 random hexamer (Invitrogen) and oligo dT (Invitrogen) was used. Reactions were performed in a thermal cycler (Applied Biosciences) with the following conditions: initial incubation at 25°C for 5min, reverse transcription at 50°C for 50min, and enzyme inactivation at 70°C for 15min. The resulting single-stranded cDNA was diluted by a factor of 100 and stored at −20°C prior to qPCR.

Green Fluorescence Protein (*GFP*) expression in reporter assays (see “Design of reporter constructs” below) was quantified using one-step RT-qPCR (i.e., combining reverse transcription and PCR amplification in the same tube). This was done because the reporter GFP is a single-exon gene (Fig. S12), so that amplification of *GFP* transcripts could be confounded by even trace amounts of GFP DNA. The entire experiment was conducted on a clean bench free of DNA contaminants, and PCR-grade water (IBI Scientific, Dubuque, IA) was used to assemble the reaction. As GFP expression was preliminarily found to be low, 300ng DNase-treated RNA was used for each reaction. To prepare no-reverse-transcriptase (NRT) controls, total RNA samples from 3 biological replicates were pooled in equal mass ratios and received the same treatment. All NRT controls showed Ct >35 (Table S2), indicating sufficient removal of genomic DNA.

qPCR reactions were assembled using SsoAdvanced SYBR Green PCR Supermix kit (Bio-Rad, Hercules, CA) on Bio-Rad CFX96 Real-time PCR system. Amplification was performed in 10µl total volumes with a 4µl template (1:100 diluted cDNA or 300ng DNase treated total RNA) and 100nM of each primer in a 96-well optical plate (Bio-Rad). Melt-curve analysis was performed on the PCR products to assess the presence of unintended products. Thermocycling conditions are set as follows. For qPCR: initial denaturing at 95°C for 1min, followed by 40 cycles of 95°C for 10s and 60°C for 10s. For RT-qPCR: Reverse transcription at 50°C for 10min, initial denaturing at 95°C for 1min, followed by 40 cycles of 95°C for 10s and 60°C for 10s. For melt-curve analysis: from 65°C to 95°C at an increment of 0.5°C, hold 5s for each temperature step. For both one-step or two-step qPCR, three biological replicates were made for each group, and each reaction was technically replicated three times to obtain an average Ct value. Technical reproducibility was consistent with standard deviations within 0.5 Ct (Scott Adams 2007). *Ribosomal protein L32* (*Rpl32*) was chosen as a reference gene for its stable expression level (Ponton et al. 2011). Standard curves were built to determine primer amplification performance (e.g., primer efficiency) (Fig. S14). Specifically, qPCR was performed on a diluted DNA template covering at least 6 log range. qPCR amplification metrics were determined for each gene with the slope of a linear regression model (Pfaffl 2001). Relative efficiencies were calculated according to the equation: E = (dilution factor − 1/slope − 1) x 100%. Primer sequences, design considerations, coefficient of determination (R^2^), and amplification efficiencies are summarized in Table S9. As all primers had near-perfect amplification efficiency, the ΔΔCt method (Livak and Schmittgen 2001) was used for the relative quantification of genes of interest (*eloF* and *GFP*).

To model normalized expression levels, a two-way ANOVA with interaction effects between genotypes (species) and sex was used, similar to the section “Statistical analysis of mutant CHC profiles.” Statistical significance for genotype, sex, and their interactions was tested by comparing the full model with a reduced model after dropping the term of interest. Type III variance partitioning was used, and Tukey’s method was used to determine which construct has a significantly higher expression level.

### Cuticular lipids extraction

Virgin *D. prolongata* with wild-type or mutant *eloF* were individually isolated within 12 hours after eclosion. After aging for 7 days, individual flies were frozen at −20°C, transferred to pure hexane (Sigma-Aldrich), soaked for 5 min at room temperature, and vortexed for 30 seconds. To ensure complete CHC extraction, 40µl of pure hexane was used for females and 80µl for males, due to the large size difference. Crude extracts were air-dried overnight and stored at 4°C before GC-MS analysis. To quantify the absolute amount of each analyte, hexane containing 10ng/µl n-heneicosane (nC26, Sigma-Aldrich) and 10ng/µl n-triacotane (nC30, Sigma-Aldrich) as alkane standards was used to resolubilize crude extracts. 40µl of this solvent was used for females and 80µl for males.

### Gas chromatography (GC) and mass spectrometry (MS) analyses

GC-MS analysis was performed as in Luo et al., 2019 with the following modifications. The oven temperature was programmed to first ramp from 160°C to 280°C at a rate of 8C/min, hold at 280°C for 1 min, and increase from 280°C to 315°C at a rate of 15°C /min, followed by a final 1min hold at 315°C. The flow rate of carrier gas (helium) was optimized to 1ml/min. Individual chromatographic peaks were first called using the built− in ChemStation integrator of MSD ChemStation Enhanced Data Analysis Software vF.01.00 (Agilent Technologies, Santa Clara, CA), with initial peak width of 0.030 and an initial threshold of 16. Manual adjustments were made to include minor peaks and deconvolute overlapping peaks. Analytes were then identified (Table S3) and quantified as described previously (Luo et al. 2019). Briefly, all CHCs were normalized by alkane standards and scaled in units of nanograms per individual fly.

### Female perfuming experiments

Synthetic (Z)-9-Pentacosene (9P) was purchased from Cayman Chemical (Ann Arbor, MI), and (Z)-9-Heptacosene (9H) was kindly provided by Dr. Jocelyn Miller (University of California, Riverside). To prepare perfuming vials, batches of hexane solutions containing 9P (9P treatment), 9H (9H treatment), or nothing (control) were added to and air-dried inside 2mL glass vials (Agilent Technologies, #5182-0715, Santa Clara, CA). 50µg 9P and 10µg 9H were used to ensure consistent and biologically reasonable perfuming (Fig. 1). Flies were perfumed according to a modified protocol of Billeter et al., 2009, briefly summarized as follows. Groups of eight virgin, 7-day old female flies were placed inside clean 2mL glass vials (Agilent Technologies) and vortexed on medium speed to capture the CHC profile before the perfuming study. To perfume with synthetic hydrocarbons, the same group of 8 flies was subsequently transferred to the perfuming vial prepared as described above and vortexed intermittently. 4 groups of flies were prepared per day, resulting in a total of 8 groups containing 64 individuals. Perfumed flies were allowed to recover for 3 hours and divided randomly into two equal groups, with one group of four used immediately for assessing male-female interactions (assay group) and the other saved for confirming the transfer of desired CHCs (validation group). 200µl of pure hexane was used to extract CHCs from the validation group. Both pre-and post-perfuming crude extracts were resolubilized with alkane standards as described above (see “cuticular lipids extraction”), except that 20µl was used for pre-perfuming samples (N = 8) and 100µl for post-perfumed samples (N = 8). To quantify changes in the CHCs of interest, all samples were analyzed by GC-MS as described above.

### Behavioral assays

Cameras and the behavior arena were set up as previously described (Toyoshima and Matsuo 2023). For male-female interaction experiments, a single virgin female was paired with a single virgin male inside a food podium. For male-male interaction experiments, two virgin males of the same genotype were placed together without any females being present. For misdirected courtship experiments, a pair consisting of one wild-type male and one wild-type female was combined with a single decapitated male, whose genotype was either wild-type or *eloF[-]*. The genotypes and numbers of individuals are reported separately for each experiment in the figure legends. The flies were videotaped for 1 hour, and binary metrics of previously characterized behaviors, including “encounter,” “threatening,” “courtship,” “leg vibration,” “wing vibration”, “copulation”, and “boxing” were scored from the video recordings (Setoguchi et al. 2014; Kudo et al. 2015).

To test whether male-biased hydrocarbons are transferred to females during mating, 6-8 day-old virgin males and females from the reference genome strain were placed together in single pairs (n = 16). Behavior was observed in the morning for 1 hour to determine whether mating occurred. To test for quantitative changes in CHC profiles, whole-body pheromone extractions were performed on mated and unmated females on the same day after the observation concluded. Socially naive females and males were included as controls.

### Statistical analysis of behavioral changes

A logistic regression model was used for each binary behavior (e.g., courtship) with the genotype as the only explanatory variable, and an ordinary linear regression model for each continuous behavior (e.g., copulation duration). Z-tests were performed on coefficients from logistic regression to determine the p-value for each comparison between *eloF[-]* mutant and wild-type alleles. t-tests were performed on coefficients from ordinary linear regression.

### Statistical estimation of hydrocarbon transfer

Hydrocarbon transfer was estimated from the increase in the abundance of the analyte of interest (9P or 9H) after perfuming. For each group of 4 females used in behavioral tests, we created a parallel group of 4 females that were subjected to the same perfuming procedure but were not used in behavioral assays (see “female perfuming preparation” above). This replicate group was used to validate the transfer of desired CHCs, in conjunction with pheromones extracted from the same group before perfuming. Instead of simply taking the difference in 9P (or 9H) abundance before and after perfuming, we calibrated the post-perfuming abundance of 9P (or 9H) by a method analogous to standard curves to mitigate technical variation as follows. For each perfuming group of eight flies, a calibration curve was made by regressing the post-perfuming on the pre-perfuming abundances of all CHCs except those modified in treatment (e.g., leaving out 9P in 9P treatment). A general agreement was found between pre-post pairs of endogenous CHCs, with coefficients of determination (R^2^) ranging from 0.85 to 0.98 (Fig. S15). Leveraging this property, expected post-perfuming abundance of 9P (or 9H) if no synthetic 9P (or 9H) were transferred (i.e., the “counterfactual” abundance) was then predicted based on the sample-specific standard curve. Likewise, 95% confidence intervals were constructed around the expected abundance. Finally, the hydrocarbon transfer was estimated as the difference between the observed and “counterfactual” abundance.

### Gene editing by CRISPR/Cas9 mutagenesis

To create null mutants for *elongase F* (*eloF*) in *D. prolongata*, two guide RNAs were designed that target its first exon (Fig. 6). Guide RNA sequences were as follows: gRNA43: 5’-TCTGCTATTTGTCCTCAAGGTGG-3’ and gRNA84: 5’-AGAGTACCCAGAGCAACCCATGG-3’. Embryo injection and mutation screening were conducted as described (Takau and Matsuo 2022). Deletion of sequences between two guide RNAs was confirmed by Sanger sequencing. Two mutant strains were obtained: one with a frameshift mutation resulting in an early stop codon, and the other with a 45bp in-frame deletion resulting in the loss of 15 amino acids (Fig. 6).

### Statistical analysis on mutant CHC profiles

To determine the effects of *eloF* on pheromone production in *D. prolongata*, we examined the CHC profiles of both homozygous *eloF* mutant strains generated by CRISPR/Cas9 mutagenesis. The reference genome strain, in which these mutants were induced, was used as the control. Multivariate and univariate analyses were performed on the absolute quantity (on a logarithmic scale) of 18 consensus CHCs that are shared between sexes and collectively account for >98% of total CHC abundance. Prior to principal component analysis (PCA), CHC abundances were centered to zero means but not standardized to unit variance, so PCA was conducted on the sample covariance matrix. In the PCA scatter plot, 95% confidence regions for each group (genotype x sex) were estimated assuming underlying bivariate t-distributions. To determine whether (1) pheromone profiles of wild-type and *eloF* mutant flies were significantly different and (2) whether mutation effects differed between sexes, we used two-way ANOVA models with interaction effects between sex and genotype, followed by Tukey’s range test for all pairwise comparisons. The ANOVA model was specified as follows: Log(abundance) ∼ sex + genotype + sex * genotype. Data management (R package suite “tidyverse”) and statistical modeling (R packages “car,” “multcomp,” “lsmeans”) were conducted by in-house R scripts (R Core Team 2022), with plots generated by the “ggplot2” package and subsequently polished by the Inkscape software (version 0.92.4, https://inkscape.org).

### Comparative sequence analysis

Sequences surrounding the *eloF* locus (∼2kb) were extracted from reference genomes of each species (Kim et al. 2021; Luecke et al. 2024). To study sequence evolution, multiple alignments of DNA sequences were conducted using Clustal Omega (version 1.2.2 (Sievers et al. 2011)) using the default parameters. To examine sequence divergence of *eloF* orthologs, single nucleotide variants (SNVs) in the coding sequence (CDS) of *eloF* were called (Table S5) by manual inspection of RNA-seq reads that mapped to a nearby region using the software IGV (version 2.4.11, Broad Institute). Open reading frames (ORFs) were predicted based on the standard genetic code and required a minimum of 400 base pairs. To identify the genetic nature of “*honghaier*,” a putative transposable element, and its associated ORF, the web application BLASTn (version 2.13.0+) was used to search against all NCBI databases and the database of known transposable elements Dfam (Storer et al. 2021). To visualize the phylogenetic distribution of honghaier and associated ORF (Table S6), local standalone blastn databases were made from genome assemblies, and command-line-based BLASTn (version 2.2.31+) was used.

To assess the sequence complexity of the honghaier insertion, a preliminary dot plot (not shown) was made using the EMBOSS (version 6.5.7) tool dotmatcher, with a word size of 10. *De novo* motif discovery was subsequently made to identify the repeating units using MEME-suite (https://meme-suite.org) software MEME (version 5.3.2, (Bailey and Elkan 1994)). The following command-line flags were used: “-dna-mod anr-nmotifs 3-revcomp”. The AT content was estimated by averaging the occurrence of adenosine (A) and thymine (T) in a window of 50 bp. Unless otherwise noted, sequence analysis was conducted in Geneious Prime (version 2021.0.3, Biomatters, www.geneious.com).

### Transcription factor (TF) binding motif analysis

Since the exact motif sequences that activate gene expression in adult oenocytes are largely unknown, we used de novo prediction to identify TF-binding motifs that are enriched in adult oenocytes. A list of genes annotated as being differentially expressed in adult oenocytes over other tissues (referred to as oenocyte markers hereafter) in *D. melanogaster* was downloaded from single-cell Fly Cell Atlas (“10X relaxed dataset”, (Li et al. 2021)). Marker genes were stringently filtered using log fold change cutoff > 1 and a p-value cutoff of 1e-10. Using R Bioconductor packages “org.Dm.eg.db” (feature annotation database, version 3.14.0), “TxDb.Dmelanogaster.UCSC.dm6.ensGene” (transcript database, version 3.12.0), and “BSgenome.Dmelanogaster.UCSC.dm6” (genome database, version 1.4.1), genes were further filtered by the following criteria. Oenocyte marker genes must (1) have a matching FlyBase unique gene identifier and (2) map to chromosome X, 2, or 3. This resulted in a final set of 956 oenocyte-enriched markers (Supplemental file 8). This list included the previously reported oenocyte markers *desaturase F* (*desatF*, (Chertemps et al. 2006)) and *elongase F* (*eloF*, (Chertemps et al. 2007)). For each oenocyte marker, up to 1kb of the upstream promoter region was extracted for motif enrichment analysis.

Motif enrichment analysis was performed on the retrieved upstream sequences (Supplemental file 9) by Meme Suite software AME (version 5.3.2, (McLeay and Bailey 2010)) using the following command line flags:-control--shuffle---scoring avg-method fisher. The iDMMPMM motif database downloaded from Meme Suite provided 39 known motifs with well-supported DNase-I footprint evidence (Kulakovskiy and Makeev 2009). We observed significant enrichment for binding motifs associated with the TFs *bric-a-brac* (*bab1*, p = 1.75e-60) and *Mothers against dpp* (*Mad,* p = 2.14e-21). Both these genes are expressed in the adult oenocytes of *D. melanogaster* (Supplemental file 7). Other candidates were not considered because their p values were several orders higher than the top 2 candidates.

Using *bab1* and *Mad* as candidate motifs that may underlie oenocyte development, motif occurrence analysis was performed on non-protein-coding regions of *eloF* across 5 species in the *rhopaloa* subgroup using Meme Suite software FIMO (version 5.3.2, (Grant et al. 2011)). The following command line flags were used: “--parse-genomic-coord,--thresh 0.001”. As no matches corresponding to *Mad* were found, only *bab1* binding motifs were reported (Fig. S9). In addition to tissue-specific motifs, individual motif occurrence analysis was performed on sex-related motifs by FIMO. The binding motifs of the *doublesex* (*dsx*) TF were retrieved from Shirangi et al. (Shirangi et al. 2009), FlyReg (Bergman et al. 2005), Fly Factor Survey (https://mccb.umassmed.edu/ffs), and JASPAR (9th release, https://jaspar.genereg.net). As no match was found for the motif reported by Shirangi et al., sex motifs included three targets: *dsx* from JASPAR and *dsx-F* and *dsx-M* from FlyReg (where both proteins have identical binding sequences).

### Design of reporter constructs

We generated transgenic *D. melanogaster* strains that carried orthologous *eloF* sequences from *D. prolongata* and *D. carrolli*. “Long” constructs were designed to cover the entire *eloF* locus and its whole flanking region (between the flanking genes CG16904 on the left and CG8534 on the right): “Dpro *eloF* WT^(l)^” with the allele from *D. prolongata* and “Dcar *eloF* WT^(l)^” with the allele from *D. carrolli* (Fig. S12). The downstream region of *eloF* contains a putative transposable element (TE) insertion, which we named “*honghaier*”, in *D. prolongata* but not in *D. carrolli* or any other species at this location. Two additional constructs were therefore produced by a TE swap: one engineered allele had *honghaier* removed from the *D. prolongata* allele (“Dpro *eloF* WT^(l)^ - TE”), and the other had *honghaier* inserted into the *D. carrolli* allele (“Dcar *eloF* WT^(l)^ + TE,” Fig. S12). In addition, “short” reporter constructs were designed with the DNA sequences of the downstream region of *eloF* (between CG16904 on the left and *eloF* on the right; note that the two genes are transcribed in head-to-head orientation). Similar to the “long” constructs, two of the short constructs were wild-type alleles of each species, “Dpro *eloF* WT^(s)^” and “Dcar *eloF* WT^(s)^”, while the other two were produced by the *honghaier* swap: “Dpro *eloF* WT^(s)^ - TE” and “Dcar *eloF* WT^(s)^ + TE,” Fig. S12.

To clone the reporter sequences, DNA fragments were amplified by SeqAmp (Takara Bio, San Jose, CA), a proofreading DNA polymerase (See Table S9 for primers used in this cloning experiment). PCR-amplified DNA fragments were first Gibson-cloned into linearized pCR8 vectors (Invitrogen, Carlsbad, CA) using Gibson Assembly Master Mix (New England Biolabs, Ipswich, MA) according to kit recommendations. We then conducted a Gateway reaction to transfer the DNA inserts into the destination vector pGreenFriend (Fig. S12, (Miller et al. 2014)) by Gateway recombination reaction using LR Clonase II Enzyme mix (Invitrogen). The pGreenFriend vector has a GFP reporter driven by the Drosophila Synthetic Core Promoter (Fig. S12). Final constructs were bulk-purified using a QIAGEN midi-prep kit (QIAGEN, Redwood City, CA) and confirmed by Sanger sequencing (McLab Sequencing, San Francisco, CA). Chemically competent *E. coli* strain NEB5alpha H2987 (New England Biolabs) was used for transformation.

### Transgenic strains

The pGreenFriend vector has a single attB site that allows it to integrate into attP anchor sites in the *D. melanogaster* genome (Fig. S12). 30µg of purified plasmids were sent to BestGene (https://www.thebestgene.com) for embryo injection. The genotype of injected flies was *y^1^ w^67c23^; P{CaryP}attP40*, with the attp40 landing site on the second chromosome (Markstein et al. 2008). Transformed G0 flies were crossed to *yw* flies, and the resulting G1 progeny were genotyped to verify successful integration. Heterozygous flies carrying attP insertion (attP40*) were selected based on orange eye color. Confirmed insertions were balanced and flies homozygous for the attP40* site with the reporter insertion were selected from these balanced strains and used for antibody staining and RT-qPCR.

### Tissue dissection and antibody staining

Homozygous transgenic flies were isolated as virgins and the dorsal abdominal body wall was dissected in 1x Tris-NaCl-Triton (TNT) buffer (100mM Tris pH7.5, 300mM NaCl, 0.5% Triton-X). Flies with the *yw* genotype were used as negative control to account for oenocyte autofluorescence. A standard fixation protocol was adopted (Tanaka et al. 2009). Tissues were pooled and fixed in a fixation buffer (100mM Tris pH7.5, 300mL NaCl, 4% paraformaldehyde (Electron Microscopy Sciences, Hatfield, PA)) for 20min on a rotation platform set to a gentle speed at a frequency of 0.33Hz. Fixed tissue was then washed in 1mL 1xTNT for 15min three times. Post-wash tissues were stored in 1mL fresh 1xTNT at 4°C until antibody staining.

For GFP staining, fixed tissues were first transferred to 3×3 dissection plates and washed with 300µl 1xPhosphate-Buffered Saline with Triton-X 100 (PBST buffer) for 15 min. To reduce non-specific antibody binding, washed tissues were blocked in 180µl of 5% normal goat serum (Jackson Immunoresearch, West Grove, PA) for 30min, followed by three times 1x PBST wash (300 µl, 15min each). Tissues were stained using 300µl of primary antibody at 4°C overnight and washed three times in 300µl 1x PBST on the following day. Immediately after primary staining, tissues were stained using 300µl of secondary antibody at 4°C for 1h and washed three times in 300µl 1x PBST (15min each). Stained tissues were stored in 1mL 1xPBST at 4°C and covered with aluminum. All antibody staining steps were performed in the dark and incubated on a nutator set to gentle speed. The ingredients of buffers used are summarized as follows. 1xPBST buffer was prepared by adding Triton-X to 1xPhosphate-Buffered Saline (1×1xPBS, Corning, Manassas, VA) to a final concentration of 0.4% (v2v). The blocking solution was freshly made by adding goat serum (Jackson Immunoresearch) in 1xPBST to a final concentration of 5% (v2v) and stored at 4°C upon use. Staining solutions were freshly made by diluting primary (Chicken-anti-GFP, Invitrogen) or secondary antibodies (goat anti-chicken-AF488, Jackson Immunoresearch) in 1x blocking solution to a final concentration of 1:200. At least four dissected cuticle filets were studied for each short construct.

To facilitate mounting, fully stained dorsal cuticles were flattened by trimming to an approximately rectangular shape. Under the dissecting microscope, the margins of the A1 segment and A6/A7 segments were first removed, and the lateral sides of the remaining segments were trimmed by fine scissors. After flattening, the dorsal cuticle was placed on a cover slide (22×22 mm, thickness 1.5, Corning) with the interior facing up. A 20µl of antifade FluromountG reagent (Electron Microscopy Sciences) was added and spread evenly to reduce the formation of air bubbles. The mounting slide (3’’ x 1’’ x 1mm, Fisher Scientific, Pittsburgh, PA) was placed on top of the cover slide to finish preparation. Mounted slides were stored in a slide binder at 4°C before imaging.

### Confocal microscopy

Mounted tissues were imaged using a Leica SP8 confocal microscope. The 488 nm laser was used to visualize GFP. Images were taken every 5 µm using a 20X objective and digital zoom. Images were further processed in Fiji ImageJ (Schindelin et al. 2012) to project everything on a z-stack with maximum intensity.

## Acknowledgements

We thank Dr. Jocelyn Miller (University of California, Riverside) for providing synthetic 9-heptacosene, Logan Blair and David Luecke for advice on experiments and comments on the manuscript, and Majken Horton, Madison Hypes, Yingxin Su, and Olga Barmina for technical assistance. This work was supported by NIH grant R35 GM122592 to AK, JSPS grants No. 18H02507 and No. 24K01767 to TM, funds from the UC Davis Center for Population Biology to YL, and David and Lucile Packard Foundation grant 2014-40378 to SRR. Confocal imaging was made possible by the MCB Light Microscopy Core Facility at UC Davis, funded by the NIH grant S10OD026702, with training and support by Thomas Wilkop.

## Supplemental Information

Supplemental Figures S1 – S15

Supplemental Tables S1 – S9

**Figure S1.**
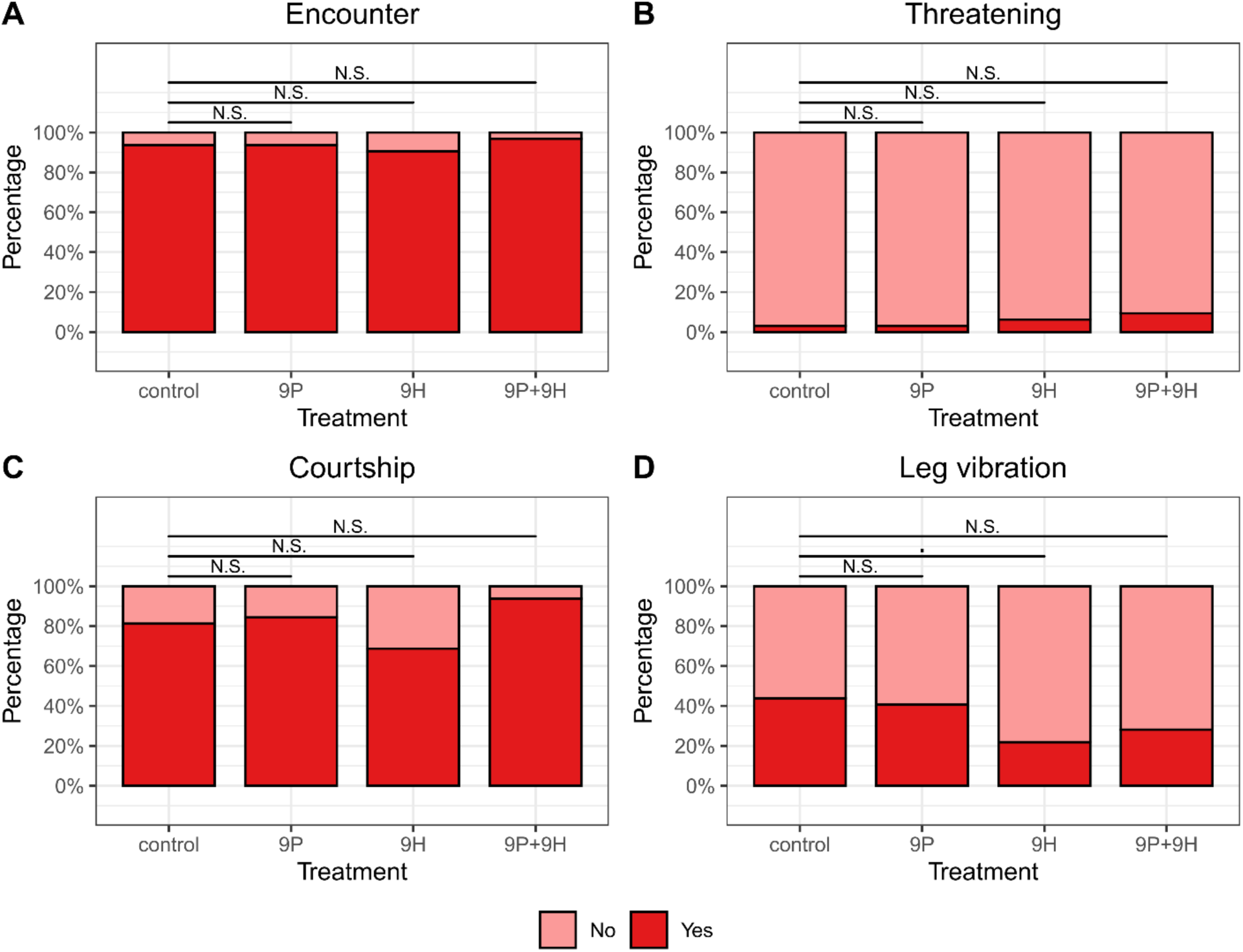
Variable effect of perfuming on male-female interactions. Stacked bar plots showing success rates of (A) Encounter, (B) Threatening, (C) Courtship, and (D) Leg vibration across three perfuming conditions (N = 32 for each treatment). N.S., nonsignificant results based on comparison between treatment and control in a logistic regression model. P values are as follows: *** p < 0.001, ** p < 0.01, *, p < 0.05, p < 0.1.

**Figure S2.**
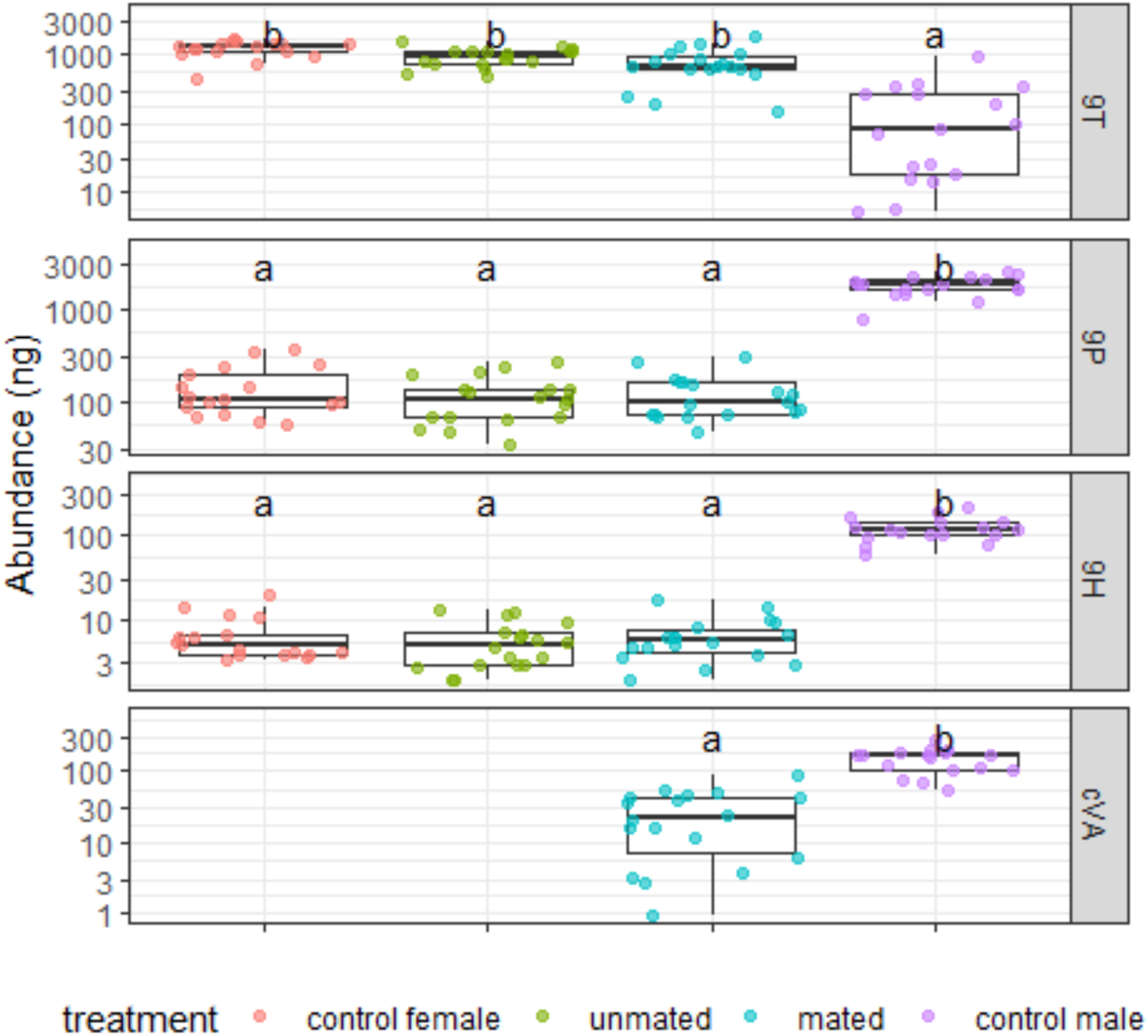
Male-biased long-chain CHCs are not transferred to females during mating. Boxplots showing the abundance of 9T, 9P and 9H, with cis-vaccenyl-acetate (cVA) as a positive control. Shown are control wild-type females, WT females that did not mate with a WT male, WT females that mated with a WT male, and control WT males. Pheromone abundance is measured in nanograms per fly and shown on log10 scale. Overlayed jitter points are samples of each sex * mating status combination, color-coded by genotype. Significance results of all pairwise comparisons (Tukey HSD test followed by significant omnibus ANOVA F-tests) are summarized in the format of compact letter display (using R packages “multicomp” and “lsmeans”).

**Figure S3A.**
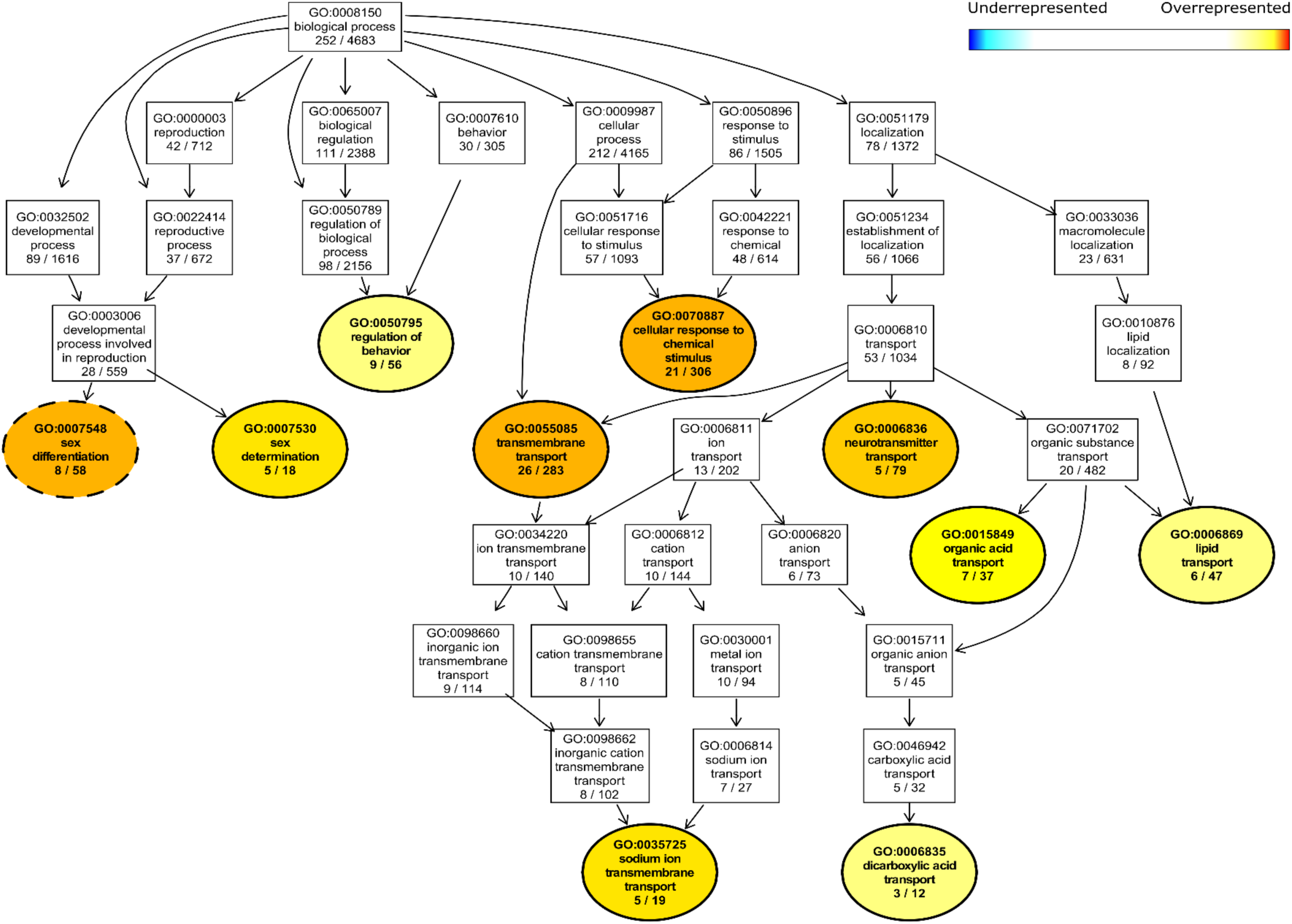
Sex differentiation and substance transport show differential enrichment between males and females of *D. prolongata*. Directed Acyclic Graph (DAG) of significant GO terms and their parent terms in biological processes. Significant (p < 0.05) and non-significant GO terms are color-coded and represented by ellipses and rectangular boxes, respectively. Significant GO terms can be underrepresented (blue) or overrepresented (red) based on Fisher’s exact test. Arrows indicate hierarchical relationships. GO terms at the same hierarchical level are placed at the same vertical position. Significant GO terms that are also enriched between males and females of *D. carrolli* have dashed borders.

**Figure S3B.**
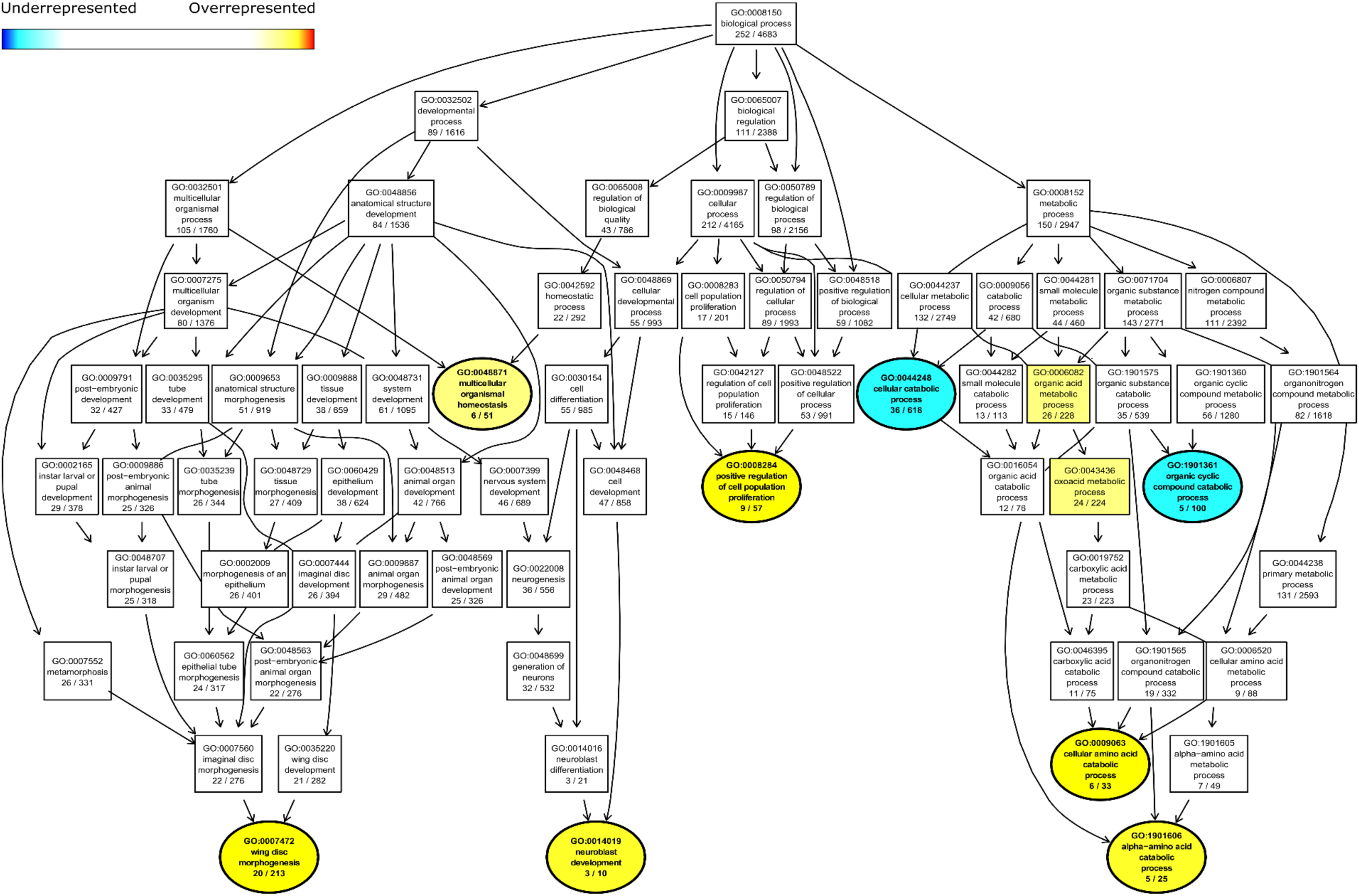
Terminal processes of amino acid metabolism show differential enrichment between males and females of *D. prolongata*. Directed Acyclic Graph (DAG) of significant GO terms and their parent terms in biological processes. Significant (p < 0.05) and non-significant GO terms are color-coded and represented by ellipses and rectangular boxes, respectively. Significant GO terms can be underrepresented (blue) or overrepresented (red) based on Fisher’s exact test. Arrows indicate hierarchical relationships. GO terms at the same hierarchical level are placed at the same vertical position. Significant GO terms that are also enriched between males and females of *D. carrolli* have dashed borders.

**Figure S4A.**
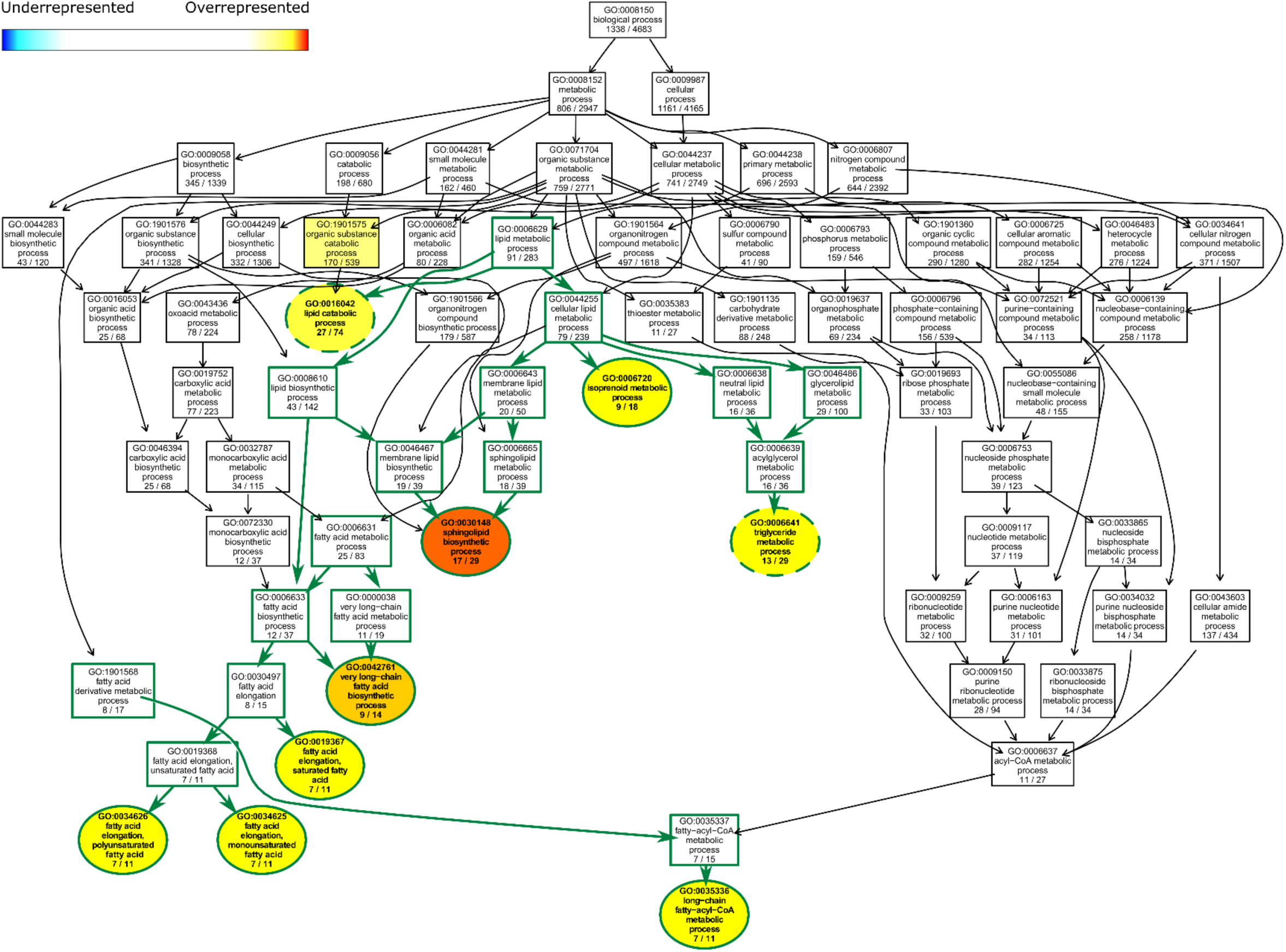
Terminal lipid metabolism processes show differential enrichment between males of *D. prolongata* and *D. carrolli*. Directed Acyclic Graph (DAG) of significant GO terms and their parent terms in biological processes. Significant (p < 0.05) and non-significant GO terms are color-coded and represented by ellipses and rectangular boxes, respectively. Significant GO terms can be underrepresented (blue) or overrepresented (red) based on Fisher’s exact test. Arrows indicate hierarchical relationships. GO terms at the same level are positioned at the same vertical position. GO terms under the lipid metabolic process (GO:0006629) are connected by green arrows and have green borders. Significant GO terms that are also enriched between females of *D. prolongata* and *D. carrolli* have dashed borders.

**Figure S4B.**
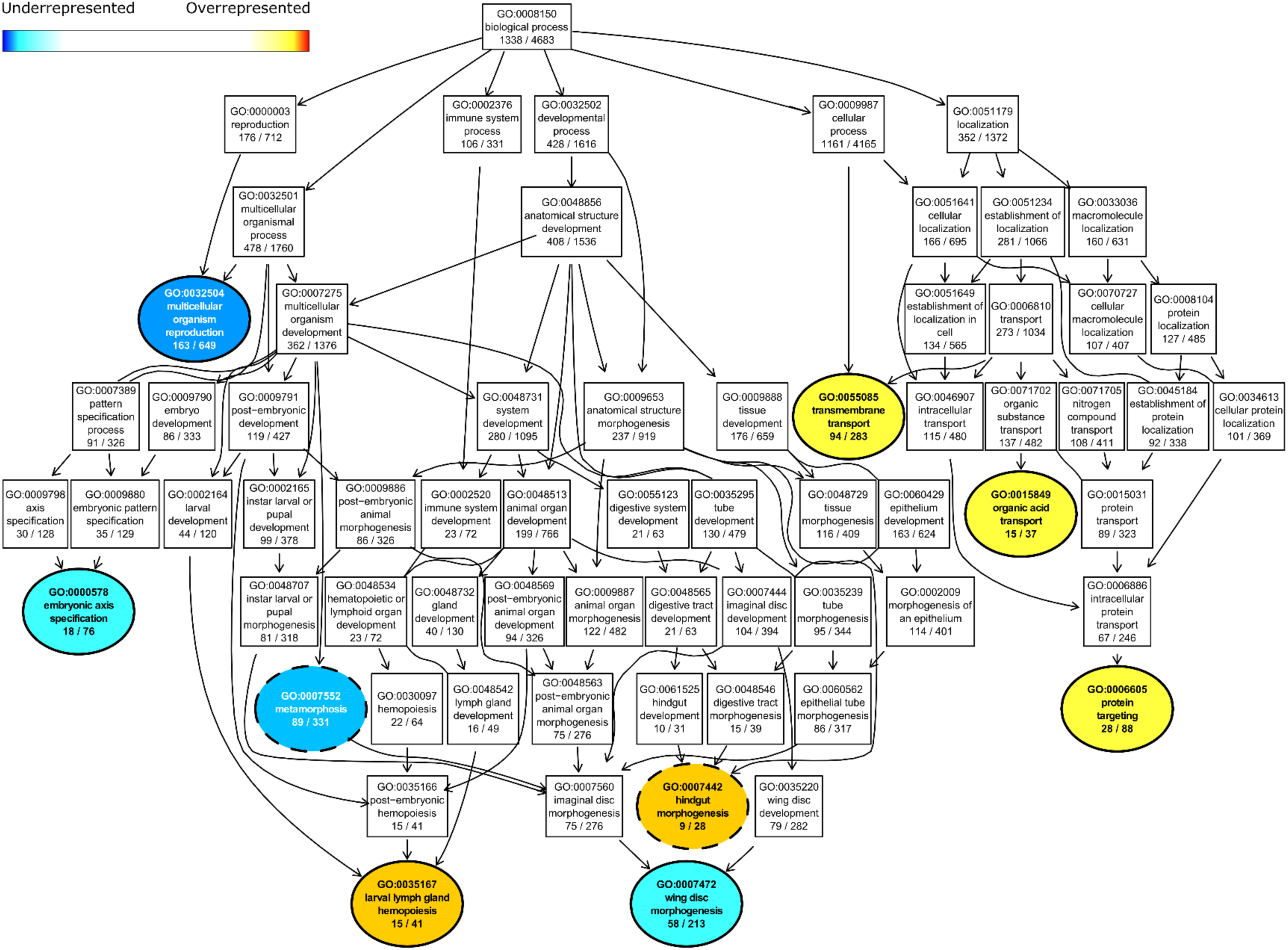
Substance transport, development, and reproduction show differential enrichment between males of *D. prolongata* and *D. carrolli*. Directed Acyclic Graph (DAG) of significant GO terms and their parent terms in biological processes. Significant (p < 0.05) and non-significant GO terms are color-coded and represented by ellipses and rectangular boxes, respectively. Significant GO terms can be underrepresented (blue) or overrepresented (red) based on Fisher’s exact test. Arrows indicate hierarchical relationships. GO terms at the same level are positioned at the same vertical position. Significant GO terms that are also enriched females of *D. prolongata* and *D. carrolli* have dashed borders.

**Figure S4C.**
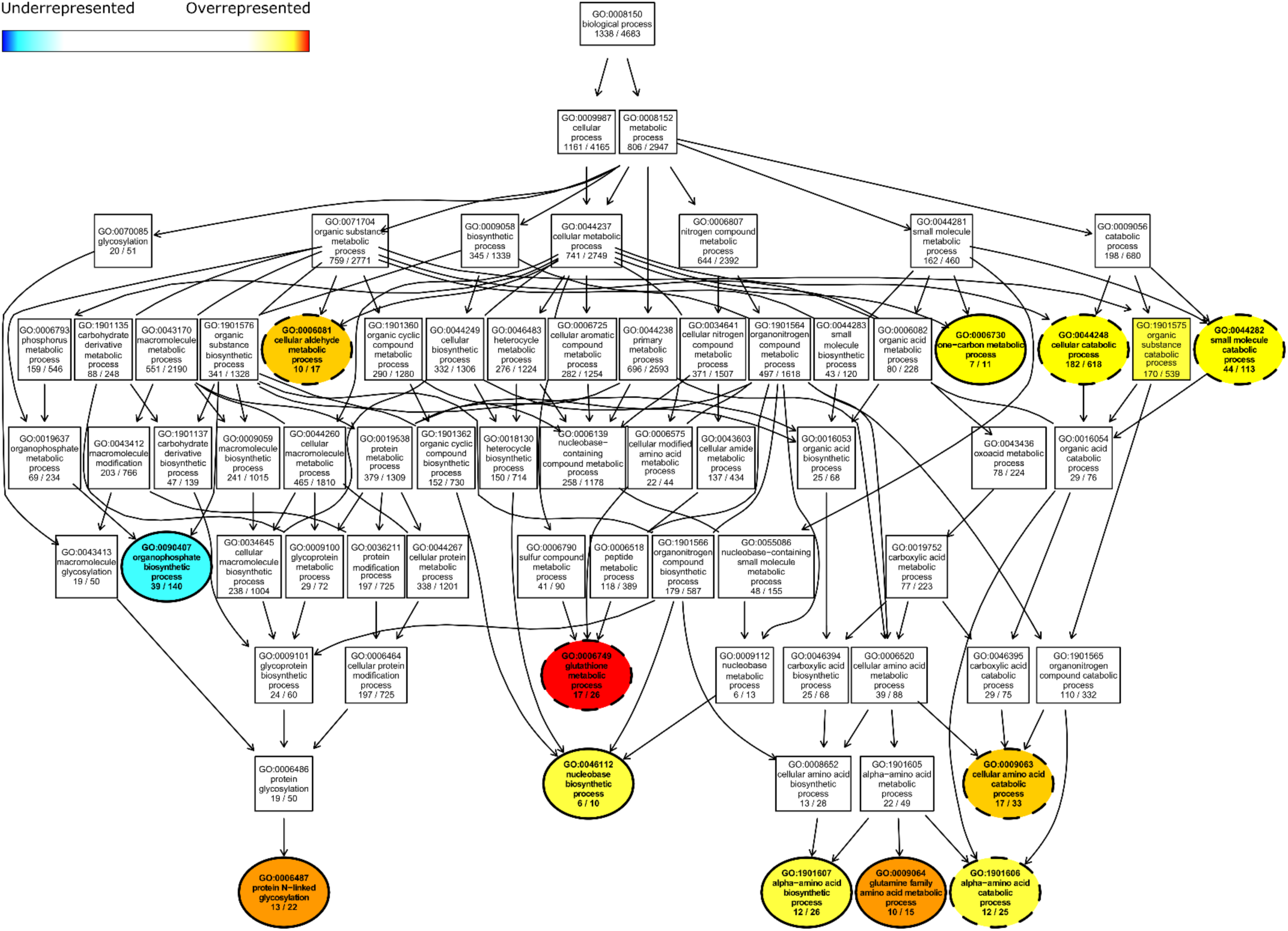
Amino acid metabolism processes show differential enrichment between males of *D. prolongata* and *D. carrolli*. Directed Acyclic Graph (DAG) of significant GO terms and their parent terms in biological processes. Significant (p < 0.05) and non-significant GO terms are color-coded and represented by ellipses and rectangular boxes, respectively. Significant GO terms can be underrepresented (blue) or overrepresented (red) based on Fisher’s exact test. Arrows indicate hierarchical relationships. GO terms at the same level are positioned at the same vertical position. Significant GO terms that are also enriched between females of *D. prolongata* and *D. carrolli* have dashed borders.

**Figure S4D.**
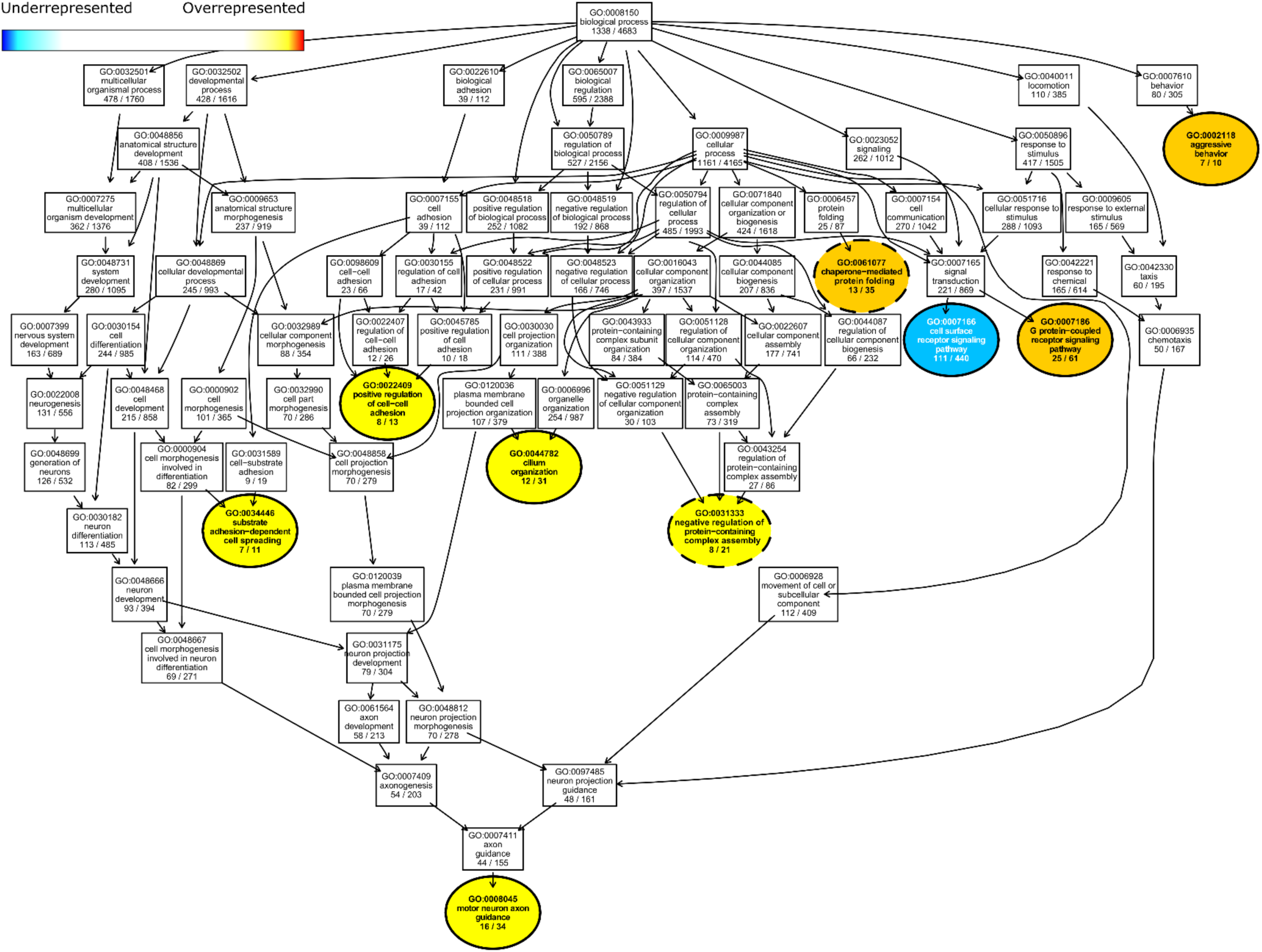
Signal transduction, cell-cell adhesion, and aggressive behavior show differential enrichment between males of *D. prolongata* and *D. carrolli*. Directed Acyclic Graph (DAG) of significant GO terms and their parent terms in biological processes. Significant (p < 0.05) and non-significant GO terms are color-coded and represented by ellipses and rectangular boxes, respectively. Significant GO terms can be underrepresented (blue) or overrepresented (red) based on Fisher’s exact test. Arrows indicate hierarchical relationships. GO terms at the same level are positioned at the same vertical position. Significant GO terms that are also enriched between females of *D. prolongata* and *D. carrolli* have dashed borders.

**Figure S5.**
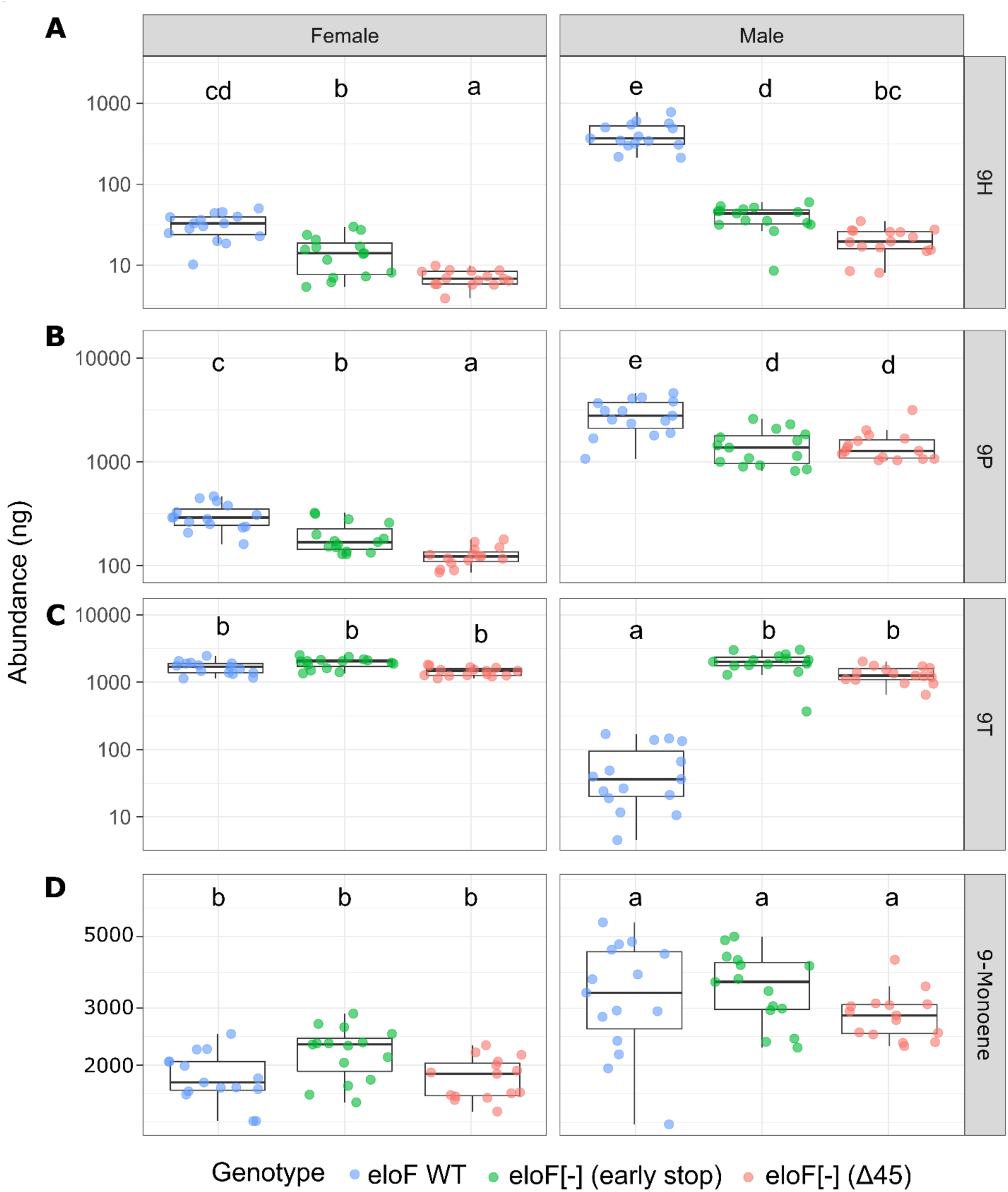
*eloF* is responsible for elongating the precursors of long-chain 9-monoenes. Boxplots showing the abundance of 9H (A), 9P (B), 9T (C), and the aggregate 9-Monoenes (D) across genotypes in each sex, with abundance in nanograms shown on log_10_ scale. Overlayed jitter points are samples of each sex * genotype combination, with color-coded genotypes. Significance of all pairwise comparisons (Tukey HSD test followed by significant omnibus ANOVA F-tests) are summarized in the format of compact letter display (using R packages “multicomp” and “lsmeans”). Note the decrease in the abundance of 9P and 9H in both sexes, and an increase in the abundance of 9T in males, in *eloF* mutants, while the total abundance of 9-monoenes remains approximately constant.

**Figure S6.**
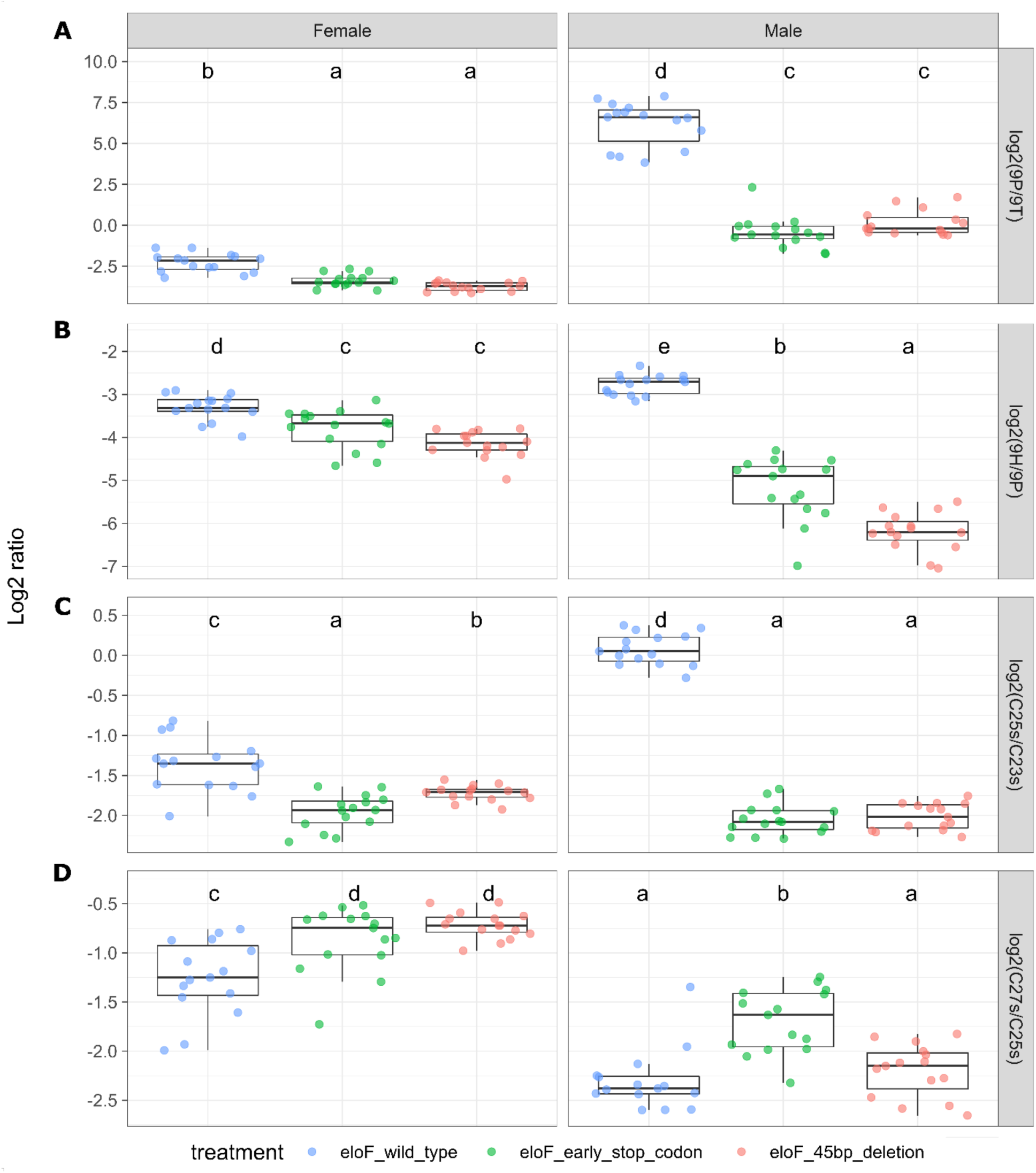
*eloF* is responsible for elongating the precursors of long-chain CHCs. Boxplots showing the log_2_ ratio of 9-monoenes (A-B) and other CHCs (C-D) with adjacent odd-numbered carbons across genotypes in each sex.

**Figure S7.**
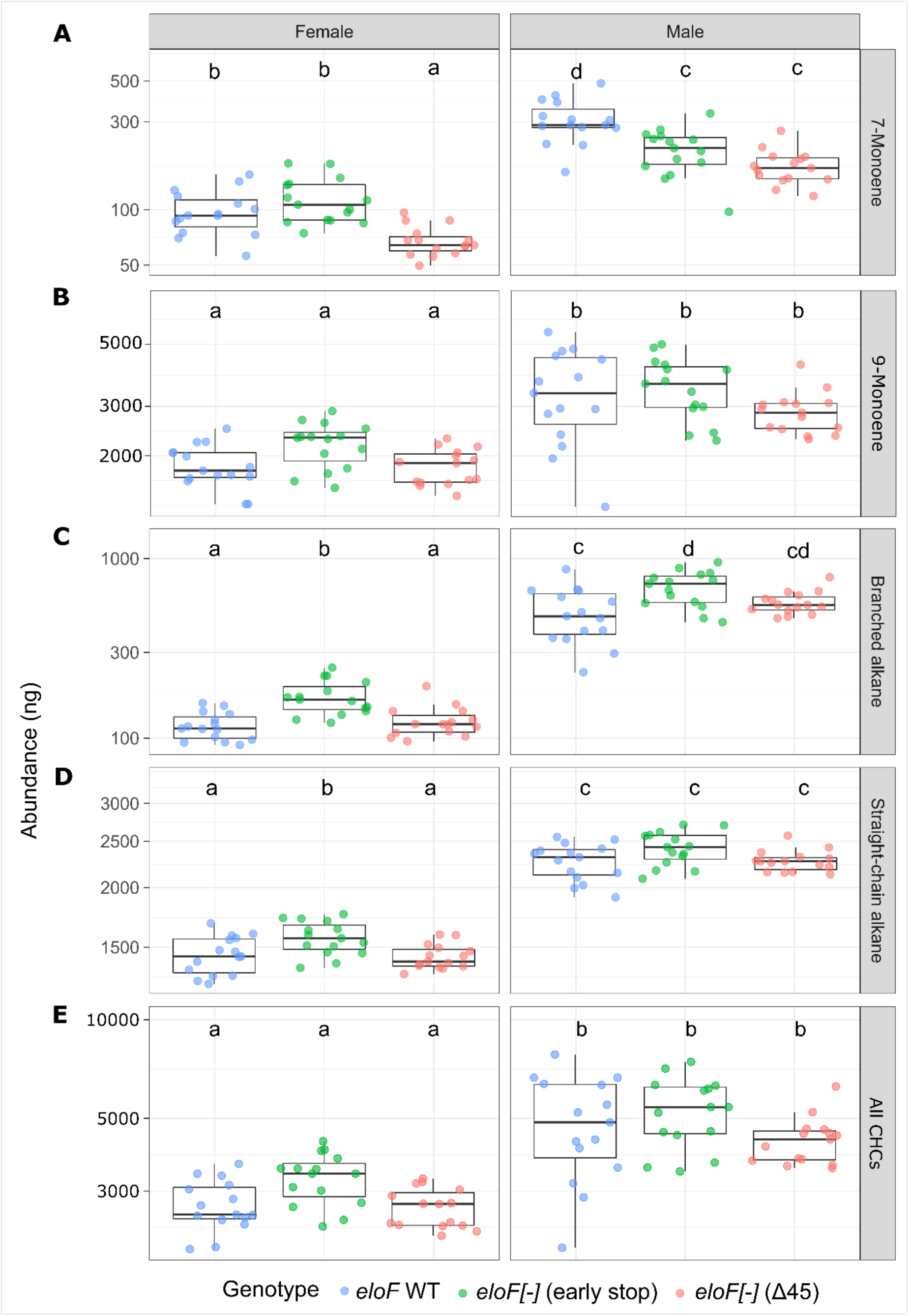
Little effect of *eloF* mutations on total CHC abundance. Boxplots showing the aggregate abundance of 7-Monoenes (A), 9-Monoenes (B), branched alkanes (C), straight-chain alkanes (D), and overall CHCs (E) across genotypes in each sex, with abundance in nanograms shown on log_10_ scale.

**Figure S8.**
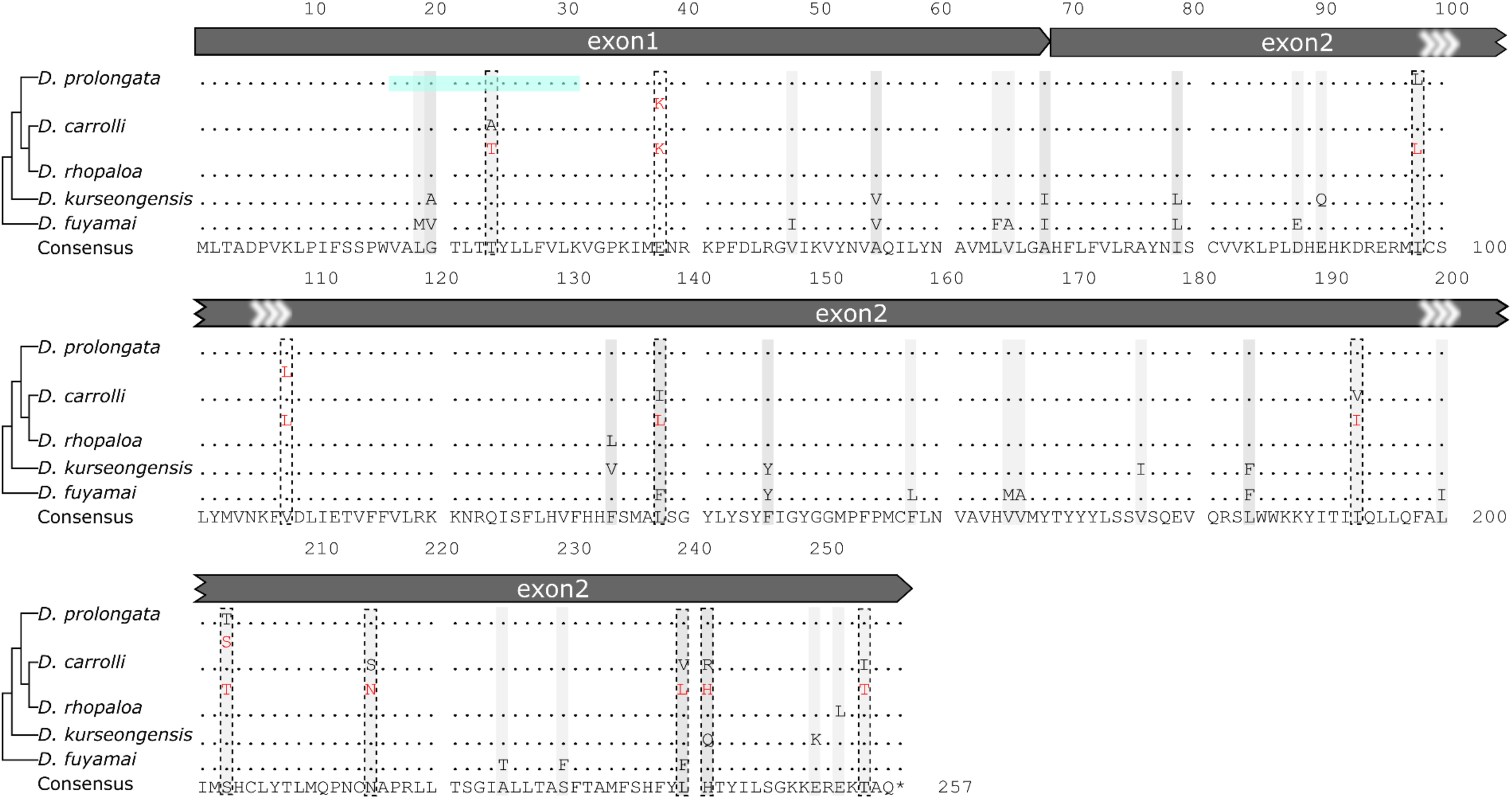
No fixed protein sequence differences between *D. prolongata* and *D. carrolli eloF* orthologs. Multiple alignment on translated amino acid sequences across five species in the *rhopaloa* species subgroup, with species phylogeny on the left and the consensus sequence at the bottom. Numbers above the consensus sequence are coordinates showing the consensus length (257 AA). For the alleles of each species, site-wise disagreement from the consensus is represented in gray shade. For *D. carrolli* and *D. prolongata*, single nucleotide polymorphisms (SNPs) that lead to changes in amino acids are highlighted in red. Polymorphic sites are represented in dashed rectangles. In *D. prolongata*, amino acid sequences deleted in one CRISPR mutant (*eloF*[-] Δ45) are in cyan shade. Feature annotations are displayed above the protein sequence, with dark gray boxes representing *eloF* exons. All features have their direction labeled as arrowheads.

**Figure S9.**
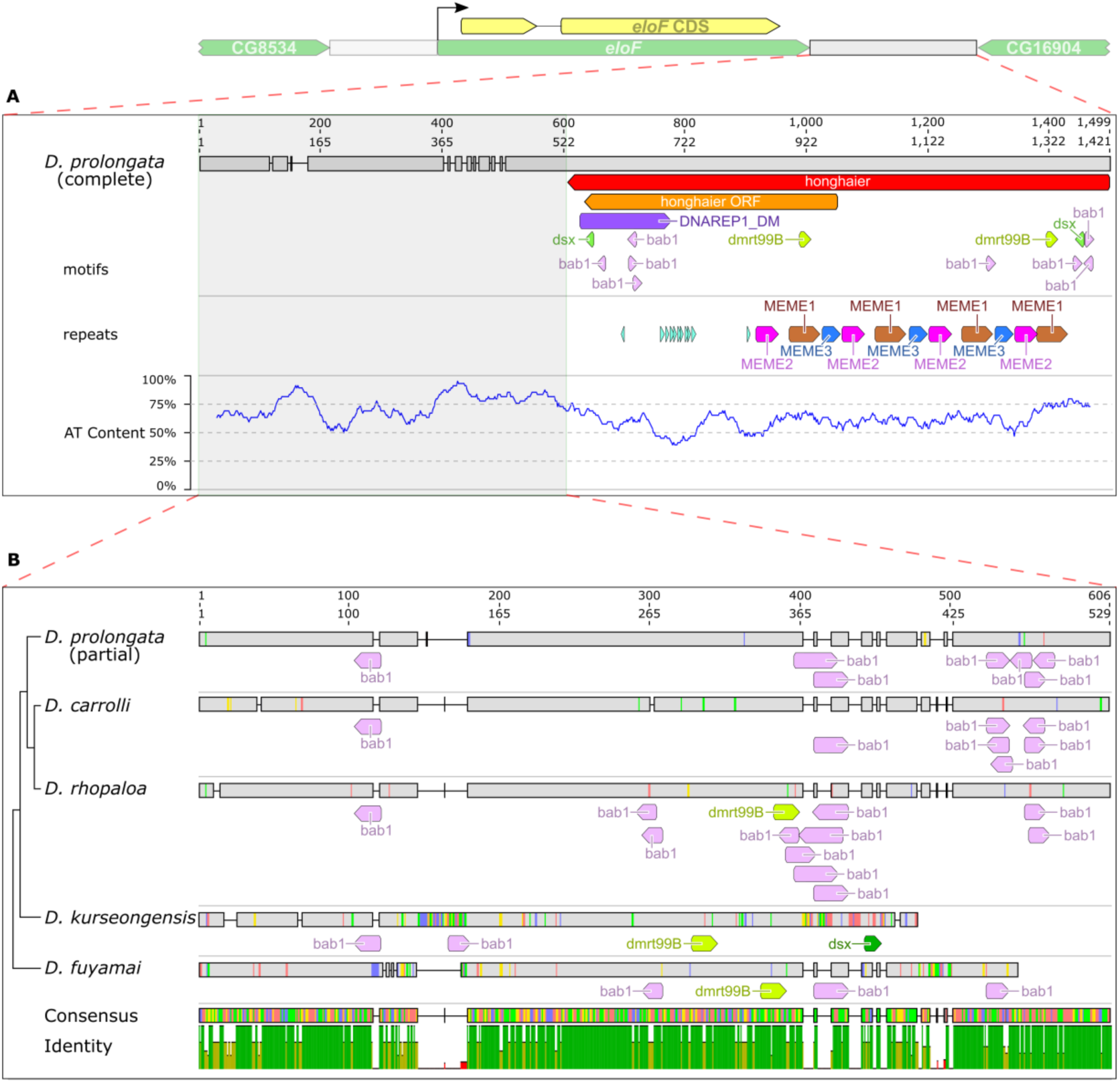
*D. prolongata*-specific *honghaier* insertion in the downstream region of *eloF*. (A) The downstream region of *eloF* in *D. prolongata*, showing the insertion of the TE-like repetitive element *honghaier*. Feature annotations are displayed below DNA sequence, with the red box representing the *honghaier* insertion, the orange box representing its predicted ORF, and the purple box showing the BLAST hit to the DNAREP_DM1 transposable element (Dfam). The motif track shows putative binding sites for transcription factors including *dsx* (JASPAR, dark green), *dsx* (FlyReg, light green), *dmrt99B* (JASPAR, yellow-green), and *bab1* (iDMMPMM, pink). The repeat track includes short TGTC repeats (cyan) and three *de novo* motifs: MEME-1 (brown), MEME-2 (pink), and MEME-3 (steel blue). (B) Alignment of the conserved downstream region of *eloF* (shaded region in A) across species, with species phylogeny on the left and consensus sequence at the bottom. Numbers above the DNA sequence are coordinates showing the length of the consensus (606 bp) and alignment (529 bp). For the alleles of each species, nucleotide-wise disagreement from the consensus is represented in a color-coded vertical line for nucleotide substitutions (A: red, C: blue, G: yellow, T: green), and a horizontal line for nucleotide deletions. The track of percent identity is color coded as follows: green for perfect (100%) agreement, yellow-green for intermediate (30-99%) agreement, and red for low (<30%) agreement. All features have their direction labeled as arrowheads when applicable.

**Figure S10.**
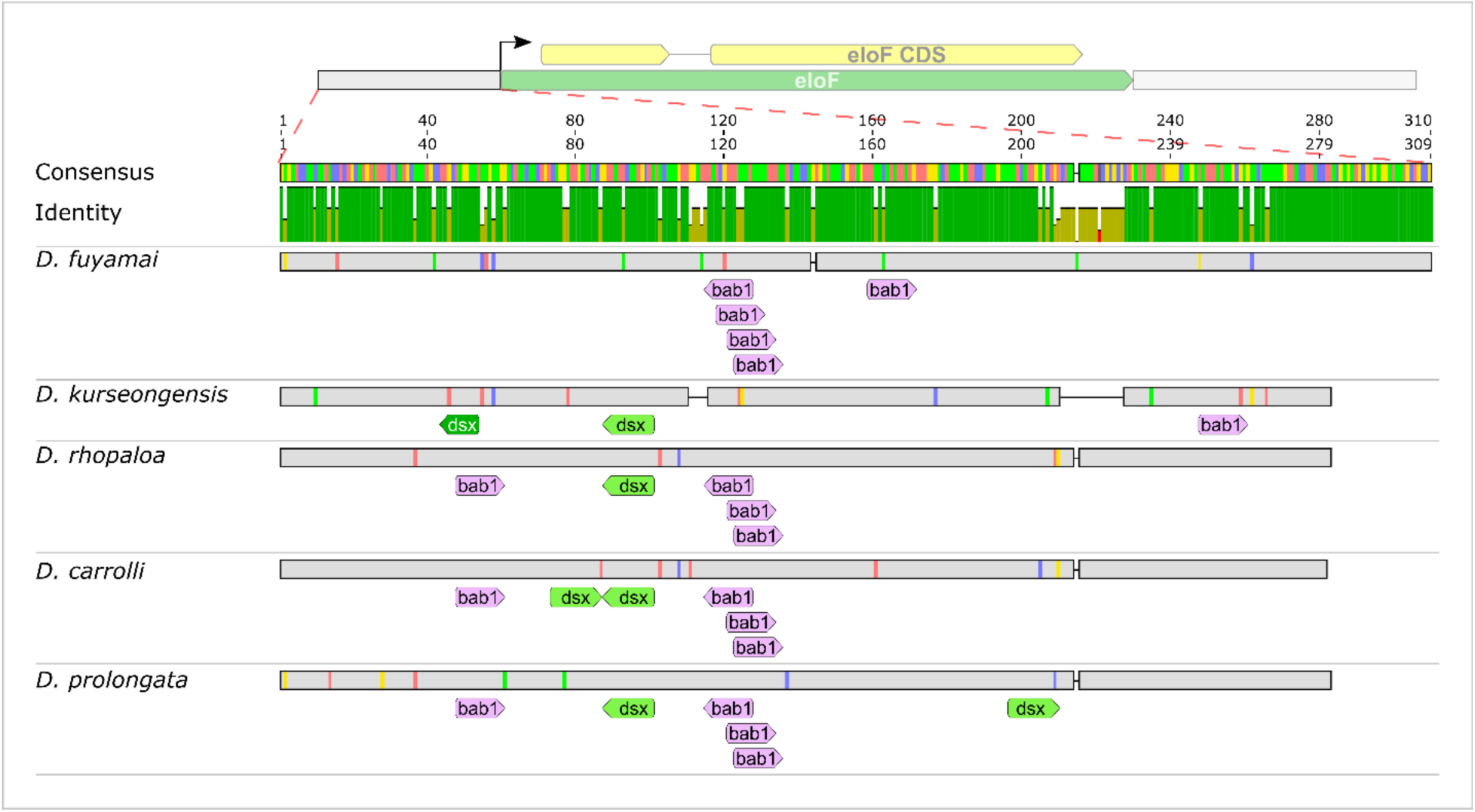
The upstream region of *eloF* is conserved in the *rhopaloa* species subgroup. Multiple alignment of the upstream region of *eloF*, with schematic gene structure displayed on top. The track of percent identity is color-coded as follows: green for perfect (100%) agreement, yellow-green for intermediate (30-99%) agreement, and red for low (<30%) agreement. Numbers above the percent identity track are coordinates showing the length of the consensus (310 bp) and alignment (309 bp). For alleles from each species, nucleotide-wise disagreement from the consensus is represented in a color-coded vertical line for nucleotide substitutions (A: red, C: blue, G: yellow, T: green), and a horizontal line for nucleotide deletions. Predicted transcription factor (TF) binding motifs are displayed below the DNA sequence as follows: *dsx* (JASPAR, dark green); *dsx* (FlyReg, light green); *bab1* (iDMMPMM, pink). All features have their direction labeled as arrowheads when applicable.

**Figure S11.**
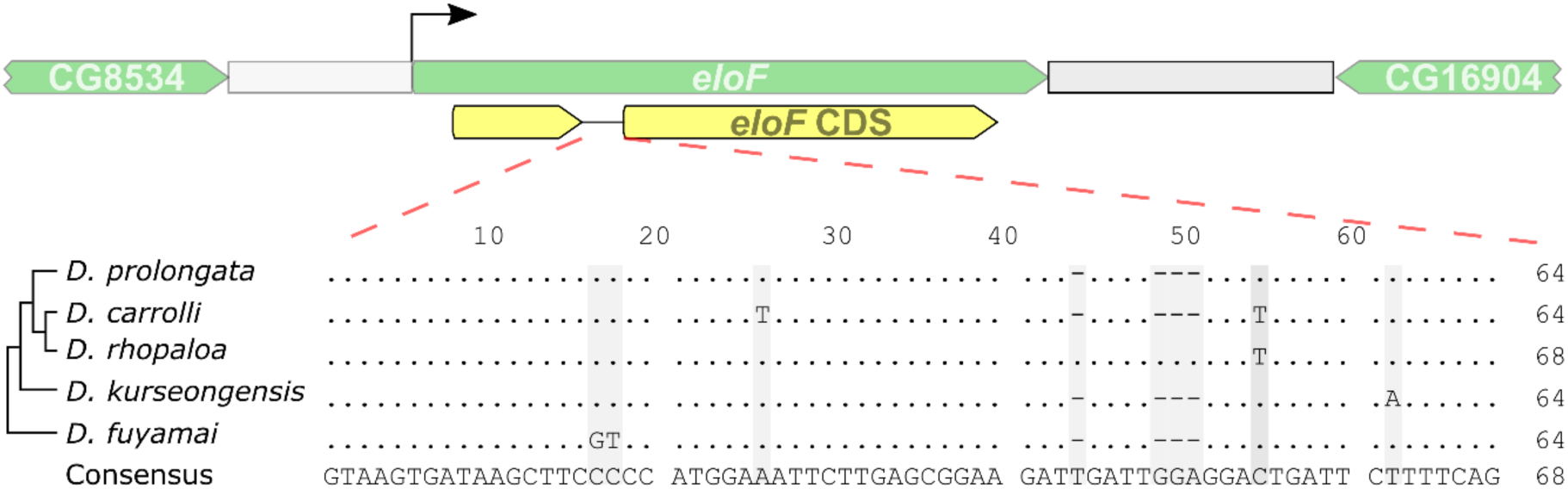
The intron of *eloF* in conserved in the *rhopaloa* species subgroup. Multiple alignment of the intronic region of *eloF*, with genomic context displayed on top. Numbers above the DNA sequence are coordinates showing the consensus length (68 bp). For the alleles of each species, site-wise disagreement from the consensus is represented in gray shade. No sex (*dsx*) or tissue (*bab1*) motifs were identified.

**Figure S12.**
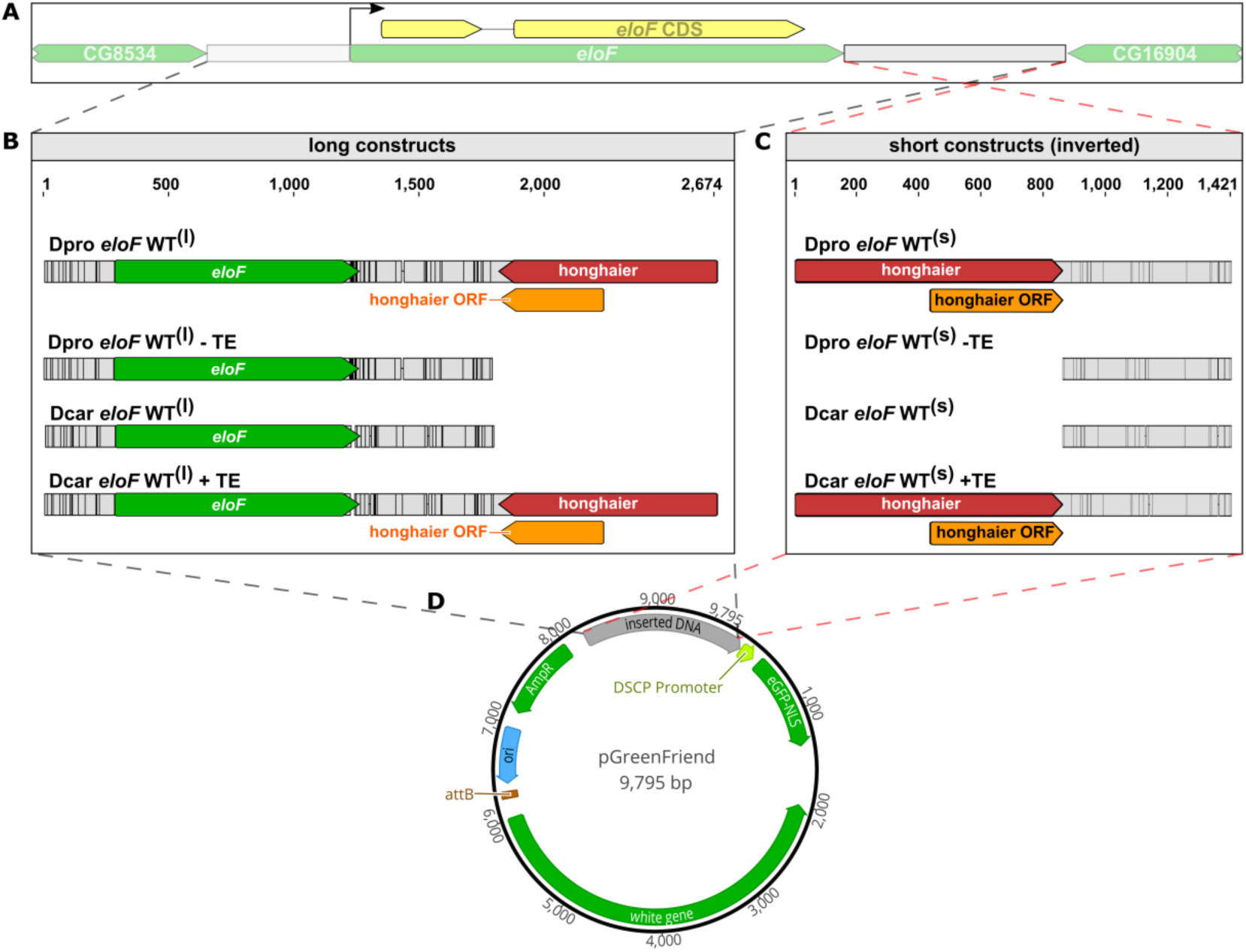
Design of GFP reporter constructs containing *eloF* sequences. (A) Schematic illustration of the *eloF* locus and the two flanking genes. (B) “Long” constructs containing the entire *eloF* locus including flanking sequences. *Dpro eloF* WT^(l)^ and *Dcar eloF* WT^(l)^ carry wild-type *eloF* loci from *D. prolongata* and *D. carrolli*, respectively. The other two constructs were made by removing the *honghaier* TE insertion from the *D. prolongata* sequence (*Dpro eloF* WT^(l)^-TE) or adding the *D. prolongata honghaier* insertion to the *D. carrolli* sequence (*Dcar eloF* WT^(l)^+TE). The *eloF* locus is placed into the pGreenFriend vector in the forward orientation, so that *eloF* is transcribed in the same direction as GFP while the *honghaier* insertion is in the opposite direction. (C). “Short” constructs containing only the downstream *eloF* sequences. As in the “long” constructs, two constructs contain the wild-type alleles from *D. prolongata* and *D. carrolli*, while the other two were made by TE swap. Here, the downstream *eloF* sequences are placed into the pGreenFriend vector in the flipped orientation, so that the direction of the *honghaier* insertion is the same as GFP transcription. In (B) and (C), alignment coordinates are displayed on top. Black lines indicate disagreement between the *D. prolongata* and *D. carrolli* alleles, vertical for single nucleotide variants and horizontal for short indels. Feature annotations are displayed below DNA sequence, with green box representing genes, yellow box representing CDS, red box representing the *honghaier* insertion, and the orange box representing its predicted ORF. All features have their direction labeled by arrowheads when applicable. (D) Schematic illustration of pGreenFriend vector, where GFP is driven by the *Drosophila* synthetic core promoter (DSCP, yellow-green).

**Figure S13.**
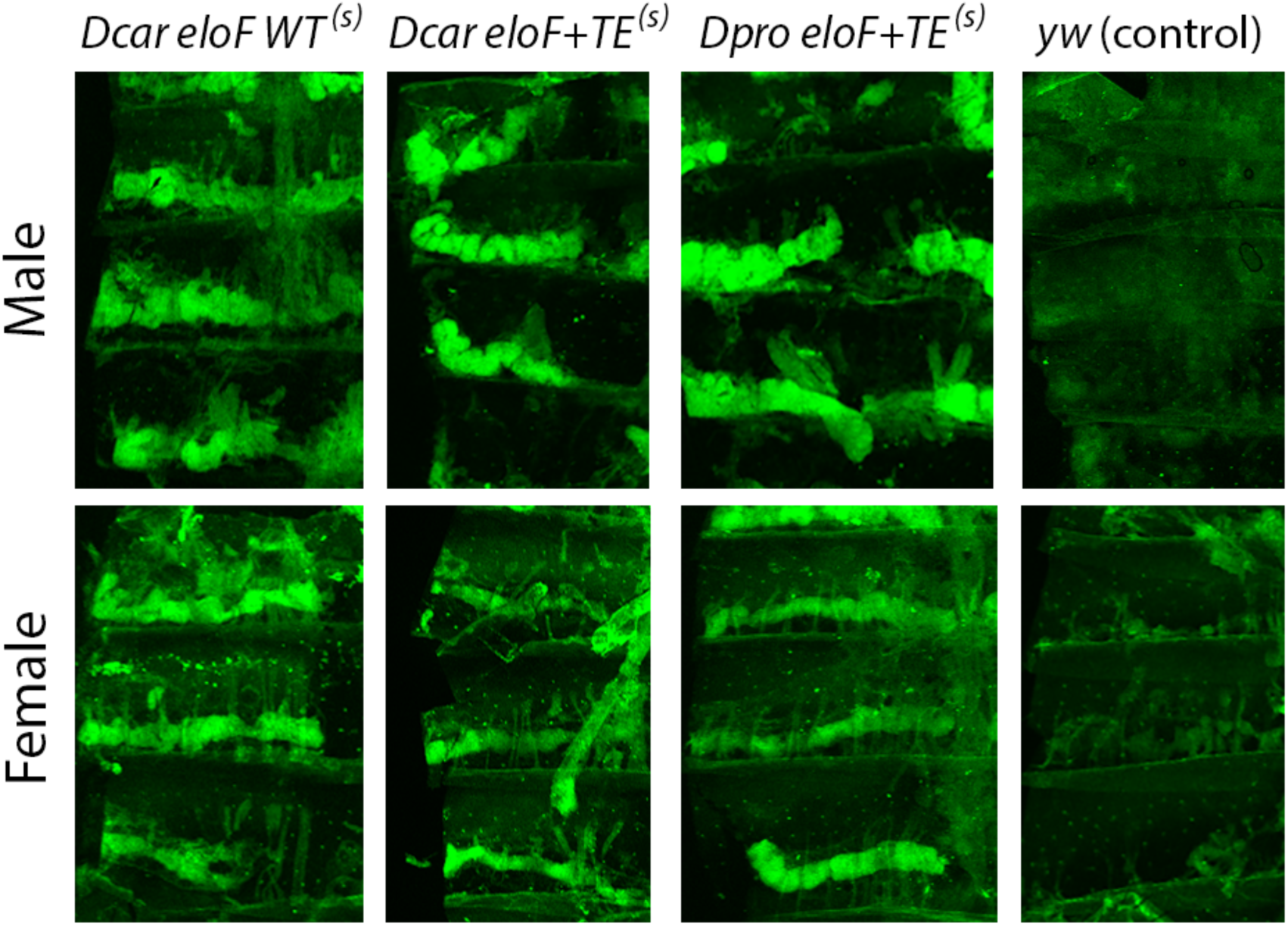
*eloF* downstream sequences drive GFP expression in adult abdominal oenocytes. Confocal images of GFP protein stained with anti-GFP antibodies, showing dissected male and female dorsal abdominal body walls. Non-transgenic *yw* flies are used as a negative control. Transgenic flies carry the “short” constructs containing the *eloF* downstream region (see Fig S12).

**Figure S14.**
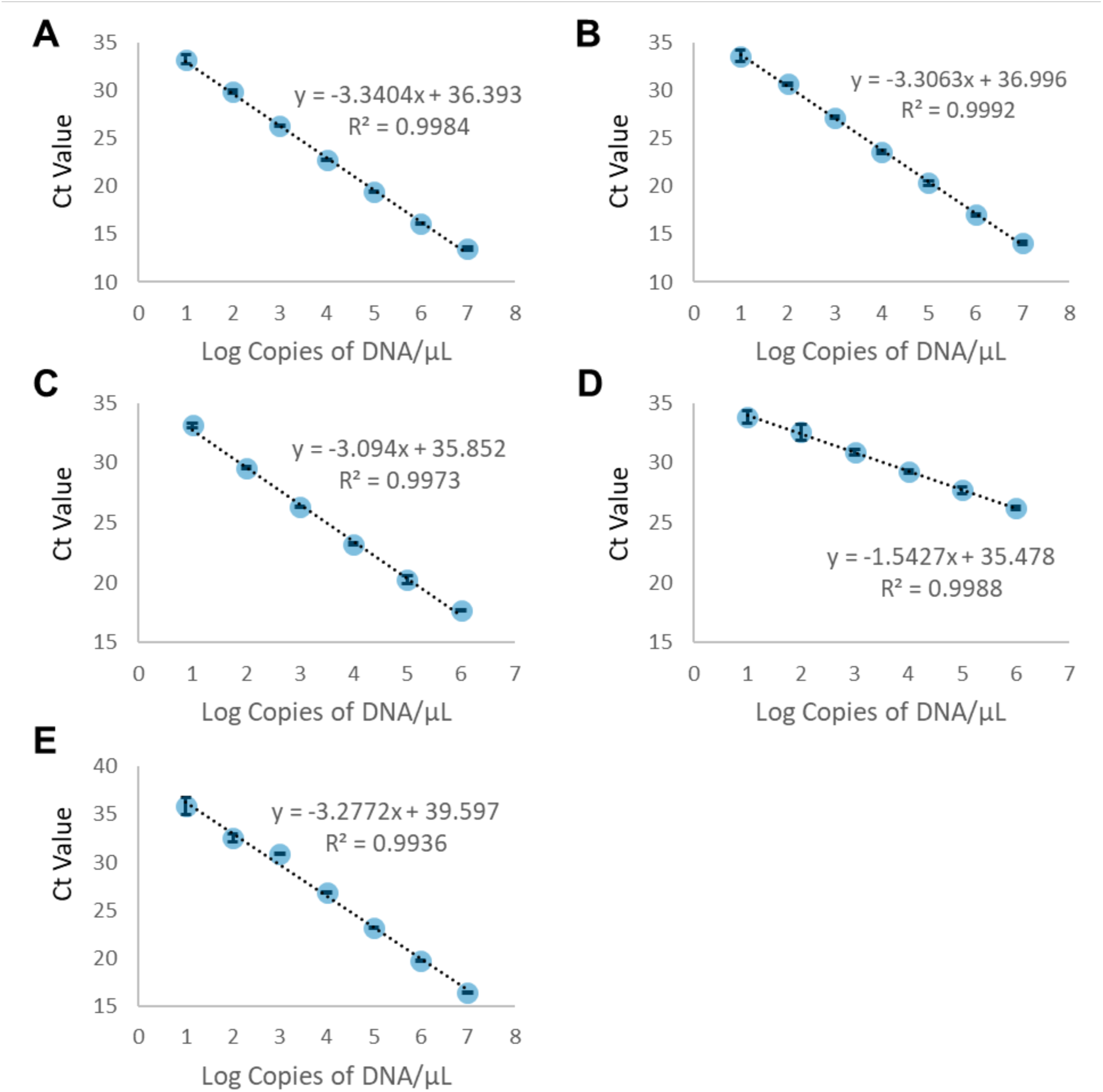
All primers used for quantitative PCR have near-perfect amplification performance. Standard curves of *Rpl32* are based on cDNA from mixed-sex whole-body RNA of *D. prolongata* (A) and *D. carrolli* (B). Standard curves of *eloF* are based on cDNA from mixed-sex whole-body RNA of *D. prolongata* (C) and *D. carrolli* (D). Standard curve of GFP is based on empty pGreenFriend vector (E). Dilution factors are 10-fold for (A), (B) and (E); 8-fold for (C), and 3-fold for (D). Points represent average values, with error bars showing standard deviations calculated from three technical replicates. Lines represent the best linear fit, showing the estimated equation and coefficient of determination (R^2^).

**Figure S15.**
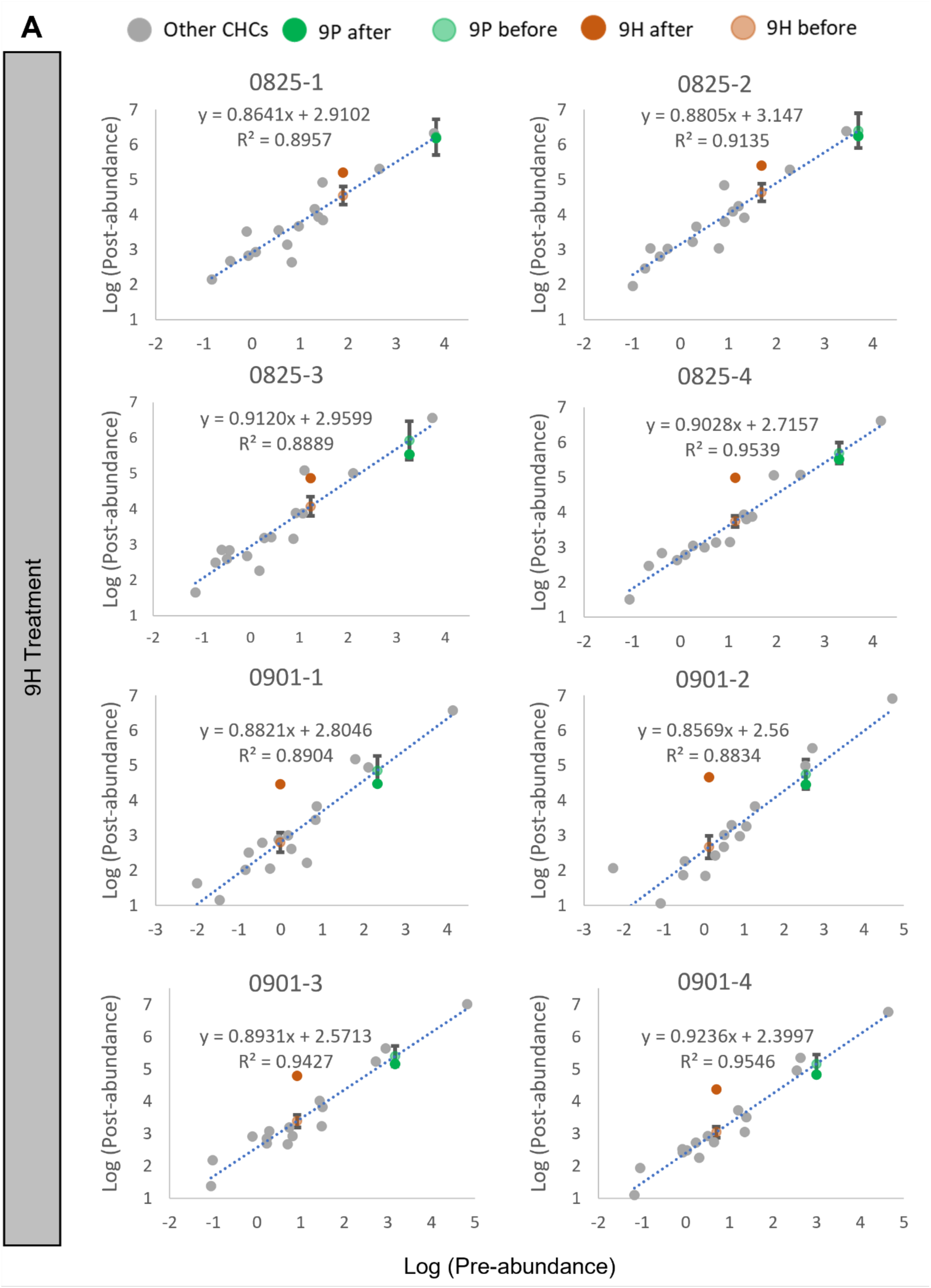
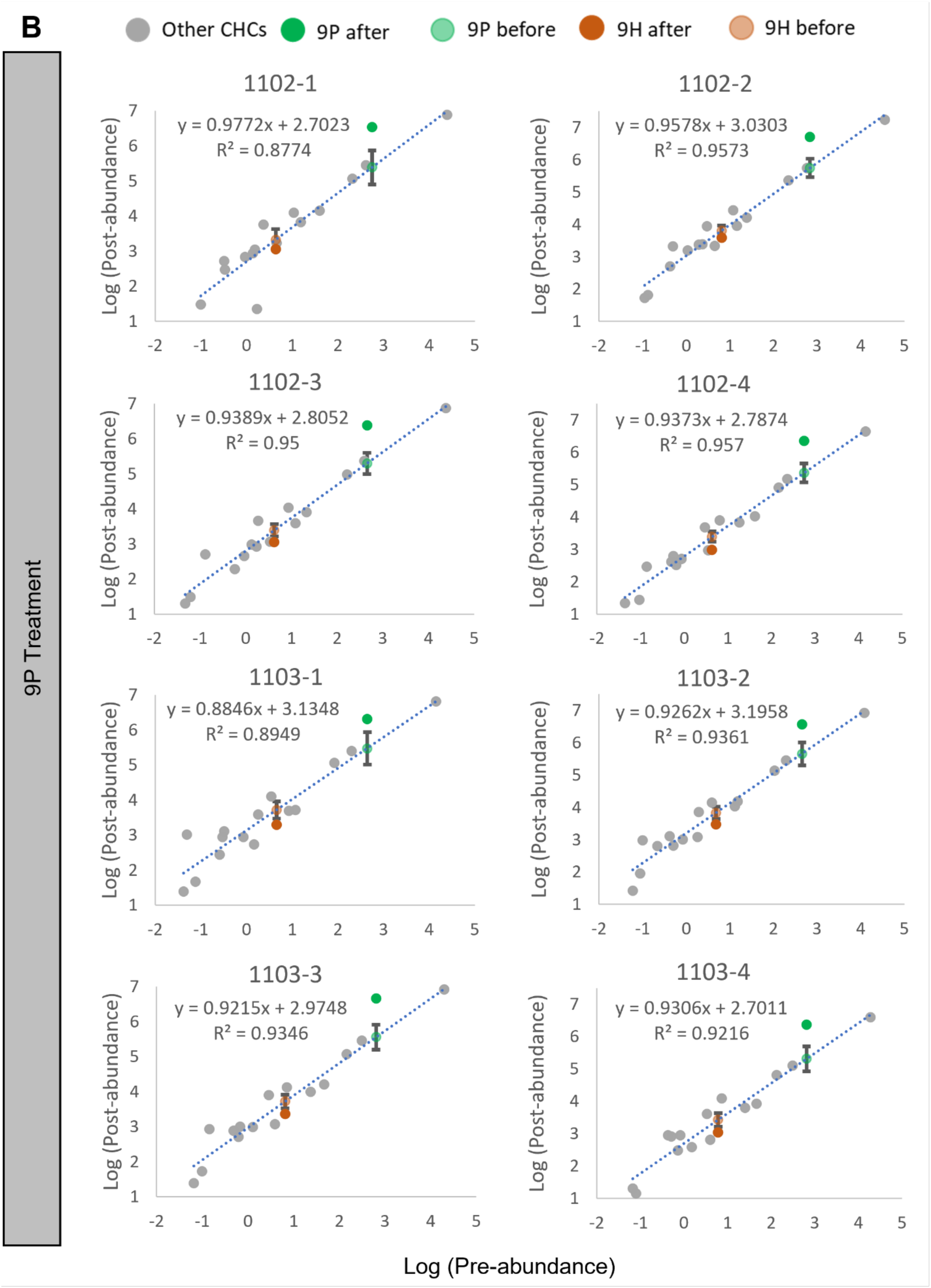
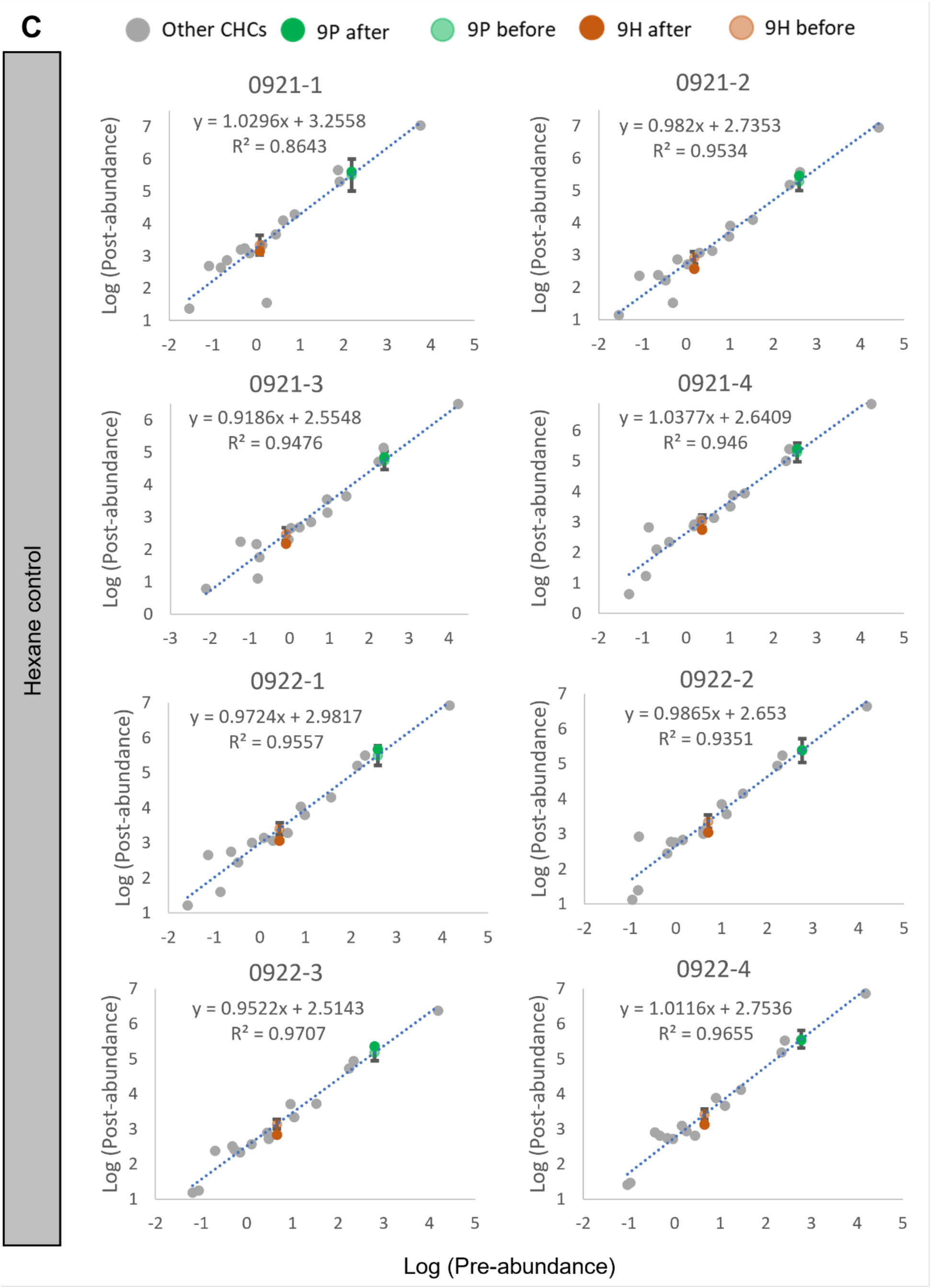
Calibration of candidate CHC transfer based on unperfumed CHCs. Panels of calibration standard curves for 9H treatment (A), 9P treatment (B), and hexane control (C). Each panel represents CHC profiles sampled from an independent group of 8 flies subjected to the same experimental procedures as those used in behavioral studies. Within each panel, each point represents individual CHC, with its abundance before perfuming procedure (pre-abundance) indicated as the x coordinate value, and abundance after perfuming procedure (post-abundance) indicated as the y coordinate value. CHCs other than the spiked-in compound (9P or 9H) were used to build the standard curve (dotted blue line), showing the estimated equation and coefficient of determination (R^2^). Counterfactual post-perfuming abundances of 9P and 9H are estimated from standard curves (light green point for 9P and orange point for 9H) as if no synthetic compounds were added, along with their 95% confidence interval (error bars). Abundance is measured in nanograms and standardized to abundance per individual fly.

**Table S1.** Significant GO terms in the comparison between males of *D. prolongata* and *D. carrolli*.

| GO.ID | Term | Annotated | Significant | Expected | Fisher | KS | Rank in<br>KS | Rank in<br>Fisher | Mean<br>rank |
| --- | --- | --- | --- | --- | --- | --- | --- | --- | --- |
| GO:0006749 | glutathione metabolic process | 26 | 17 | 7.43 | 1.00E-04 | 3.00E-04 | 2 | 1 | 1.5 |
| GO:0030148 | sphingolipid biosynthetic process | 29 | 17 | 8.29 | 0.00067 | 0.0047 | 9 | 2 | 5.5 |
| GO:0032504 | multicellular organism<br>reproduction | 649 | 163 | 185.43 | 0.00171 | 0.0014 | 4 | 3 | 3.5 |
| GO:0009064 | glutamine family amino acid<br>metabolic process | 15 | 10 | 4.29 | 0.00241 | 0.0036 | 8 | 4 | 6 |
| GO:0006487 | protein N-linked glycosylation | 22 | 13 | 6.29 | 0.00263 | 0.0076 | 14 | 5 | 9.5 |
| GO:0009063 | cellular amino acid catabolic<br>process | 33 | 17 | 9.43 | 0.00442 | 5.00E-04 | 3 | 6 | 4.5 |
| GO:0035167 | larval lymph gland hemopoiesis | 41 | 15 | 11.71 | 0.00459 | 0.0143 | 28 | 7 | 17.5 |
| GO:0042761 | very long-chain fatty acid<br>biosynthetic process | 14 | 9 | 4 | 0.00575 | 0.0138 | 27 | 8 | 17.5 |
| GO:0061077 | chaperone-mediated protein<br>folding | 35 | 13 | 10 | 0.00581 | 0.0079 | 16 | 9 | 12.5 |
| GO:0007442 | hindgut morphogenesis | 28 | 9 | 8 | 0.00668 | 0.0147 | 29 | 10 | 19.5 |
| GO:0002118 | aggressive behavior | 10 | 7 | 2.86 | 0.00785 | 0.0062 | 12 | 11 | 11.5 |
| GO:0006720 | isoprenoid metabolic process | 18 | 9 | 5.14 | 0.00865 | 0.0097 | 19 | 12 | 15.5 |
| GO:0006081 | cellular aldehyde metabolic<br>process | 17 | 10 | 4.86 | 0.00868 | 0.0028 | 7 | 13 | 10 |
| GO:0007186 | G protein-coupled receptor<br>signaling pathway | 61 | 25 | 17.43 | 0.00881 | 0.0212 | 38 | 14 | 26 |
| GO:0022409 | positive regulation of cell-cell<br>adhesion | 13 | 8 | 3.71 | 0.0133 | 0.0274 | 44 | 16 | 30 |
| GO:0007552 | metamorphosis | 331 | 89 | 94.57 | 0.01397 | 0.0121 | 24 | 17 | 20.5 |
| GO:0044248 | cellular catabolic process | 618 | 182 | 176.57 | 0.01423 | 0.0238 | 40 | 18 | 29 |
| GO:0006641 | triglyceride metabolic process | 29 | 13 | 8.29 | 0.01461 | 0.0265 | 42 | 19 | 30.5 |
| GO:0007166 | cell surface receptor signaling<br>pathway | 440 | 111 | 125.71 | 0.01534 | 0.0102 | 20 | 20 | 20 |
| GO:0006730 | one-carbon metabolic process | 11 | 7 | 3.14 | 0.0163 | 0.0019 | 5 | 22 | 13.5 |
| GO:0019367 | fatty acid elongation, saturated<br>fatty acid | 11 | 7 | 3.14 | 0.0163 | 0.0291 | 49 | 23 | 36 |
| GO:0034625 | fatty acid elongation,<br>monounsaturated fatty acid | 11 | 7 | 3.14 | 0.0163 | 0.0291 | 50 | 24 | 37 |
| GO:0034626 | fatty acid elongation,<br>polyunsaturated fatty acid | 11 | 7 | 3.14 | 0.0163 | 0.0291 | 51 | 25 | 38 |
| GO:0034446 | substrate adhesion-dependent<br>cell spreading | 11 | 7 | 3.14 | 0.0163 | 0.0333 | 56 | 26 | 41 |
| GO:0035336 | long-chain fatty-acyl-CoA<br>metabolic process | 11 | 7 | 3.14 | 0.0163 | 0.0394 | 65 | 27 | 46 |
| GO:0008045 | motor neuron axon guidance | 34 | 16 | 9.71 | 0.01655 | 0.0127 | 25 | 28 | 26.5 |
| GO:0044282 | small molecule catabolic process | 113 | 44 | 32.29 | 0.01741 | 0.0187 | 36 | 29 | 32.5 |
| GO:0015849 | organic acid transport | 37 | 15 | 10.57 | 0.02317 | 0.0083 | 18 | 31 | 24.5 |
| GO:0044782 | cilium organization | 31 | 12 | 8.86 | 0.02323 | 0.0181 | 35 | 32 | 33.5 |
| GO:0006605 | protein targeting | 88 | 28 | 25.14 | 0.02325 | 0.0418 | 68 | 33 | 50.5 |
| GO:0000578 | embryonic axis specification | 76 | 18 | 21.71 | 0.0236 | 0.0283 | 48 | 34 | 41 |
| GO:0031333 | negative regulation of protein-<br>containing complex assembly | 21 | 8 | 6 | 0.02568 | 0.0282 | 47 | 36 | 41.5 |
| GO:1901606 | alpha-amino acid catabolic<br>process | 25 | 12 | 7.14 | 0.03047 | 0.0021 | 6 | 41 | 23.5 |
| GO:0055085 | transmembrane transport | 283 | 94 | 80.86 | 0.03356 | 0.0341 | 59 | 42 | 50.5 |
| GO:0046112 | nucleobase biosynthetic process | 10 | 6 | 2.86 | 0.03749 | 0.0351 | 60 | 43 | 51.5 |
| GO:0090407 | organophosphate biosynthetic<br>process | 140 | 39 | 40 | 0.03853 | 0.0059 | 11 | 47 | 29 |
| GO:1901607 | alpha-amino acid biosynthetic<br>process | 26 | 12 | 7.43 | 0.04225 | 0.0369 | 64 | 48 | 56 |
| GO:0016042 | lipid catabolic process | 74 | 27 | 21.14 | 0.04811 | 0.0403 | 67 | 50 | 58.5 |
| GO:0007472 | wing disc morphogenesis | 213 | 58 | 60.86 | 0.04826 | 0.0271 | 43 | 51 | 47 |

Fisher: raw p-values from Fisher's exact test
KS: raw p-values from the Kolmogorov-Smirnov test

**Table S2.** qPCR analysis of GFP transcript expression driven by *eloF* “long” constructs (complete *eloF* locus including flanking regions).

| Sex | Genotype |  | Reference gene (*Rpl32*) | Gene of interest (*GFP*) |
| --- | --- | --- | --- | --- |
| Female | Dpro *eloF* WT<sup>(l)</sup> | Replicate 1 | 14.81070235 | >40 |
|  |  | Replicate 2 | 14.46329199 | >40 |
|  |  | Replicate 3 | 14.51088284 | >40 |
|  |  | NRT control |  | >40 |
|  | Dcar *eloF* WT<sup>(l)</sup> | Replicate 1 | 14.79372247 | >40 |
|  |  | Replicate 2 | 14.66467155 | >40 |
|  |  | Replicate 3 | 14.37498109 | >40 |
|  |  | NRT control |  | >40 |
| Male | Dpro *eloF* WT<sup>(l)</sup> | Replicate 1 | 13.84065353 | 37.68637158 |
|  |  | Replicate 2 | 14.91291474 | >40 |
|  |  | Replicate 3 | 13.9237942 | 39.51370016 |
|  |  | NRT control |  | >40 |
|  | Dcar *eloF* WT<sup>(l)</sup> | Replicate 1 | 14.06775107 | 35.15658614 |
|  |  | Replicate 2 | 13.83850168 | 34.86276857 |
|  |  | Replicate 3 | 13.65032641 | 34.52029242 |
|  |  | NRT control |  | >40 |

Note: qPCR amplification Ct values are reported, where non-detects are labeled as >40.

**Table S3.** Cuticular lipid description.

| Compound name | Abbrevia<br>tion | Chain<br>length | Chemical class | isConsen<br>sus | Kovat's<br>index | Characteristic<br>ions (m/z) |
| --- | --- | --- | --- | --- | --- | --- |
| 9-Heneicosene | 9Hen | 21 | 9-Monoene | FALSE | NA | 294 |
| 7-Heneicosene | 7Hen | 21 | 7-Monoene | FALSE | NA | 294 |
| n-Heneicosane | nC21 | 21 | Straight-chain<br>alkane | TRUE | 2100 | 296 |
| 9-Docosene | 9D | 22 | 9-Monoene | FALSE | 2173~21<br>80 | 308 |
| 11-cis-vaccenyl<br>acetate | cVA | 20 | Acetate ester | FALSE | 2190~21<br>91 | 250, 310 |
| n-Docosane | nC22 | 22 | Straight-chain<br>alkane | TRUE | 2200 | 310 |
| 2-Methyl-<br>docosane | 23Br | 23 | Branched<br>alkane | TRUE | 2263~22<br>64 | 281, 309, 324 |
| x,y-Tricosadiene | xyTD | 23 | Diene | FALSE | 2270~22<br>71 | 320 |
| 9-Tricosene | 9T | 23 | 9-Monoene | TRUE | 2274~22<br>78 | 322 |
| 7-Tricosene | 7T | 23 | 7-Monoene | TRUE | 2280~22<br>82 | 322 |
| n-Tricosane | nC23 | 23 | Straight-chain<br>alkane | TRUE | 2300 | 324 |
| 2-Methyl-<br>tricosane | 24Br | 24 | Branched<br>alkane | FALSE | 2363~23<br>64 | 295, 323, 338 |
| 9-Tetracosene | 9Te | 24 | 9-Monoene | TRUE | 2374~23<br>76 | 336 |
| 7-Tetracosene | 7Te | 24 | 7-Monoene | TRUE | 2377~23<br>79 | 336 |
| n-Tetracosane | nC24 | 24 | Straight-chain<br>alkane | TRUE | 2400 | 338 |
| 2-Methyl-<br>tetracosane | 25Br | 25 | Branched<br>alkane | TRUE | 2463~24<br>64 | 309, 337, 352 |
| 9-Pentacosene | 9P | 25 | 9-Monoene | TRUE | 2475~24<br>78 | 350 |
| 7-Pentacosene | 7P | 25 | 7-Monoene | TRUE | 2482~24<br>84 | 350 |
| n-Pentacosane | nC25 | 25 | Straight-chain<br>alkane | TRUE | 2500 | 352 |
| 9-Hexacosene | 9He | 26 | 9-Monoene | FALSE | 2574~25<br>82 | 364 |
| 2-Methyl-<br>hexacosane | 27Br | 27 | Branched<br>alkane | TRUE | 2663~26<br>64 | 337, 365, 380 |
| 9-Heptacosene | 9H | 27 | 9-Monoene | TRUE | 2676~26<br>77 | 378 |
| 7-Heptacosene | 7H | 27 | 7-Monoene | TRUE | 2683~26<br>85 | 378 |
| n-Heptacosane | nC27 | 27 | Straight-chain<br>alkane | TRUE | 2700 | 380 |
| 2-Methyl-<br>octacosane | 29Br | 29 | Branched<br>alkane | TRUE | 2859~28<br>61 | 365, 393, 408 |
| n-Nonacosane | nC29 | 29 | Straight-chain<br>alkane | FALSE | 2900 | 408 |

**Table S4.** *eloF[-]* mutant behavior.

| Genotype | Mating behavior |  |  |  |  | Fighting behavior |  |  |  |  | Misdirected courtship |  |
| --- | --- | --- | --- | --- | --- | --- | --- | --- | --- | --- | --- | --- |
|  | N | courtship (rate) | mean courtship duration [min] (SE) | leg vibration (rate) | copulation (rate) | mean copulation duration [min] (SE) | N | threatening (rate) | boxing (rate) | mean boxing duration [min] (SE) | N | occurrence (rate) |
| eloF WT | 32 | 21 (0.656) | 3.344 (0.739) | 8 (0.25) | 6 (0.188) | 6.033 (0.506) | 23 | 22 (0.957) | 9 (0.391) | 2.217 (1.016) | 40 | 3 (0.075) |
| eloF<sup>[-]</sup> (Δ45) | 22 | 12 (0.545) | 7.300 (2.334) | 9 (0.409) | 6 (0.273) | 5.467 (0.436) | 15 | 15 (1.000) | **12 (0.800) \*** | 4.913 (1.175) | 40 | 7 (0.175) |
| eloF<sup>[-]</sup> (early stop) | 26 | 17 (0.654) | 4.754 (1.243) | 4 (0.154) | 2 (0.077) | 5.550 (0.350) | 30 | 27 (0.900) | 10 (0.333) | 2.077 (0.800) | 40 | **13 (0.325) \*\*** |

**Table S5.** Sites segregating in the coding region of *eloF* in *D. prolongata* and *D. carrolli*.

| DNA |  |  |  |  | Protein |  |  |  |  |
| --- | --- | --- | --- | --- | --- | --- | --- | --- | --- |
| Position | Dpro_Ref | Dpro_Alt | Dcar_Ref | Dcar_Alt | Position | Dpro_Ref | Dpro_Alt | Dcar_Ref | Dcar_Alt |
| 51 | T |  | C | T | 17 | Val |  | Val | Val |
| 69 | A | G | G | A | 23 | Thr | Thr | Thr | Thr |
| 70 | A |  | G | A | 24 | Thr |  | Ala | Thr |
| 90 | C | G | C | G | 30 | Leu | Leu | Leu | Leu |
| 102 | G | C | C | G | 34 | Pro | Pro | Pro | Pro |
| 112 | G | A | G | A | 38 | Glu | Lys | Glu | Lys |
| 135 | G | A | G | A | 45 | Leu | Leu | Leu | Leu |
| 153 | C | G | C | G | 51 | Val | Val | Val | Val |
| 292 | C |  | A | C | 98 | Leu |  | Ile | Leu |
| 322 | G | T | G | T | 108 | Val | Leu | Val | Leu |
| 411 | T |  | C | T | 137 | Ala |  | Ala | Ala |
| 412 | TTA |  | ATT | TTA | 138 | Leu |  | Ile | Leu |
| 450 | T |  | C | T | 150 | Gly | Gly | Gly | Gly |
| 577 | A |  | G | A | 193 | Ile |  | Val | Ile |
| 583 | T | C | C | T | 195 | Leu | Leu | Leu | Leu |
| 600 | G |  | A | G | 200 | Leu | Leu | Leu | Leu |
| 607 | A |  | T | A | 203 | Thr |  | Ser | Thr |
| 642 | T | C | C | T | 214 | Cys | Cys | Cys | Cys |
| 644 | A |  | G | A | 215 | Asn |  | Ser | Asn |
| 666 | T | C | C | T | 222 | Ser | Ser | Ser | Ser |
| 718 | T |  | G | T | 240 | Leu |  | Val | Leu |
| 722 | A |  | G | A | 241 | His |  | Arg | His |
| 761 | C |  | T | C | 254 | Thr |  | Ile | Thr |
| 765 | C |  | G | C | 255 | Ala |  | Ala | Ala |

Note: Rows shaded in yellow have non-synonymous divergent sites between the two reference genomes, but in at least one species the alternative allele matches the reference allele of the other species.

**Table S6.** High fidelity *honghaier* sequence occurrence.

| Species | honghaier ORF (414bp) count | honghaier (894bp) count |
| --- | --- | --- |
| *D. prolongata* | 1942 | 274 |
| *D. carrolli* | 1617 | 363 |
| *D. rhopaloa* | 3432 | 854 |
| *D. kurseongensis* | 2180 | 310 |
| *D. fuyamai* | 1742 | 11 |
| *D. elegans* | 0 | 0 |
| *D. melanogaster* | 0 | 0 |

Note: using BLASTn 2.2.31+. The cutoffs for *honghaier* high fidelity homologs are >90% query cover and >90% percent identity.

**Table S7.** RNA-seq data summary.

| **Sample** | **Raw Reads** | **Clean Reads** | **Effective Rate (%)** | **Error Rate (%)** | **Q20(%)** | **Q30(%)** | **GC Content (%)** |
| --- | --- | --- | --- | --- | --- | --- | --- |
| Dpro_F_1 | 30344414 | 18193778 | 59.96 | 0.01 | 98.16 | 95.34 | 55.05 |
| Dpro_F_2 | 25460124 | 15361829 | 60.34 | 0.01 | 98.06 | 95.12 | 54.3 |
| Dpro_F_3 | 36316550 | 19245326 | 52.99 | 0.01 | 98.32 | 95.69 | 53.77 |
| Dpro_F_4 | 37396183 | 19525669 | 52.21 | 0.01 | 98.18 | 95.44 | 54.78 |
| Dpro_M_1 | 26796536 | 16587626 | 61.9 | 0.01 | 98.16 | 95.34 | 53.14 |
| Dpro_M_2 | 28896516 | 18187156 | 62.94 | 0.01 | 97.93 | 94.87 | 53.13 |
| Dpro_M_3 | 25315750 | 13188023 | 52.09 | 0.01 | 98.09 | 95.31 | 52.68 |
| Dpro_M_5 | 7206195 | 3294602 | 45.72 | 0.01 | 98.3 | 95.66 | 51.72 |
| Dcar_F_1 | 29174061 | 15503044 | 53.14 | 0.01 | 98.07 | 95.23 | 52.38 |
| Dcar_F_2 | 28112240 | 15288035 | 54.38 | 0.01 | 98.17 | 95.4 | 53.78 |
| Dcar_F_4 | 36082291 | 21015328 | 58.24 | 0.01 | 98.38 | 95.84 | 51.09 |
| Dcar_F_6 | 35006571 | 19051828 | 54.42 | 0.01 | 97.97 | 95.05 | 53.41 |
| Dcar_M_1 | 22562361 | 13618258 | 60.36 | 0.01 | 98.24 | 95.53 | 53.19 |
| Dcar_M_2 | 27694508 | 15030603 | 54.27 | 0.01 | 98.25 | 95.58 | 52.69 |
| Dcar_M_3 | 25947915 | 15636694 | 60.26 | 0.01 | 98.14 | 95.34 | 51.78 |
| Dcar_M_5 | 21910989 | 11529155 | 52.62 | 0.01 | 98.15 | 95.39 | 51.62 |

Raw reads: the total amount of reads of raw data (read1 + read2), every four lines taken as one unit.
Clean reads: the total amount of reads of clean data, each of four lines taken as one unit.
Effective Rate (%): (Clean reads/Raw reads) \* 100%
Error rate: base error rate
Q20, Q30: (Base count of Phred value > 20 or 30) / (Total base count)
GC content: (G & C base count) / (Total base count)

**Table S8.** RNA-seq read mapping statistics.

| Sample | M Aligned<br>(% Aligned) | Overlapping<br>Genes | No Feature | Ambiguous<br>Features | Multi<br>mapping | Unmapped |
| --- | --- | --- | --- | --- | --- | --- |
| Dpro_F_1 | 12.2 (92.8%) | 11112851<br>(84.3%) | 797692<br>(6.1%) | 326756<br>(2.5%) | 796396<br>(6.0%) | 147450<br>(1.1%) |
| Dpro_F_2 | 10.9 (92.9%) | 9764869<br>(83.1%) | 863405<br>(7.3%) | 288170<br>(2.5%) | 709311<br>(6.0%) | 124662<br>(1.1%) |
| Dpro_F_3 | 13.2 (89.5%) | 11762091<br>(80.0%) | 1052235<br>(7.2%) | 348850<br>(2.4%) | 922849<br>(6.3%) | 618591<br>(4.2%) |
| Dpro_F_4 | 12.8 (86.9%) | 11607811<br>(78.6%) | 876462<br>(5.9%) | 352797<br>(2.4%) | 961721<br>(6.5%) | 974142<br>(6.6%) |
| Dpro_M_1 | 12.7 (92.3%) | 11382212<br>(82.9%) | 882537<br>(6.4%) | 416154<br>(3.0%) | 909163<br>(6.6%) | 145253<br>(1.1%) |
| Dpro_M_2 | 12.9 (92.2%) | 11595790<br>(83.2%) | 849273<br>(6.1%) | 407331<br>(2.9%) | 924248<br>(6.6%) | 167338<br>(1.2%) |
| Dpro_M_3 | 11.3 (85.8%) | 10172545<br>(76.9%) | 812473<br>(6.1%) | 354630<br>(2.7%) | 848611<br>(6.4%) | 1033239<br>(7.8%) |
| Dpro_M_5 | 3.9 (89.3%) | 3468903<br>(79.9%) | 268893<br>(6.2%) | 137750<br>(3.2%) | 348931<br>(8.0%) | 114818<br>(2.6%) |
| Dcar_F_1 | 13.2 (95.5%) | 12195693<br>(88.0%) | 663784<br>(4.8%) | 375294<br>(2.7%) | 460058<br>(3.3%) | 166504<br>(1.2%) |
| Dcar_F_2 | 13.4 (96.4%) | 12414444<br>(89.5%) | 603605<br>(4.4%) | 349775<br>(2.5%) | 367662<br>(2.7%) | 132232<br>(1.0%) |
| Dcar_F_4 | 12.9 (95.7%) | 11847580<br>(87.8%) | 670845<br>(5.0%) | 385196<br>(2.9%) | 438529<br>(3.3%) | 148504<br>(1.1%) |
| Dcar_F_6 | 14.4 (91.2%) | 13315646<br>(84.0%) | 738847<br>(4.7%) | 395172<br>(2.5%) | 441754<br>(2.8%) | 951881<br>(6.0%) |
| Dcar_M_1 | 12 (96.2%) | 11039955<br>(88.6%) | 514678<br>(4.1%) | 435143<br>(3.5%) | 362917<br>(2.9%) | 110586<br>(0.9%) |
| Dcar_M_2 | 13 (94.0%) | 11949586<br>(86.2%) | 636828<br>(4.6%) | 448819<br>(3.2%) | 367462<br>(2.7%) | 459323<br>(3.3%) |
| Dcar_M_3 | 12.6 (96.2%) | 11563218<br>(88.6%) | 591867<br>(4.5%) | 401992<br>(3.1%) | 373925<br>(2.9%) | 124977<br>(1.0%) |
| Dcar_M_5 | 10.2 (91.3%) | 9297876<br>(83.3%) | 580338<br>(5.2%) | 307213<br>(2.8%) | 348561<br>(3.1%) | 624109<br>(5.6%) |

Ambiguous features: reads that overlap with two or more features
Multi mapping: reads that map to more than one location in the genome

**Table S9.**
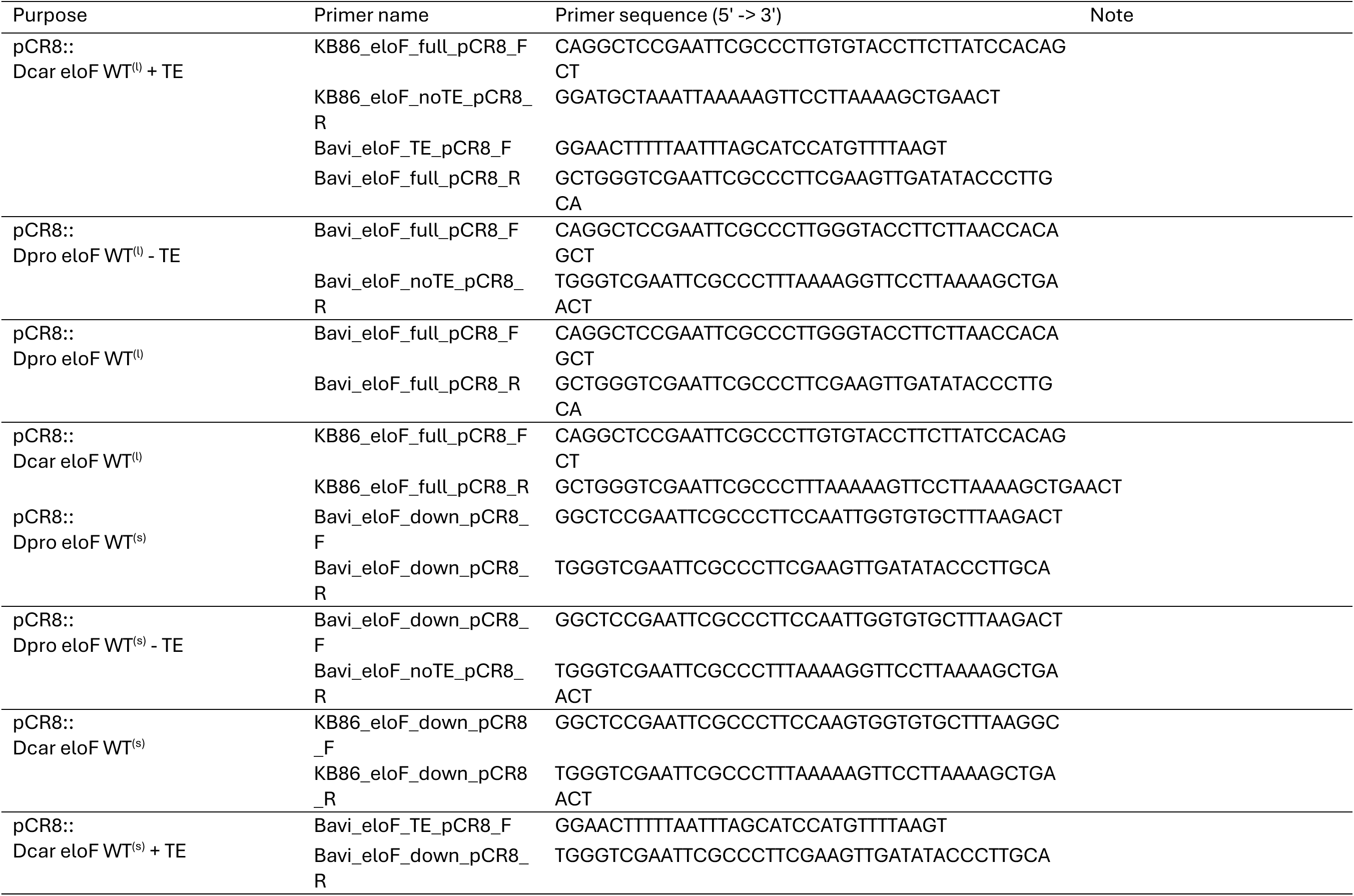
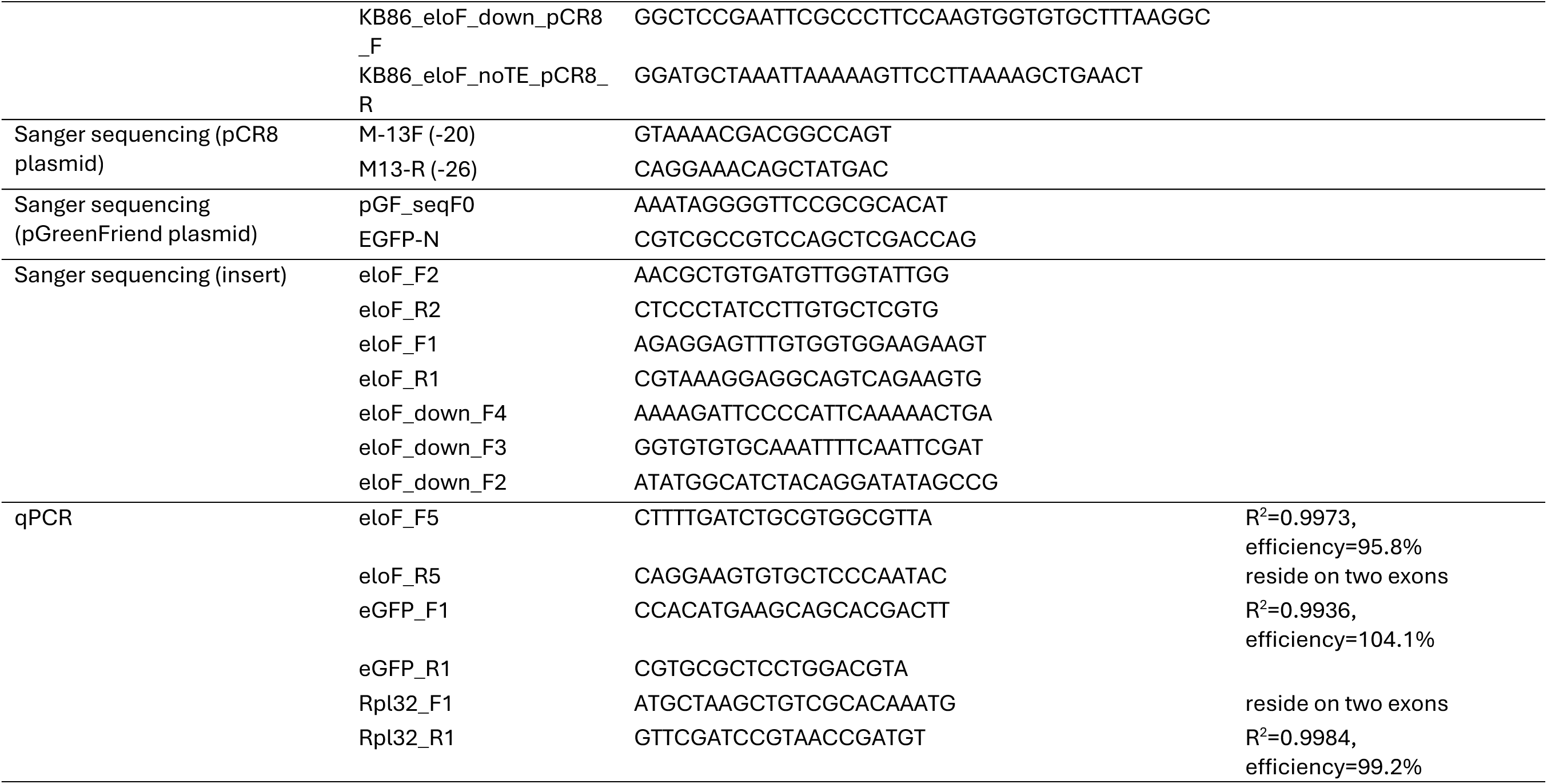
Primers used for cloning and qPCR.

## References

Ackermann M, Strimmer K. 2009. A general modular framework for gene set enrichment analysis. BMC Bioinformatics. 10(1):47. doi:10.1186/1471-2105-10-47.

Adams TS, Dillwith JW, Blomquist GJ. 1984. The role of 20-hydroxyecdysone in housefly sex pheromone biosynthesis. J Insect Physiol. 30(4):287–294. doi:10.1016/0022-1910(84)90129-X.

Alexa A, Rahnenfuhrer J. 2022. topGO: Enrichment Analysis for Gene Ontology. doi:10.18129/B9.bioc.topGO. [accessed 2022 Nov 1]. https://bioconductor.org/packages/topGO/.

Alexa A, Rahnenführer J, Lengauer T. 2006. Improved scoring of functional groups from gene expression data by decorrelating GO graph structure. Bioinformatics. 22(13):1600–1607. doi:10.1093/bioinformatics/btl140.

Almojil D, Bourgeois Y, Falis M, Hariyani I, Wilcox J, Boissinot S. 2021. The Structural, Functional and Evolutionary Impact of Transposable Elements in Eukaryotes. Genes. 12(6):918. doi:10.3390/genes12060918.

Amino K, Matsuo T. 2020. Intra-Versus Inter-Sexual Selection on Sexually Dimorphic Traits in Drosophila prolongata. Zoolog Sci. 37(3):210–216. doi:10.2108/zs200010.

Andersson M. 1994. Sexual selection. Princeton, NJ: Princeton Univ. Press.

Bailey TL, Elkan C. 1994. Fitting a mixture model by expectation maximization to discover motifs in biopolymers. Proc Int Conf Intell Syst Mol Biol. 2:28–36.

Baron A, Denis B, Wicker-Thomas C. 2018. Control of pheromone production by ovaries in Drosophila. J Insect Physiol. 109:138–143. doi:10.1016/j.jinsphys.2018.07.003.

Bartelt RJ, Schaner AM, Jackson LL. 1985. cis-Vaccenyl acetate as an aggregation pheromone inDrosophila melanogaster. J Chem Ecol. 11(12):1747–1756. doi:10.1007/BF01012124.

Barth NKH, Li L, Taher L. 2020. Independent Transposon Exaptation Is a Widespread Mechanism of Redundant Enhancer Evolution in the Mammalian Genome. Genome Biol Evol. 12(3):1–17. doi:10.1093/gbe/evaa004.

Benjamini Y, Hochberg Y. 1995. Controlling the False Discovery Rate: A Practical and Powerful Approach to Multiple Testing. J R Stat Soc Ser B Methodol. 57(1):289–300.

Bergman CM, Carlson JW, Celniker SE. 2005. Drosophila DNase I footprint database: a systematic genome annotation of transcription factor binding sites in the fruitfly, Drosophila melanogaster. Bioinforma Oxf Engl. 21(8):1747–1749. doi:10.1093/bioinformatics/bti173.

Bergman DT, Jones TR, Liu V, Ray J, Jagoda E, Siraj L, Kang HY, Nasser J, Kane M, Rios A, et al. 2022. Compatibility rules of human enhancer and promoter sequences. Nature. 607(7917):176–184. doi:10.1038/s41586-022-04877-w.

Bi J, Xiang Y, Chen H, Liu Z, Grönke S, Kühnlein RP, Huang X. 2012. Opposite and redundant roles of the two Drosophila perilipins in lipid mobilization. J Cell Sci. 125(15):3568–3577. doi:10.1242/jcs.101329.

Billeter J-C, Atallah J, Krupp JJ, Millar JG, Levine JD. 2009. Specialized cells tag sexual and species identity in Drosophila melanogaster. Nature. 461(7266):987–991. doi:10.1038/nature08495.

Blomquist GJ, Adams TS, Dillwith JW. 1984. Induction of female sex pheromone production in male houseflies by ovary implants or 20-hydroxyecdysone. J Insect Physiol. 30(4):295–302. doi:10.1016/0022-1910(84)90130-6.

Blomquist GJ, Adams TS, Halarnkar PP, Gu P, Mackay ME, Brown LA. 1992. Ecdysteroid induction of sex pheromone biosynthesis in the housefly, Musca domestica—Are other factors involved? J Insect Physiol. 38(4):309–318. doi:10.1016/0022-1910(92)90131-V.

Blomquist GJ, Bagnères A-G. 2010. Insect hydrocarbons biology, biochemistry, and chemical ecology. Cambridge, UK: Cambridge Univ. Press.

Blomquist GJ, Tillman JA, Reed JR, Gu P, Vanderwel D, Choi S, Reitz RC. 1995. Regulation of enzymatic activity involved in sex pheromone production in the housefly, Musca domestica. Insect Biochem Mol Biol. 25(6):751–757. doi:10.1016/0965-1748(95)00015-N.

Bontonou G, Wicker-Thomas C. 2014. Sexual Communication in the Drosophila Genus. Insects. 5(2):439–458. doi:10.3390/insects5020439.

Boughman JW, Rundle HD, Schluter D. 2005. Parallel evolution of sexual isolation in sticklebacks. Evolution. 59(2):361–373. doi:10.1111/j.0014-3820.2005.tb00995.x.

Bourbon H-MG, Benetah MH, Guillou E, Mojica-Vazquez LH, Baanannou A, Bernat-Fabre S, Loubiere V, Bantignies F, Cavalli G, Boube M. 2022. A shared ancient enhancer element differentially regulates the bric-a-brac tandem gene duplicates in the developing Drosophila leg. PLOS Genet. 18(3):e1010083. doi:10.1371/journal.pgen.1010083.

Bourque G, Leong B, Vega VB, Chen X, Lee YL, Srinivasan KG, Chew J-L, Ruan Y, Wei C-L, Ng HH, et al. 2008. Evolution of the mammalian transcription factor binding repertoire via transposable elements. Genome Res. 18(11):1752–1762. doi:10.1101/gr.080663.108.

Bray S, Amrein H. 2003. A Putative Drosophila Pheromone Receptor Expressed in Male-Specific Taste Neurons Is Required for Efficient Courtship. Neuron. 39(6):1019–1029. doi:10.1016/S0896-6273(03)00542-7.

Broder ED, Elias DO, Rodríguez RL, Rosenthal GG, Seymoure BM, Tinghitella RM. 2021. Evolutionary novelty in communication between the sexes. Biol Lett. 17(2):20200733. doi:10.1098/rsbl.2020.0733.

Buchinger TJ, Li W. 2023. Chemical communication and its role in sexual selection across Animalia. Commun Biol. 6(1):1178. doi:10.1038/s42003-023-05572-w.

Campbell PM, Healy MJ, Oakeshott JG. 1992. Characterisation of juvenile hormone esterase in Drosophila melanogaster. Insect Biochem Mol Biol. 22(7):665–677. doi:10.1016/0965-1748(92)90045-G.

Cantarel BL, Korf I, Robb SMC, Parra G, Ross E, Moore B, Holt C, Alvarado AS, Yandell M. 2008. MAKER: An easy-to-use annotation pipeline designed for emerging model organism genomes. Genome Res. 18(1):188–196. doi:10.1101/gr.6743907.

Carroll SB. 2008. Evo-Devo and an Expanding Evolutionary Synthesis: A Genetic Theory of Morphological Evolution. Cell. 134(1):25–36. doi:10.1016/j.cell.2008.06.030.

Chertemps T, Duportets L, Labeur C, Ueda R, Takahashi K, Saigo K, Wicker-Thomas C. 2007. A female-biased expressed elongase involved in long-chain hydrocarbon biosynthesis and courtship behavior in Drosophila melanogaster. Proc Natl Acad Sci. 104(11):4273–4278. 10.1073/pnas.0608142104.

Chertemps T, Duportets L, Labeur C, Ueyama M, Wicker-Thomas C. 2006. A female-specific desaturase gene responsible for diene hydrocarbon biosynthesis and courtship behaviour in Drosophila melanogaster. Insect Mol Biol. 15(4):465–473. 10.1111/j.1365-2583.2006.00658.x.

Chiang YN, Tan KJ, Chung H, Lavrynenko O, Shevchenko A, Yew JY. 2016. Steroid Hormone Signaling Is Essential for Pheromone Production and Oenocyte Survival. PLOS Genet. 12(6):e1006126. doi:10.1371/journal.pgen.1006126.

Chung H, Carroll SB. 2015. Wax, sex and the origin of species: Dual roles of insect cuticular hydrocarbons in adaptation and mating. Bioessays. 37(7):822–830. doi:10.1002/bies.201500014.

Chung H, Loehlin DW, Dufour HD, Vaccarro K, Millar JG, Carroll SB. 2014. A Single Gene Affects Both Ecological Divergence and Mate Choice in *Drosophila*. Science. 343(6175):1148–1151. doi:10.1126/science.1249998.

Chuong EB, Elde NC, Feschotte C. 2017. Regulatory activities of transposable elements: from conflicts to benefits. Nat Rev Genet. 18(2):71–86. doi:10.1038/nrg.2016.139.

Cinnamon E, Makki R, Sawala A, Wickenberg LP, Blomquist GJ, Tittiger C, Paroush Z, Gould AP. 2016. Drosophila Spidey/Kar Regulates Oenocyte Growth via PI3-Kinase Signaling. PLoS Genet. 12(8):e1006154. doi:10.1371/journal.pgen.1006154.

Combs PA, Krupp JJ, Khosla NM, Bua D, Petrov DA, Levine JD, Fraser HB. 2018. Tissue-specific cis-regulatory divergence implicates eloF in inhibiting interspecies mating in Drosophila. Curr Biol. 28(24):3969–3975. doi:10.1016/j.cub.2018.10.036.

Course MM, Scott AI, Schoor C, Hsieh C-H, Papakyrikos AM, Winter D, Cowan TM, Wang X. 2018. Phosphorylation of MCAD selectively rescues PINK1 deficiencies in behavior and metabolism. Mol Biol Cell. 29(10):1219–1227. doi:10.1091/mbc.E18-03-0155.

Cridland JM, Thornton KR, Long AD. 2015. Gene Expression Variation in Drosophila melanogaster Due to Rare Transposable Element Insertion Alleles of Large Effect. Genetics. 199(1):85–93. doi:10.1534/genetics.114.170837.

Dallerac R, Labeur C, Jallon J-M, Knipple DC, Roelofs WL, Wicker-Thomas C. 2000. A Δ9 desaturase gene with a different substrate specificity is responsible for the cuticular diene hydrocarbon polymorphism in Drosophila melanogaster. Proc Natl Acad Sci. 97(17):9449–9454. doi:10.1073/pnas.150243997.

Datta SR, Vasconcelos ML, Ruta V, Luo S, Wong A, Demir E, Flores J, Balonze K, Dickson BJ, Axel R. 2008. The Drosophila pheromone cVA activates a sexually dimorphic neural circuit. Nature. 452(7186):473–477. doi:10.1038/nature06808.

Dembeck LM, Böröczky K, Huang W, Schal C, Anholt RR, Mackay TF. 2015. Genetic architecture of natural variation in cuticular hydrocarbon composition in Drosophila melanogaster. eLife. 4:e09861. 10.7554/eLife.09861.001.

Dobin A, Davis CA, Schlesinger F, Drenkow J, Zaleski C, Jha S, Batut P, Chaisson M, Gingeras TR. 2013. STAR: ultrafast universal RNA-seq aligner. Bioinforma Oxf Engl. 29(1):15–21. doi:10.1093/bioinformatics/bts635.

Drăghici S, Sellamuthu S, Khatri P. 2006. Babel’s tower revisited: A universal resource for cross-referencing across annotation databases. Bioinforma Oxf Engl. 22(23):2934–2939. doi:10.1093/bioinformatics/btl372.

Everaerts C, Farine J-P, Cobb M, Ferveur J-F. 2010. Drosophila Cuticular Hydrocarbons Revisited: Mating Status Alters Cuticular Profiles. PloS One. 5(3):e9607. 10.1371/journal.pone.0009607.

Fan P, Manoli DS, Ahmed OM, Chen Y, Agarwal N, Kwong S, Cai AG, Neitz J, Renslo A, Baker BS, et al. 2013. Genetic and neural mechanisms that inhibit Drosophila from mating with other species. Cell. 154(1):89–102. 10.1016/j.cell.2013.06.008.

Fang S, Takahashi A, Wu C-I. 2002. A mutation in the promoter of desaturase 2 is correlated with sexual isolation between Drosophila behavioral races. Genetics. 162(2):781–784. doi:10.1093/genetics/162.2.781.

Ferveur J-F. 2005. Cuticular hydrocarbons: their evolution and roles in Drosophila pheromonal communication. Behav Genet. 35(3):279–295. 10.1007/s10519-005-3220-5.

Ferveur J-F, Cobb M, Boukella H, Jallon J-M. 1996. World-wide variation in Drosophila melanogaster sex pheromone: behavioural effects, genetic bases and potential evolutionary consequences. Genetica. 97(1):73–80. doi:10.1007/BF00132583.

Ferveur J-F, Jallon J-M. 1996. Genetic control of male cuticular hydrocarbons in Drosophila melanogaster. Genet Res. 67(3):211–218. doi:10.1017/S0016672300033693.

Ferveur J-F, Savarit F, O’Kane CJ, Sureau G, Greenspan RJ, Jallon J-M. 1997. Genetic feminization of pheromones and its behavioral consequences in Drosophila males. Science. 276(5318):1555–1558. 10.1126/science.276.5318.1555.

Ferveur J-F, Sureau G. 1996. Simultaneous influence on male courtship of stimulatory and inhibitory pheromones produced by live sex-mosaic Drosophila melanogaster. Proc R Soc Lond B Biol Sci. 263(1373):967–973. doi:10.1098/rspb.1996.0143.

Finet C, Slavik K, Pu J, Carroll SB, Chung H. 2019. Birth-and-Death Evolution of the Fatty Acyl-CoA Reductase (FAR) Gene Family and Diversification of Cuticular Hydrocarbon Synthesis in Drosophila. Genome Biol Evol. 11(6):1541–1551. doi:10.1093/gbe/evz094.

González J, Macpherson JM, Petrov DA. 2009. A Recent Adaptive Transposable Element Insertion Near Highly Conserved Developmental Loci in Drosophila melanogaster. Mol Biol Evol. 26(9):1949–1961. doi:10.1093/molbev/msp107.

Gordon HB, Valdez L, Letsou A. 2018. Etiology and treatment of adrenoleukodystrophy: new insights from Drosophila. Dis Model Mech. 11(6):dmm031286. doi:10.1242/dmm.031286.

Gould A, Morrison A, Sproat G, White RA, Krumlauf R. 1997. Positive cross-regulation and enhancer sharing: two mechanisms for specifying overlapping Hox expression patterns. Genes Dev. 11(7):900– 913. doi:10.1101/gad.11.7.900.

Grant CE, Bailey TL, Noble WS. 2011. FIMO: scanning for occurrences of a given motif. Bioinformatics. 27(7):1017–1018. doi:10.1093/bioinformatics/btr064.

Gratten J, Beraldi D, Lowder B v, McRae A f, Visscher P m, Pemberton J m, Slate J. 2007. Compelling evidence that a single nucleotide substitution in TYRP1 is responsible for coat-colour polymorphism in a free-living population of Soay sheep. Proc R Soc B Biol Sci. 274(1610):619–626. doi:10.1098/rspb.2006.3762.

Grillet M, Dartevelle L, Ferveur J-F. 2006. A Drosophila male pheromone affects female sexual receptivity. Proc R Soc Lond B Biol Sci. 273(1584):315–323. doi:10.1098/rspb.2005.3332.

Grillet M, Everaerts C, Houot B, Ritchie MG, Cobb M, Ferveur J-F. 2012. Incipient speciation in Drosophila melanogaster involves chemical signals. Sci Rep. 2(1):224. doi:10.1038/srep00224.

Herpin A, Schartl M, Depincé A, Guiguen Y, Bobe J, Hua-Van A, Hayman ES, Octavera A, Yoshizaki G, Nichols KM, et al. 2021. Allelic diversification after transposable element exaptation promoted gsdf as the master sex determining gene of sablefish. Genome Res. doi:10.1101/gr.274266.120. [accessed 2021 Aug 9]. https://genome.cshlp.org/content/early/2021/07/23/gr.274266.120.

Hof AE van’t, Campagne P, Rigden DJ, Yung CJ, Lingley J, Quail MA, Hall N, Darby AC, Saccheri IJ. 2016. The industrial melanism mutation in British peppered moths is a transposable element. Nature. 534(7605):102–105. doi:10.1038/nature17951.

Hopkins BR, Kopp A. 2021. Evolution of sexual development and sexual dimorphism in insects. Curr Opin Genet Dev. 69:129–139. doi:10.1016/j.gde.2021.02.011.

Howard RW, Blomquist GJ. 2005. Ecological, behavioral, and biochemical aspects of insect hydrocarbons. Annu Rev Entomol. 50(1):371–393. 10.1146/annurev.ento.50.071803.130359.

Howard RW, Jackson LL, Banse H, Blows MW. 2003. Cuticular hydrocarbons of Drosophila birchii and D. serrata: identification and role in mate choice in D. serrata. J Chem Ecol. 29(4):961–976. 10.1023/A:1022992002239.

Ishii K, Hirai Y, Katagiri C, Kimura MT. 2001. Sexual isolation and cuticular hydrocarbons in Drosophila elegans. Heredity. 87(4):392–399. doi:10.1046/j.1365-2540.2001.00864.x.

Jallon J-M. 1984. A few chemical words exchanged by Drosophila during courtship and mating. Behav Genet. 14(5):441–478. 10.1007/BF01065444.

Jallon JM, Antony C, Benamar O. 1981. An anti-aphrodisiac produced by Drosophila-Melanogaster males and transferred to females during copulation. C R Acad Sci Ser III Sci. Vie 292:1147–1149.

Jiggins CD, Estrada C, Rodrigues A. 2004. Mimicry and the evolution of premating isolation in Heliconius melpomene Linnaeus. J Evol Biol. 17(3):680–691. doi:10.1111/j.1420-9101.2004.00675.x.

Kapitonov VV, Jurka J. 2003. Molecular paleontology of transposable elements in the *Drosophila melanogaster* genome. Proc Natl Acad Sci. 100(11):6569–6574. doi:10.1073/pnas.0732024100.

Khallaf MA, Auer TO, Grabe V, Depetris-Chauvin A, Ammagarahalli B, Zhang D-D, Lavista-Llanos S, Kaftan F, Weißflog J, Matzkin LM, et al. 2020. Mate discrimination among subspecies through a conserved olfactory pathway. Sci Adv. 6(25):eaba5279. doi:10.1126/sciadv.aba5279.

Khallaf MA, Cui R, Weißflog J, Erdogmus M, Svatoš A, Dweck HKM, Valenzano DR, Hansson BS, Knaden M. 2021. Large-scale characterization of sex pheromone communication systems in Drosophila. Nat Commun. 12(1):4165. doi:10.1038/s41467-021-24395-z.

Kim BY, Wang JR, Miller DE, Barmina O, Delaney E, Thompson A, Comeault AA, Peede D, D’Agostino ER, Pelaez J, et al. 2021. Highly contiguous assemblies of 101 drosophilid genomes. Coop G, Wittkopp PJ, Sackton TB, editors. eLife. 10:e66405. doi:10.7554/eLife.66405.

King-Jones K, Charles J-P, Lam G, Thummel CS. 2005. The Ecdysone-Induced DHR4 Orphan Nuclear Receptor Coordinates Growth and Maturation in Drosophila. Cell. 121(5):773–784. doi:10.1016/j.cell.2005.03.030.

Kishita Y, Tsuda M, Aigaki T. 2012. Impaired fatty acid oxidation in a Drosophila model of mitochondrial trifunctional protein (MTP) deficiency. Biochem Biophys Res Commun. 419(2):344–349. doi:10.1016/j.bbrc.2012.02.026.

Kopp A. 2012. Dmrt genes in the development and evolution of sexual dimorphism. Trends Genet. 28(4):175–184. doi:10.1016/j.tig.2012.02.002.

Kopp A, Duncan I, Carroll SB. 2000. Genetic control and evolution of sexually dimorphic characters in Drosophila. Nature. 408(6812):553–559. doi:10.1038/35046017.

Kudo A, Shigenobu S, Kadota K, Nozawa M, Shibata TF, Ishikawa Y, Matsuo T. 2017. Comparative analysis of the brain transcriptome in a hyper-aggressive fruit fly, Drosophila prolongata. Insect Biochem Mol Biol. 82:11–20. doi:10.1016/j.ibmb.2017.01.006.

Kudo A, Takamori H, Watabe H, Ishikawa Y, Matsuo T. 2015. Variation in morphological and behavioral traits among isofemale strains of Drosophila prolongata (Diptera: Drosophilidae). Entomol Sci. 18(2):221–229. doi:10.1111/ens.12116.

Kulakovskiy IV, Makeev VJ. 2009. Discovery of DNA motifs recognized by transcription factors through integration of different experimental sources. Biophysics. 54(6):667–674. doi:10.1134/S0006350909060013.

Kurtovic A, Widmer A, Dickson BJ. 2007. A single class of olfactory neurons mediates behavioural responses to a Drosophila sex pheromone. Nature. 446(7135):542–546. doi:10.1038/nature05672.

Kwon D, Mucci D, Langlais KK, Americo JL, DeVido SK, Cheng Y, Kassis JA. 2009. Enhancer-promoter communication at the Drosophila engrailedlocus. Development. 136(18):3067–3075. doi:10.1242/dev.036426.

Larkin A, Marygold SJ, Antonazzo G, Attrill H, Dos Santos G, Garapati PV, Goodman JL, Gramates LS, Millburn G, Strelets VB, et al. 2021. FlyBase: updates to the Drosophila melanogaster knowledge base. Nucleic Acids Res. 49(D1):D899–D907. doi:10.1093/nar/gkaa1026.

Lassance J-M, Groot AT, Liénard MA, Antony B, Borgwardt C, Andersson F, Hedenström E, Heckel DG, Löfstedt C. 2010. Allelic variation in a fatty-acyl reductase gene causes divergence in moth sex pheromones. Nature. 466(7305):486–489. doi:10.1038/nature09058.

Laturney M, Billeter J-C. 2016. Drosophila melanogaster females restore their attractiveness after mating by removing male anti-aphrodisiac pheromones. Nat Commun. 7(1):12322. doi:10.1038/ncomms12322.

Laturney M, Moehring AJ. 2012. The Genetic Basis of Female Mate Preference and Species Isolation in Drosophila. Int J Evol Biol. 2012:e328392. doi:10.1155/2012/328392.

Law CW, Chen Y, Shi W, Smyth GK. 2014. voom: precision weights unlock linear model analysis tools for RNA-seq read counts. Genome Biol. 15(2):R29. doi:10.1186/gb-2014-15-2-r29.

Lawniczak MKN, Barnes AI, Linklater JR, Boone JM, Wigby S, Chapman T. 2007. Mating and immunity in invertebrates. Trends Ecol Evol. 22(1):48–55. doi:10.1016/j.tree.2006.09.012.

Li H, Janssens J, Waegeneer MD, Kolluru SS, Davie K, Gardeux V, Saelens W, David F, Brbić M, Leskovec J, et al. 2021. Fly Cell Atlas: a single-cell transcriptomic atlas of the adult fruit fly. [accessed 2021 Sep 1]. https://www.biorxiv.org/content/10.1101/2021.07.04.451050v1.

Liénard MA, Hagström ÅK, Lassance J-M, Löfstedt C. 2010. Evolution of multicomponent pheromone signals in small ermine moths involves a single fatty-acyl reductase gene. Proc Natl Acad Sci. 107(24):10955–10960. doi:10.1073/pnas.1000823107.

Linn CE, Young MS, Gendle M, Glover TJ, Roelofs WL. 1997. Sex pheromone blend discrimination in two races and hybrids of the European corn borer moth, Ostrinia nubilalis. Physiol Entomol. 22(3):212–223. doi:10.1111/j.1365-3032.1997.tb01161.x.

Livak KJ, Schmittgen TD. 2001. Analysis of relative gene expression data using real-time quantitative PCR and the 2(-Delta Delta C(T)) Method. Methods San Diego Calif. 25(4):402–408. doi:10.1006/meth.2001.1262.

Loker R, Mann RS. 2022. Divergent expression of paralogous genes by modification of shared enhancer activity through a promoter-proximal silencer. Curr Biol CB. 32(16):3545–3555.e4. doi:10.1016/j.cub.2022.06.069.

Luecke D, Luo Y, Krzystek H, Jones C, Kopp A. 2024. Highly contiguous genome assembly of Drosophila prolongata—a model for evolution of sexual dimorphism and male-specific innovations. G3 GenesGenomesGenetics.:jkae155. doi:10.1093/g3journal/jkae155.

Luecke D, Rice G, Kopp A. 2022. Sex-specific evolution of a Drosophila sensory system via interacting cis- and trans-regulatory changes. Evol Dev. 24(1–2):37–60. doi:10.1111/ede.12398.

Luecke DM, Kopp A. 2019. Sex-specific evolution of relative leg size in Drosophila prolongata results from changes in the intersegmental coordination of tissue growth. Evolution. 73(11):2281–2294. doi:10.1111/evo.13847.

Luo Y, Zhang Y, Farine J-P, Ferveur J-F, Ramírez S, Kopp A. 2019. Evolution of sexually dimorphic pheromone profiles coincides with increased number of male-specific chemosensory organs in Drosophila prolongata. Ecol Evol. 9(23):13608–13618. doi:10.1002/ece3.5819.

Lynch VJ, Leclerc RD, May G, Wagner GP. 2011. Transposon-mediated rewiring of gene regulatory networks contributed to the evolution of pregnancy in mammals. Nat Genet. 43(11):1154–1159. doi:10.1038/ng.917.

Makki R, Cinnamon E, Gould AP. 2014. The Development and Functions of Oenocytes. Annu Rev Entomol. 59(1):405–425. doi:10.1146/annurev-ento-011613-162056.

Markstein M, Pitsouli C, Villalta C, Celniker SE, Perrimon N. 2008. Exploiting position effects and the gypsy retrovirus insulator to engineer precisely expressed transgenes. Nat Genet. 40(4):476–483. doi:10.1038/ng.101.

McLeay RC, Bailey TL. 2010. Motif Enrichment Analysis: a unified framework and an evaluation on ChIP data. BMC Bioinformatics. 11(1):165. doi:10.1186/1471-2105-11-165.

Miller SW, Rebeiz M, Atanasov JE, Posakony JW. 2014. Neural precursor-specific expression of multiple Drosophila genes is driven by dual enhancer modules with overlapping function. Proc Natl Acad Sci U S A. 111(48):17194–17199. doi:10.1073/pnas.1415308111.

Minekawa K, Amino K, Matsuo T. 2020. A courtship behavior that makes monandrous females polyandrous. Evolution. 74(11):2483–2493. doi:10.1111/evo.14098.

Monnin T. 2006. Chemical recognition of reproductive status in social insects. In: Annales Zoologici Fennici. JSTOR. p. 515–530. http://www.jstor.org/stable/23736759.

Museridze M, Ceolin S, Mühling B, Ramanathan S, Barmina O, Sekhar PS, Gompel N. 2024. Entangled and non-modular enhancer sequences producing independent spatial activities.: 2024.07.08.602541. doi:10.1101/2024.07.08.602541. [accessed 2024 Sep 13]. https://www.biorxiv.org/content/10.1101/2024.07.08.602541v1.

Nachman MW, Hoekstra HE, D’Agostino SL. 2003. The genetic basis of adaptive melanism in pocket mice. Proc Natl Acad Sci. 100(9):5268–5273. doi:10.1073/pnas.0431157100.

Ng SH, Shankar S, Shikichi Y, Akasaka K, Mori K, Yew JY. 2014. Pheromone evolution and sexual behavior in Drosophila are shaped by male sensory exploitation of other males. Proc Natl Acad Sci. 111(8):3056– 3061. 10.1073/pnas.1313615111.

Ohtsuki S, Levine M, Cai HN. 1998. Different core promoters possess distinct regulatory activities in the Drosophila embryo. Genes Dev. 12(4):547–556.

Pei X-J, Fan Y-L, Bai Y, Bai T-T, Schal C, Zhang Z-F, Chen N, Li S, Liu T-X. 2021. Modulation of fatty acid elongation in cockroaches sustains sexually dimorphic hydrocarbons and female attractiveness. PLOS Biol. 19(7):e3001330. doi:10.1371/journal.pbio.3001330.

Petersen KR, Streett DA, Gerritsen AT, Hunter SS, Settles ML. 2015. Super deduper, fast PCR duplicate detection in fastq files. In: Proceedings of the 6th ACM Conference on Bioinformatics, Computational Biology and Health Informatics. New York, NY, USA: Association for Computing Machinery. (BCB‘15). p. 491–492. [accessed 2022 Dec 6]. 10.1145/2808719.2811568.

Pfaffl MW. 2001. A new mathematical model for relative quantification in real-time RT–PCR. Nucleic Acids Res. 29(9):e45. doi:10.1093/nar/29.9.e45.

Ponton F, Chapuis M-P, Pernice M, Sword GA, Simpson SJ. 2011. Evaluation of potential reference genes for reverse transcription-qPCR studies of physiological responses in Drosophila melanogaster. J Insect Physiol. 57(6):840–850. doi:10.1016/j.jinsphys.2011.03.014.

Qiu Y, Tittiger C, Wicker-Thomas C, Le Goff G, Young S, Wajnberg E, Fricaux T, Taquet N, Blomquist GJ, Feyereisen R. 2012. An insect-specific P450 oxidative decarbonylase for cuticular hydrocarbon biosynthesis. Proc Natl Acad Sci U S A. 109(37):14858–14863. doi:10.1073/pnas.1208650109.

Ritchie ME, Phipson B, Wu D, Hu Y, Law CW, Shi W, Smyth GK. 2015. limma powers differential expression analyses for RNA-sequencing and microarray studies. Nucleic Acids Res. 43(7):e47. doi:10.1093/nar/gkv007.

Robinson MD, McCarthy DJ, Smyth GK. 2010. edgeR: a Bioconductor package for differential expression analysis of digital gene expression data. Bioinformatics. 26(1):139–140. doi:10.1093/bioinformatics/btp616.

Robinson MD, Oshlack A. 2010. A scaling normalization method for differential expression analysis of RNA-seq data. Genome Biol. 11(3):R25. doi:10.1186/gb-2010-11-3-r25.

Rogers WA, Salomone JR, Tacy DJ, Camino EM, Davis KA, Rebeiz M, Williams TM. 2013. Recurrent Modification of a Conserved Cis-Regulatory Element Underlies Fruit Fly Pigmentation Diversity. PLOS Genet. 9(8):e1003740. doi:10.1371/journal.pgen.1003740.

Rusuwa BB, Chung H, Allen SL, Frentiu FD, Chenoweth SF. 2022. Natural variation at a single gene generates sexual antagonism across fitness components in Drosophila. Curr Biol. 32(14):3161–3169.e7. doi:10.1016/j.cub.2022.05.038.

Ruta V, Datta SR, Vasconcelos ML, Freeland J, Looger LL, Axel R. 2010. A dimorphic pheromone circuit in Drosophila from sensory input to descending output. Nature. 468(7324):686–690. doi:10.1038/nature09554.

Schaefer HM, Ruxton GD. 2015. Signal Diversity, Sexual Selection, and Speciation. Annu Rev Ecol Evol Syst. 46(Volume 46, 2015):573–592. doi:10.1146/annurev-ecolsys-112414-054158.

Schindelin J, Arganda-Carreras I, Frise E, Kaynig V, Longair M, Pietzsch T, Preibisch S, Rueden C, Saalfeld S, Schmid B, et al. 2012. Fiji: an open-source platform for biological-image analysis. Nat Methods. 9(7):676–682. doi:10.1038/nmeth.2019.

Scott Adams P. 2007. Data analysis and reporting. Taylor & Francis. [accessed 2022 Dec 6]. https://www.taylorfrancis.com/chapters/edit/10.4324/9780203967317-11/data-analysis-reporting-pamela-scott-adams.

Seeholzer LF, Seppo M, Stern DL, Ruta V. 2018. Evolution of a central neural circuit underlies Drosophila mate preferences. Nature. 559(7715):564–569. doi:10.1038/s41586-018-0322-9.

Senft AD, Macfarlan TS. 2021. Transposable elements shape the evolution of mammalian development. Nat Rev Genet. 22(11):691–711. doi:10.1038/s41576-021-00385-1.

Setoguchi S, Kudo A, Takanashi T, Ishikawa Y, Matsuo T. 2015. Social context-dependent modification of courtship behaviour in Drosophila prolongata. Proc R Soc Lond B Biol Sci. 282(1818):20151377. 10.1098/rspb.2015.1377.

Setoguchi S, Takamori H, Aotsuka T, Sese J, Ishikawa Y, Matsuo T. 2014. Sexual dimorphism and courtship behavior in Drosophila prolongata. J Ethol. 32(2):91–102. 10.1007/s10164-014-0399-z.

Shirangi TR, Dufour HD, Williams TM, Carroll SB. 2009. Rapid evolution of sex pheromone-producing enzyme expression in Drosophila. PLoS Biol. 7(8):e1000168. 10.1371/journal.pbio.1000168.

Shumate A, Salzberg SL. 2020. Liftoff: accurate mapping of gene annotations. Bioinforma Oxf Engl. 37(12):1639–1643. doi:10.1093/bioinformatics/btaa1016.

Sievers F, Wilm A, Dineen D, Gibson TJ, Karplus K, Li W, Lopez R, McWilliam H, Remmert M, Söding J, et al. 2011. Fast, scalable generation of high-quality protein multiple sequence alignments using Clustal Omega. Mol Syst Biol. 7:539. doi:10.1038/msb.2011.75.

Singh BK, Gupta JP. 1977. Two new and two unrecorded species of the genus Drosophila Fallen (Diptera: Drosophilidae) from Shillong, Meghalaya, India. Proc Zool Soc Calcutta. 30(1–2):31–38.

Smadja C, Butlin RK. 2008. On the scent of speciation: the chemosensory system and its role in premating isolation. Heredity. 102(1):77–97. doi:10.1038/hdy.2008.55.

Smyth GK. 2004. Linear Models and Empirical Bayes Methods for Assessing Differential Expression in Microarray Experiments. Stat Appl Genet Mol Biol. 3(1). doi:10.2202/1544-6115.1027. [accessed 2022 Nov 14]. https://www.degruyter.com/document/doi/10.2202/1544-6115.1027/html.

Steiger S, Stökl J. 2014. The role of sexual selection in the evolution of chemical signals in insects. Insects. 5(2):423–438. 10.3390/insects5020423.

Stökl J, Steiger S. 2017. Evolutionary origin of insect pheromones. Curr Opin Insect Sci. 24:36–42. doi:10.1016/j.cois.2017.09.004.

Storer J, Hubley R, Rosen J, Wheeler TJ, Smit AF. 2021. The Dfam community resource of transposable element families, sequence models, and genome annotations. Mob DNA. 12(1):2. doi:10.1186/s13100-020-00230-y.

Sundaram V, Cheng Y, Ma Z, Li D, Xing X, Edge P, Snyder MP, Wang T. 2014. Widespread contribution of transposable elements to the innovation of gene regulatory networks. Genome Res. 24(12):1963–1976. doi:10.1101/gr.168872.113.

Suzuki T. 1980. 4, 8-Dimethyldecanal: The aggregation pheromone of the flour beetles, Tribolium castaneum and T. confusum (Coleoptera: Tenebrionidae). Agric Biol Chem. 44(10):2519–2520.

Szafer-Glusman E, Giansanti MG, Nishihama R, Bolival B, Pringle J, Gatti M, Fuller MT. 2008. A Role for Very-Long-Chain Fatty Acids in Furrow Ingression during Cytokinesis in Drosophila Spermatocytes. Curr Biol. 18(18):1426–1431. doi:10.1016/j.cub.2008.08.061.

Takau A, Matsuo T. 2022. Contribution of visual stimuli to mating and fighting behaviors of Drosophila prolongata. Entomol Sci. 25(4):e12529. doi:10.1111/ens.12529.

Tanaka K, Barmina O, Kopp A. 2009. Distinct developmental mechanisms underlie the evolutionary diversification of Drosophila sex combs. Proc Natl Acad Sci. 106(12):4764–4769. doi:10.1073/pnas.0807875106.

Tena JJ, Alonso ME, de la Calle-Mustienes E, Splinter E, de Laat W, Manzanares M, Gómez-Skarmeta JL. 2011. An evolutionarily conserved three-dimensional structure in the vertebrate Irx clusters facilitates enhancer sharing and coregulation. Nat Commun. 2(1):310. doi:10.1038/ncomms1301.

Thome BL. 1982. Termite-termite interactions: workers as an agonistic caste. Psyche (Stuttg). 89(1–2):133–150.

Toda H, Zhao X, Dickson BJ. 2012. The Drosophila female aphrodisiac pheromone activates ppk23+ sensory neurons to elicit male courtship behavior. Cell Rep. 1(6):599–607. 10.1016/j.celrep.2012.05.007.

Toyoshima N, Matsuo T. 2023. Fight outcome influences male mating success in Drosophila prolongata. J Ethol. 41(2):119–127. doi:10.1007/s10164-023-00778-1.

Trizzino M, Park Y, Holsbach-Beltrame M, Aracena K, Mika K, Caliskan M, Perry GH, Lynch VJ, Brown CD. 2017. Transposable elements are the primary source of novelty in primate gene regulation. Genome Res. 27(10):1623–1633. doi:10.1101/gr.218149.116.

Tu Z. 1997. Three novel families of miniature inverted-repeat transposable elements are associated with genes of the yellow fever mosquito, Aedes aegypti. Proc Natl Acad Sci. 94(14):7475–7480. doi:10.1073/pnas.94.14.7475.

Wang L, Anderson DJ. 2010. Identification of an aggression-promoting pheromone and its receptor neurons in Drosophila. Nature. 463(7278):227–231.

Wang L, Han X, Mehren J, Hiroi M, Billeter J-C, Miyamoto T, Amrein H, Levine JD, Anderson DJ. 2011. Hierarchical chemosensory regulation of male-male social interactions in Drosophila. Nat Neurosci. 14(6):757–762. doi:10.1038/nn.2800.

Wang Z, Receveur JP, Pu J, Cong H, Richards C, Liang M, Chung H. 2022. Desiccation resistance differences in Drosophila species can be largely explained by variations in cuticular hydrocarbons. M. Riddiford L, editor. eLife. 11:e80859. doi:10.7554/eLife.80859.

Weill M, Lutfalla G, Mogensen K, Chandre F, Berthomieu A, Berticat C, Pasteur N, Philips A, Fort P, Raymond M. 2003. Insecticide resistance in mosquito vectors. Nature. 423(6936):136–137. doi:10.1038/423136b.

Wells MM, Henry CS. 1992. The role of courtship songs in reproductive isolation among populations of green lacewings of the genus Chrysoperla (Neuroptera: Chrysopidae). Evolution. 46(1):31–42. doi:10.1111/j.1558-5646.1992.tb01982.x.

West-Eberhard MJ. 2014. Darwin’s forgotten idea: the social essence of sexual selection. Neurosci Biobehav Rev. 46 Pt 4:501–508. doi:10.1016/j.neubiorev.2014.06.015.

Wicker-Thomas C, Chertemps T. 2010. Molecular biology and genetics of hydrocarbon production. Insect Hydrocarb Biol Biochem Chem Ecol.:53–74.

Wicker-Thomas C, Garrido D, Bontonou G, Napal L, Mazuras N, Denis B, Rubin T, Parvy J-P, Montagne J. 2015. Flexible origin of hydrocarbon/pheromone precursors in Drosophila melanogaster. J Lipid Res. 56(11):2094–2101. 10.1194/jlr.M060368.

Wicker-Thomas C, Guenachi I, Keita YF. 2009. Contribution of oenocytes and pheromones to courtship behaviour in Drosophila. BMC Biochem. 10:21.

Williams TM, Carroll SB. 2009. Genetic and molecular insights into the development and evolution of sexual dimorphism. Nat Rev Genet. 10(11):797–804. doi:10.1038/nrg2687.

Wu CI, Hollocher H, Begun DJ, Aquadro CF, Xu Y, Wu ML. 1995. Sexual isolation in Drosophila melanogaster: a possible case of incipient speciation. Proc Natl Acad Sci. 92(7):2519–2523. doi:10.1073/pnas.92.7.2519.

Yew JY, Chung H. 2015. Insect pheromones: An overview of function, form, and discovery. Prog Lipid Res. 59:88–105. doi:10.1016/j.plipres.2015.06.001.

Yoshimizu T, Akutsu J, Matsuo T. 2022. An Indirect Cost of Male-Male Aggression Arising from Female Response. Zoolog Sci. 39(6):514–520. doi:10.2108/zs210116.

Zwarts L, Versteven M, Callaerts P. 2012. Genetics and neurobiology of aggression in Drosophila. Fly (Austin). 6(1):35–48. doi:10.4161/fly.19249.

